# A Double Error Dynamic Asymptote Model of Associative Learning

**DOI:** 10.1101/210674

**Authors:** Niklas H. Kokkola, Esther Mondragón, Eduardo Alonso

## Abstract

In this paper a formal model of associative learning is presented which incorporates representational and computational mechanisms that, as a coherent corpus, empower it to make accurate predictions of a wide variety of phenomena that so far have eluded a unified account in learning theory. In particular, the Double Error Dynamic Asymptote (DDA) model introduces: 1) a fully-connected network architecture in which stimuli are represented as temporally clustered elements that associate to each other, so that elements of one cluster engender activity on other clusters, which naturally implements neutral stimuli associations and mediated learning; 2) a predictor error term within the traditional error correction rule (the double error), which reduces the rate of learning for expected predictors; 3) a revaluation associability rate that operates on the assumption that the outcome predictiveness is tracked over time so that prolonged uncertainty is learned, reducing the levels of attention to initially surprising outcomes; and critically 4) a biologically plausible variable asymptote, which encapsulates the principle of Hebbian learning, leading to stronger associations for similar levels of cluster activity. The outputs of a set of simulations of the DDA model are presented along with empirical results from the literature. Finally, the predictive scope of the model is discussed.

---

Associative learning aims at understanding the precise mechanisms by which humans and animals learn to relate events in their environment. Associative learning has been replicated across numerous species and procedures (Hall, 2002; Pearce & Bouton, 2001; Turkkan, 1989); its neural correlates have been extensively studied (Gomez *et al.*, 2001; Kobayashi & Poo, 2004; Marschner, Kalisch, Vervliet, Vansteenwegen, & Büchel, 2011; Panayi & Killcross, 2014; Roesch, Esber, Li, Daw, & Schoenbaum, 2012); it has proved to be a core learning mechanism in high-order cognitive processes such as judgment of causality and categorization (Shanks, 1995), and rule learning (Murphy, Mondragón, & Murphy, 2008); it underpins a good number of clinical models (Haselgrove & Hogarth, 2011; Schachtman & Reilly, 2011); and its evolutionary origins are beginning to be elucidated (Ginsburg & Jablonka, 2010). It is thus paramount that we develop comprehensive, accurate models of associative learning.

In classical conditioning, a fundamental pillar of associative learning, the repeated co-occurrence of two stimuli (e.g., an odor or tone), S1 and S2, is assumed to result in an association between their internal representations, which entails that the presence of S1 (the conditioned stimulus, CS, or ‘predictor’) will come to activate the internal representation of a S2 (the predicted stimulus or outcome from now on). When the outcome is a biologically relevant stimulus (unconditioned stimulus, US) able to elicit an unconditioned response (UR) learning results in the acquisition of a new pattern of behavior: the sole presence of the CS engenders a conditioned response (CR) similar to the UR. The response is assumed to express the strength of the association between the CS and the outcome (Pavlov, 1927), revealing that the outcome is anticipated or predicted by the CS.

In the following sections we proceed with a critical review of different models of classical conditioning structured around four major features in learning theory, namely, the learning rule, the elemental and configural bases of stimulus representation, attentional factors in CS processing, and, finally, the elusive nature of neutral and absent cue learning. We shall proceed with the description of the DDA model, which we claim provides a formal, coherent corpus for the understanding of classical conditioning. Next, we present a battery of simulations of relevant phenomena, and conclude with a discussion of the main contributions of the model.

DDA’s fully-connected network explicitly incorporates to the classical Pavlovian structure interactions between so called neutral stimuli, physically present or associatively retrieved, as well as the context in which they occur. As a result, the model accounts for phenomena that posit a problem for many models of classical conditioning, importantly, though not exclusively, learning about absent cues (mediated learning). Crucially, the DDA model provides a systematic interpretation of such effects in that they naturally emerge from the synergy of the model’s unique features, namely, a double error term and a dynamic asymptote, along with a variable attentional rate. This is in contrast to pre-existing models, which are conceived to address specific types of phenomena (e.g., Pearce, 1987) or defined on the basis of ad hoc rules which are capable to account for particular sets of results but that fail in explaining others which are sometimes contradictory in appearance (e.g., Holland, 1993; Dickinson & Burke, 1996). To our knowledge, there is no other model able of integrating all types of mediated learning phenomena, while accounting for a large number of other, in principle unrelated, phenomena, such as those derived from contextual effects on acquisition (latent inhibition) and extinction.

## The Learning Rule: Error Correction

The acquisition of a conditioned response, assumed to mirror the strength of the CS→US link, usually follows a negatively accelerating monotonically increasing curve over trials (but see Gallistel, Fairhurst, & Balsam, 2004; Glautier, 2013). In error correction models, the associative link’s rate of change is proportional to the discrepancy between the expectation and the presence of the outcome, i.e., the prediction error, which is minimized over training and results in the characteristic learning curve. This relation was initially mathematically formalized in linear operator models such as Hull’s early quantitative theory of learning (Hull, 1943) and stochastic theories of conditioning (Blough, 1975; Bush & Mosteller, 1955). A secular trend in modeling since then has been the expansion of the ontology of internal stimulus representations and learning processes governing the formation of associations between said representations. This increase in model complexity has expanded the quantity of phenomena that can be accounted for by models of learning (Alonso, Sahota, & Mondragón, 2014; Alonso & Schmajuk, 2012; Balkenius & Morén, 1998; Pearce & Bouton, 2001), and has been propelled and refined by experimental data acting as an arbiter of these models. For instance, evidence for the hypothesis that cues compete with one another for associative strength necessitated advancements from linear operator error terms. The phenomenon of blocking (Amundson & Miller, 2008; Kamin, 1968, 1969; Kohler & Ayres, 1979) showed that when the acquisition training for a cue A is followed by acquisition training with a compound AB, the novel cue B acquires next to no conditioning. Thus, it seems as if the associative link formed by cue A to the outcome prevents the formation of an equivalent link forming between B and the outcome. This result indicates that the processing of the outcome plays a significant role in determining the maximal amount of learning supported between itself and the CS. Such cue competition during learning is formalized most prominently by the Rescorla-Wagner (RW) model with its ‘global’ prediction error term (Rescorla & Wagner, 1972), which incorporates within it a sum of the values of all associative links to the US. This is an innovation upon other linear operator error terms, as for instance the Hull error term in contrast only incorporates the associative strength of an individual CS. Hence, learning between a CS and US in the RW model is driven by the total discrepancy between the US presence and the expectation elicited for it by all cues, as seen in Equation 1.

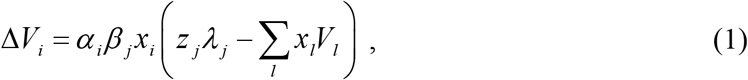

where Δ*V*_*i*_ is the change in associative strength, *α*_*i*_ and *β*_*j*_ are respectively the CS and US intensities, *x*_*i*_ is the activity of the CS being updated, *x*_*l*_ and *z*_*j*_ the activity of any present CS and the US, respectively, and *λ*_*j*_ the US asymptote determining the maximal supported learning. Finally, the summed term is the total extant learning toward the US, calculated as the dot product of the CS activities times their associative strength *V*_*l*_ toward the US. The RW model thus not only accounts for empirical data, but its formulation implied the existence of learning effects not predicted by earlier models or assumed to exist by preexisting theory, as can be tested in a wide range of simulations (Alonso, Mondragón, & Fernández, 2012; Mondragón, Alonso, Fernández, & Gray, 2013). Primary among these predictions is that if two CSs are independently conditioned to an asymptotic level, then their reinforcement in compound should lead to a decline in their associative strength. This follows from the error term supporting a maximal level of conditioning for all present stimuli on a trial, which is different from the maximal level supported for cues presented in isolation. Such a phenomenon of overprediction has been confirmed (Rescorla, 1970), along with other predictions such as super-conditioning (Rescorla, 1971a), thus strengthening the validity of the model as well as other models relying on a summed error term. Though the RW model has been successful in this regard (Miller, Barnet, & Grahame, 1995), a summed error term is not the only means of predicting cue competition results. Models relying on CS-US specific ‘local’, non-competitive error terms are capable of accounting for some of these effects by assuming other processes of competition, for instance on attentional competition between the predictors, as postulated by the model of Mackintosh (1975). Alternatively, the comparator hypothesis model (Miller & Matzel, 1988; Miller & Witnauer, 2016) explains cue competition results as a retrieval effect. It assumes that when a previously reinforced CS is presented (the target), it re-activates both representations of other CSs paired with the US and the US representation itself. The response elicited by the target CS is then proportional to the degree to which it predicts the US relative to the associative strength of the comparator CSs. Hence cue competition arises from the interference of other CS-US associations upon the target-US association.

The realization of real-time learning models extending the trial-based RW model (and other error correction models) by making predictions for learning within a trial allows for the modelling of time dependent aspects of learning and temporal relations between stimuli not accounted for by trial-level models. The SB model (Sutton & Barto, 1981), extended the delta rule used in RW by postulating that the variation in the inputs to a node representing a US was the driver of learning. Hence, both changes in the US sensory input (reinforcement) and changes in the CS contribution to this node (temporal difference) influenced the direction of learning. In addition to phenomena predicted by RW, SB accounts for inter-stimulus interval (ISI) effects and an anticipatory CR build-up (Balkenius & Morén, 1998). It nevertheless produces a few erroneous predictions, such as that a co-occurring CS and US should become highly inhibitory towards one another. Flaws of the SB model are rectified in the Temporal Difference (TD) model (Sutton & Barto, 1987) through the variations in the CS signal to the US node being dissociated from the US signal itself, and through the introduction of a time-discount factor that modulates the contribution of future predictions. The TD model uniquely predicts that learning is being driven by the need of the animal to minimize a time-discounted aggregate expectation of future reinforcement. At each time-step, the components of a given CS produce a prediction for the moment-by-moment change in US activation at the next time-step, termed the temporal difference. The difference between this prediction and the actual US activation level results in an error term (like that in the RW model), the prediction error, for that time-step. Equation 2 is used for calculating this error.

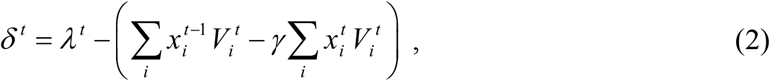

where *λ*^*t*^ is the US activation at time *t*, and the term 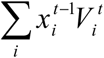 denotes the prediction for the US produced through the summation of associative activations of the US (using current associative links, but CS activation levels from the previous time-step), and 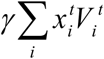 is the equivalent for the current time-step. The current time-step is multiplied by the mentioned discount parameter *γ* to reflect that the future is always slightly uncertain and therefore more recent stimulus activation carries more weight in calculating errors. The difference between these two prediction terms produces an estimate of what the activation of the US is during the current time-step. The discrepancy between the TD error and the actual US presence produces the overall prediction error for the US. As a real-time rendition of the US-processing error correction learning found in RW, TD can account for the same phenomena as the former on a trial level. In terms of predictive power, the instantiation of the TD model in simulators (Mondragón, Gray, & Alonso, 2013) has produced accounts of phenomena difficult to analyze a priori. For instance, higher-order conditioning, wherein reinforcement is observed in the absence of a US through the reinforcement effects of the temporal difference term itself. It accounts for temporal primacy effects – e.g., Kehoe, Schreurs, & Graham, (1987), wherein the presentation of a reinforced serial-compound of the form CS_A_→CS_B_→US results in a deficit in CR acquisition to CS_B_. It additionally reproduces the retardation effects of ISIs on the level of asymptotic learning (and by extension, CR over time) in an emergent fashion (Balkenius & Morén, 1998) through the aforesaid higher-order conditioning, which in itself results from the temporal-difference term back-propagating to earlier time-points. As the core of the TD model is its distinct error term, variations of TD with differing stimulus representations have been proposed. A complete serial compound model (Moore, Choi, & Brunzell, 1998) postulated that a CS is represented in time by a series of separate units, each of which becomes active following the previous. This extension produced a response curve in time with closer correspondence to empirical fact. Work in timing as well as hippocampal data inspired the micro-stimulus representation of TD (Ludvig, Sutton, Verbeek, & Kehoe, 2009), which instead assumes that the presence of a CS produces a cascade of units to become active in a bell-shaped form, with units that peak later having a correspondingly lower amplitude and higher variance. The advantage of such a stimulus representation is that it reproduces both differential responding early and late during a CS presentation, allows for effective trace conditioning due to the persistence of later micro-stimuli, and produces generalization of learning in time due to the significant activation overlap of said micro-stimuli. Extensions of the classical TD formulation have also been motivated by its lack of compound stimuli and configurations in its stimulus representation. Formulations of TD such as SSCC TD (Mondragón, Gray, Alonso, Bonardi, & Jennings, 2014) offer rules for forming such compounds both when stimuli overlap temporally, and when they are presented in succession. Steady evidence has mounted that predictions errors akin to the TD error are correlated to mid-brain dopamine function (Ludvig, Bellemare, & Pearson, 2011; Montague, Dayan, & Sejnowski, 1996; Niv, 2009; Niv, Edlund, Dayan, & O’Doherty, 2012; Schultz, 2004, 2006, 2010; Schultz, Dayan, & Montague, 1997).

In conclusion, although alternative approaches to summed error correction such as the comparator hypothesis model exist and are capable of accounting for crucial cue competition effects, the fit of the RW and TD error terms to empirical data correlating prediction errors and dopamine signaling, as well as a consideration for parsimony, motivate our use of the summed error correction rule in the DDA model we are introducing.

## The Nature of the Stimulus Representation: Elemental vs. Configural Models

Causal relations in the world are mostly more complex than the linear CS-US relationship encountered in a standard acquisition protocol, being more appropriately described by non-linear conditional probabilities. Learning models therefore must in some manner involve processes of approximating these conditional relations, such as those seen in non-linear discrimination learning. Exactly how such causal connections are approximated is highly dependent upon how the stimuli themselves are represented by a model. Two benchmarks of non-linear discriminative performance of a model have been negative patterning and biconditional discriminations. In negative patterning (NP) individual presentations of two cues A and B are followed by an outcome (a reinforcer), A+, B+, whereas compound presentations of the same cues are not, AB-. Its difficulty and hence importance lies in the simple breakdown of linearity on the compound trials. That is, the animal must learn to withhold responding on trials when two cues, which individually predict the outcome, are presented. Biconditional discriminations involve yet more complex nonlinearity. Four cues are presented in pairs, with each individual cue being presented in both a reinforced and nonreinforced compound (AB+, CD+, AC-, BD-). Therefore, simple summation of the individual cues values offers no information for solving the discrimination.

The Rescorla-Wagner (RW) model, as an elemental model, kept with the assumption of Spence (1936, 1937), Konorski (1948), and Estes (1950) that sets of elements constituting the attributes of individual stimuli enter into associations with the US directly (Harris, 2006; Wagner, 2008). As a result, in its original formulation, RW was unable to account for both negative patterning (Rescorla, Grau, & Durlach, 1985) and biconditional discriminations (Rescorla, 1972; Saavedra, 1975). The model assumes that if each individual stimulus is presented in opposite contingencies, this amounts to partial reinforcement of the cue and hence the discrimination will not be solved. In the case of NP, the model would predict greater responding on the compound trials than individual trials, due to the summation of the associative strengths of A and B. That is, the model preserves the linearity of summation by positing that the responding elicited by a given configuration of cues is directly proportional to the sum of the individual associative strengths of the constituent stimuli. The addition of elements common to multiple stimuli to RW allowed for the discrimination to be solved as the elements common to both A and B acquire superconditioning, super-asymptotic learning (Rescorla, 1971a), while the unique elements of both stimuli become inhibitory toward the outcome. Hence simply by assuming some similarity between CS representations it is possible to account for some non-linear discriminations within an elemental framework. In the case of more complicated discriminations such as bi- and tri-conditional discriminations, compound similarity may not be a sufficiently powerful mechanism. The addition of configural elements to the model, which become active during the presentation of a specific compound, allows the RW model to avoid this issue (Wagner & Rescorla, 1972). That is, it is assumed that the compound is represented by the animal as more than the sum of its parts. Hence NP is produced in a contrary fashion as compared to when common elements are involved, as the configural elements produce inhibitory links toward the outcome, while the unique elements form excitatory associations. While this configural element leads to the model predicting deviations from summation linearity in non-linear discrimination training, it nevertheless preserves linearity in the absence of such conditioning. As such, it was unable to account for evidence of linearity of summation being broken (Razran, 1939), that is for configurations of stimuli eliciting more or less response than expected by summing the response strengths elicited by individual presentations of the constituent stimuli. For instance, the observation that the presentation of a stimulus compound, after reinforcing the two stimuli separately, does not always produce more responding (Pearce, George, & Aydin, 2002) contradicted the RW model. Such evidence prompted the conception of the replaced elements model (REM) (Wagner, 2003), which maneuvered around this difficulty by a process whereby some elements of a stimulus are added (configural elements), and some are removed (unique elements), when the cue is presented in a compound. These added and replaced elements are consistent throughout presentations. The added elements correspond to elements representing context-dependent features of the stimulus, while the replaced elements are assumed to represent features of the cue that are uniquely present when the cue is presented alone. The model postulates a replacement parameter *r*, which determines what proportion of elements are replaced in this manner. Thus when *r* = 0, the model is equivalent to the RW model, while *r* = 1 produces a purely configural model (Schultheis, Thorwart, & Lachnit, 2008). The intermediate values allow the model to postulate that summation is not absolute. For instance, *r* = 0.5 leads to the prediction that no summation is observed, that is the compound of two CSs is no more predictive of the outcome than an individual CS. Thus, by assuming that different modalities of stimuli undergo different replacement rates depending on their similarity, consistent with psychophysical theory (Glautier, Redhead, Thorwart, & Lachnit, 2010), the model can explain why summation seems to depend on the types of stimuli used (Wagner, 2003).

Though the configural elements representation of RW and the REM model account for some aspects of deviance from perfect summation, they are less capable of dealing with other observed effects. Both models assume that a redundant cue, X, will facilitate the learning of a discrimination, yet the opposite is often observed (Pearce & Redhead, 1993). A further difficulty is that an underlying implication of most elemental models is the potential complete reversibility of an association between a stimulus and the outcome should the previously learned contingency be reversed (e.g., reinforcement followed by nonreinforcement). However, experiments displaying retroactive interference in feature negative discriminations, in which B+ trials failed to impair previous A+, AB-training, contradict this assumption of complete associative reversibility (Wilson & Pearce, 1992). Both the RW model and the REM extension have difficulty reproducing this effect. The former, due to its linear summation, predicts complete interference of B+ training on the A+, AB-training. That is, it expects that after B+ training the AB-compound will display less complete suppression of responding. The latter, though it in principle can reproduce the effect, is forced to postulate a very high replacement rate of B elements. However retroactive interference in feature negative discriminations has been produced by stimuli from different modalities (Pearce & Wilson, 1991), which seems to require a low replacement rate of elements. Concomitant to this procedural manipulation, however, lies the notion of stimulus modulation of the association (occasion setting), according to which a serial feature negative discrimination would endorse the feature with hierarchical modulatory properties over the excitatory association (Holland, 1985) or would increase the threshold for the US activation (Rescorla, 1985). Failure to impair the previous discrimination levels by subsequent feature reinforced training has been taken as evidence that negative modulators capabilities are relatively independent of their association to the outcome. In other words, the association between the feature and the outcome could be reversed without affecting the feature’s ability to suppress behavior.

An alternative account to the elemental approach is supplied by Pearce (1987), which in contrast presumes that nodes representing individual stimuli connect to an additional configural node representing their aggregation. This node in return forms associations with the outcome. Its configural stimulus representation, wherein the presentation of a compound activates a configural node representing it, produces many non-linear discriminations simply through learning between a configural node and the outcome not directly interfering with other configurations. Non-linear discriminations are often difficult for animals to learn. Pearce’s model accounts for this difficulty by postulating that responding to a configuration of stimuli is affected by its similarity to other configurations (as measured by a similarity index). Hence, it anticipates that the difficulty of a non-linear discrimination is directly proportional to the similarity of the constituent compounds. In the case of biconditional discriminations, the complexity of the discrimination arises from each compound being similar to another compound undergoing opposite reinforcement (e.g., AB+ and AC-). The model naturally accounts for the fact that summation between stimuli is not always being observed. As a result, it faces a challenge explaining evidence that summation does sometimes occur. Further, the model predicts a symmetrical deficit in responding (generalization decrement) when a stimulus is added in compound to a previously conditioned stimulus (external inhibition) or when a stimulus is removed from the training compound (overshadowing). Yet, it has been confirmed that overshadowing produces a larger deficit in responding than external inhibition (e. g., Brandon, Vogel, & Wagner, 2000). Finally, evidence has accumulated that familiarity with the constituent stimuli of a discrimination can facilitate the learning of the discrimination (Hall, 1991; Mondragón & Hall, 2002). This phenomenon of perceptual learning (PL) seems to suggest that the animal’s perception of the similarity of two stimuli is learned rather than given a priori as through Pearce model’s similarity index. Hence, some form of attentional or learning-based process seems to be necessary to dichotomize unique and redundant elements of stimuli when they are preexposed together, such as elements shared by the preexposed stimuli losing associability.

Configural models have been further critiqued from a theoretical perspective. In terms of parsimony, they require representations of both individual stimuli as well as configurations, thus making them more complex than purely elemental models – though Ghirlanda (2015) has demonstrated that a mapping between elemental and configural representations can be constructed analytically. Furthermore, they seem to take for granted representational information that is usually postulated by purely associative means, namely that a combination of stimuli co-occurred. Finally, they require some limiting process on the quantity of configurations that can form to avoid an infinite generation of different configural representations.

Harris (2006) and Harris and Livesey (2010) introduced a purely elemental model utilizing a unique attentional process to avoid the aforesaid problems of the elemental approach. In the model a finite ‘attentional buffer’, which makes the activation of elements stronger and more persistent (and therefore contribute more towards responding), is proposed. Entry into this buffer is regulated by the change in activation of a given element when it is presented or predicted and is proportional to the salience of that element (with the saliences of elements of a stimulus assumed to be normally distributed). That is, elements that undergo a strong increase in their activation can enter the attentional buffer. If the buffer is at full capacity, elements with larger changes in activation displace elements with smaller changes in activation. Further, the extent of reinforcement by a US is proportional to the number of US elements that are pushed into the buffer. However, in general, the quantity of CS elements in the buffer does not influence the direction of learning. The buffer simply speeds up learning as the activation of an element functions as a de facto salience. An important implication is that when a stimulus compound is presented, only the most salient elements can enter the buffer. Hence the model can account for partial summation. This partial summation, together with an assumption of common elements equips the model with the capability of predicting various non-linear discriminations. It additionally predicts that these discriminations are solved more slowly when a redundant cue is present, as the redundant cue leaves less capacity in the attentional buffer for relevant cues. The facilitation of discriminative learning by preexposure, i.e., perceptual learning, arises through a process of common elements losing associability through self-prediction hindering their entry into the buffer. Crucially, the model can explain both the lack of retroactive interference in feature negative discriminations (Wilson & Pearce, 1992), as well as the discrepancy between external inhibition and overshadowing (Brandon, Vogel & Wagner, 2000). In the case of the former, it postulates that during the B+ presentations, the elements of B that entered the attentional buffer on AB-trials are more inhibitory than the ones which did not. Thus, the elements that did not enter the buffer acquire more excitation than the ones that did. Consequently, when the AB compound is once again presented, much of the increase in B’s associative strength will not be manifested due to the most excitatory elements not entering the buffer. In the latter case, the model explains that in the case of external inhibition, the novel stimulus pushes elements of the compound out of the buffer. However, these elements nevertheless remain active at a lesser level, thus still contributing towards responding. When a cue is removed as in an overshadowing test, its elements are completely inactive, and hence the asymmetry between the phenomena is accounted for. Thus, the Harris model shows that many of the apparent pitfalls of elemental models can be avoided using a richer stimulus representation and a process of selective attention; though the exact nature and substratum of the attentional buffer remains to be validated by empirical data. Further, many of the unique predictions of the model could be explained also for instance by the simpler assumption of the salience of a cue directly influencing the strength of its activation.

In conclusion, both elemental and configural approaches to stimulus representations have distinct advantages in explaining non-linear discrimination learning. Configural models can offer a solution to how animals learn complex discriminations, generalization between them, as well as for partial summation. Elemental models must postulate additional learning and representational mechanisms to account for many of these effects. For instance, configural elements or the attentional buffer of Harris significantly extend the reach of elemental analysis. They nevertheless have the advantage of maintaining representational simplicity as well as encapsulating more information in terms of pure associative learning. That is, in many formulations they avoid the problem of how configural representations emerge in the first place. As such, we have favored an elemental approach for the current model.

## Attentional Factors: CS Processing

As the Harris model assumes that any excitatory link from one stimulus to another induces activation in the recipient stimulus, it can explain various preexposure and habituation phenomena as well. In effect, the associability of a cue, all else being equal, is therefore directly proportional to its novelty. In addition, the model’s prediction that the effective associability of stimuli decreases in relation to how many further stimuli are active has been experimentally validated (Lachnit, Schultheis, König, Üngör, & Melchers, 2008). This sets it in opposition to other paradigms of selective attention. For instance, the Mackintosh model assumes that animals attend to cues that are relatively better predictors of outcomes than other cues (Mackintosh, 1975). It models attention to a cue, *α*_*i*_, as changing in proportion to the relative predictiveness of the cue (in relation to other cues) for the outcome. The implication is that selective attention is learned, retained for future learning, and presumably aids in reducing proactive interference between stimuli, thereby speeding up learning as discussed in (Kruschke, 2011). Cue competition effects are thereby explained by purely attentional means and thus the model only requires the linear operator delta rule familiar from Hull’s model. The Mackintosh model’s unique assumption that the selective attention paid to a cue has direct reinforcing effects has allowed it to predict phenomena that pose a problem for other models. For example, it can explain unblocking, wherein the surprising partial omission of reinforcement during a blocking treatment attenuates the blocking of a cue (Dickinson, Hall, & Mackintosh, 1976). It thus avoids the prediction of overprediction pushing the blocked cue towards becoming inhibitory, which the RW model predicts in some circumstances for this treatment. An alternative explanation to that offered by the Mackintosh model for this effect is however that differential reinforcement is represented by the animal as different reinforcers. As such, this result can be accounted for by the RW model. The Mackintosh model also accounts for the learned irrelevance effect (Bonardi & Hall, 1996; Mackintosh, 1973), whereby a CS uncorrelated with US presentations shows poorer subsequent acquisition, due to the best relative predictor accruing the most associability, and hence conditioning more quickly than competing cues. A well-known difficulty faced by the model is however the phenomenon of Hall-Pearce negative transfer (Hall & Pearce, 1979), wherein reinforcing a CS with a weak US, thus making it a better predictor, hinders subsequent conditioning between the same CS and a stronger US. In this case, it erroneously predicts greater excitatory learning between the previous best predictor of the outcome and the outcome. The model also is unable to account for superconditioning (Rescorla, 1971a, 2004), wherein presenting a conditioned inhibitor of an outcome together with a novel cue leads to stronger excitatory conditioning to the novel cue than if it were conditioned individually. As the Mackintosh model does not assume a competitive error term, it cannot account for this effect. Superconditioning is however predicted by Le Pelley’s extension of the Mackintosh model (Le Pelley, 2004), owing to the use of a combined error term.

The Hall-Pearce negative transfer effect is easily accounted for by the Pearce and Hall (PH) model (Pearce & Hall, 1980), which postulated that attention rises when the outcome is uncertain. That is, the associability of a CS rises in accordance to the general uncertainty of the outcome instead of tracking the relative predictiveness of the cue. By implication, in the Hall-Pearce negative transfer effect, prior training with a weaker US produces a decline in associability which delays subsequent learning with the stronger US. In contrast to the Mackintosh model, the Pearce and Hall model has trouble explaining learned irrelevance, as the predictor in a learned irrelevance procedure will accrue more associability due to their lack of correlation with the occurrence of an outcome. In the context of a standard acquisition and extinction protocol the PH model predicts a sudden increase in associability when the contingency is changed (i.e., the beginning of acquisition or extinction), with a gradual decline thereafter. Further, a great success of the model has been predicting latent inhibition (LI) (Channell & Hall, 1983), wherein preexposure of a CS attenuates subsequent acquisition training with the same CS. It predicts this effect as arising from the preexposure leading to a decline in the associability of the CS, which subsequently rises again during acquisition training. Empirical evidence exists for the hypothesis that prediction errors increase the associability of cues, however a confounding factor is the difficulty of disentangling increases in associability (speed of learning) from direct reinforcement (extent of learning) produced by prediction errors (Holland & Schiffino, 2016). Uncovering and formalizing the precise nature of modulating factors of attention, and their relation to the dopamine system (Ahveninen *et al.*, 2000; Nieoullon, 2002), is hence critical in determining the plausibility of the associabilities of the Mackintosh and PH models. It has been noted that the attentional mechanisms underlying these two models are not necessarily in opposition. The existence of evidence supporting each model has fostered the proposal of dual-factor attentional models. One such model is the Le Pelley model (Le Pelley, 2004), which combines both the associability of the Mackintosh model and of the PH model (with the latter given more influence). According to Le Pelley’s model, both the general uncertainty of the outcome, as well as the relative predictiveness of a cue determine the overall associability of the cue. As such, the Le Pelley model can explain phenomena that have proven difficult for either model in isolation. It predicts, for instance, Hall-Pearce negative transfer as well as the learned irrelevance effect simultaneously. It however introduces further complexity, as the two rules can often cancel the influence of one another. Thus, it is also difficult to isolate their respective effects empirically.

It is worth noticing that the PH attention rule, when instantiated in a real-time model, can produce an increase in associability for the best predictor of the outcome if said predictor produces a prediction error for the US before the US onset. That is, the rule produces more complicated emergent behavior when instantiated in a real-time model. Further, it has been found that even trial-level models based on the PH attention rule can produce Mackintosh-like effects under certain conditions (Le Pelley, Haselgrove, & Esber, 2012).

The assumption that novelty of cues is crucial to how learning unfolds, pioneered by the Mackintosh and PH models, was expanded upon by the SLGK model (Kutlu & Schmajuk, 2012). It postulates that the novelty of every stimulus affects its speed of learning towards other stimuli. Further, this novelty is modelled through stimuli learning to predict each other through associative links (expectancy). Therefore, not only reinforced, but also non-reinforced (i.e., ‘neutral’ or ‘silent’) learning is incorporated. The model is configural in the sense that it postulates a layer of units intermediate between the sensory units and the US unit. These configural units have non-modifiable random incoming connections from all CS representations, and their connection to the outcome is adjusted through a delta rule. Their associability or associative rate is taken to be very low initially, however if the US expectancy remains high during training (i.e., the animal is unable to learn a given contingency), then the associability of the configural units rises. It hence accounts for the difficulty of animals to solve non-linear discriminations. This mechanism of detecting, through a persistent outcome error, when a purely linear approach has failed offers a robust approach for introducing non-linearity into models of conditioning while pre-empting a combinatorial explosion. It can be utilized to change the associability of other representational elements besides configural cues (e.g., common elements) to reflect when the animal cannot solve a learning problem.

Learning between non-reinforcing cues, i.e., ‘silent learning’ incorporated by the model endows it with the capacity to account for various preexposure effects. For example, it explains the attenuated positively accelerating response curve of a latent inhibited stimulus during subsequent acquisition. This response curve further approaches the standard learning asymptote after a sufficient number of reinforced trials (Lubow, 1965). In this regard, it shares similarities with two other models that postulate both silent learning and novelty-based associability, SOP (Wagner, 1981) and the McLaren and Mackintosh (2000) models. In all three models, latent inhibition emerges from the preexposed CS losing novelty by being predicted by the context and itself (unitization). This loss of novelty slows down subsequent acquisition training in a way different from that proposed by the aforementioned PH model and its Hall-Rodriguez extension (Hall & Rodriguez, 2010). These predict that the repeated preexposure of a cue reduces the attention paid to it, thus decreasing its associability. Hall and Rodriguez (2010) further predicts the formation of a CS→noUS link, which thereafter interferes with the response elicited by the CS→US link formed in the subsequent acquisition training. This latter mechanism bears a similarity to the explanation offered by the comparator model (Miller & Matzel, 1988), which also stresses processing during retrieval, although in this case, unlike in Hall and Rodriguez’s, the effect is specific to a given US. Nevertheless, the neutral-learning based account of SLGK, SOP and McLaren-Mackintosh seems to hold a categorical advantage in explaining latent inhibition, as they do not need to assume such a noUS representation or the existence of an extant associative link to it. Hence, they avoid predicting inhibitory properties of the preexposed CS towards the US, which have not been found in summation tests (Rescorla, 1971b). Additionally, assuming a noUS representation by implication would lead to generalization of inhibition to another outcome. Evidence seems to contradict such an assumption, as learning with a reinforcer has been demonstrated to be outcome specific, for instance in outcome devaluation experiments (Colwill & Motzkin, 1994).

In conclusion, models that assume that selective attention is acquired by the most predictive cue, such as the Mackintosh model, offer a robust account of effects such as unblocking and learned irrelevance. They however are unable to offer a convincing explanation of preexposure effects. Similarly, alternative explanations for unblocking and learned irrelevance can often be produced by error correction models. In the former case by assuming that differences in US presentations imply differences in the asymptote of learning; in the latter case by postulating that uncorrelated CS-US exposure results in a null correlation. In contrast, models that postulate a loss in associability as the outcome error declines (PH) or when a cue becomes less novel (SLGK, SOP, and McLaren-Mackintosh) are uniquely capable of explaining both the result and mechanisms behind the loss in associability occurring during preexposure. The latter models can also predict the context specificity of LI observed when preexposure is conducted in a context different from that of conditioning. Moreover, as above-mentioned, when instantiated in a real-time model, the uncertainty-driven variable associability of the PH model could produce emergent Mackintosh-like effects through the best predictor of an outcome yielding a larger prediction error before the onset of the outcome, thus increasing its own associability towards the outcome. We have hence chosen to incorporate a similar mechanism, with important changes involving revaluation of persistent uncertainty, to account for learned attentional bias in the DDA model.

## Neutral and Absent-Cue Learning

If an animal is learning the contingencies governing events, does it revaluate the presumed most probable contingency when it is contradicted by subsequent learning? If so, how? Since it is assumed that prior learning is stored as associations, this revaluation by implication involves the modification of previously formed associations. Behavioral data exists, which demonstrates that a present cue can retrieve the representation of another cue and thereby invoke revaluation of the latter’s previously formed associations towards a reinforcer (Dickinson, 1996; Holland, 1983; Holland & Forbes, 1982). In addition, neural data indicates that the left hippocampus is involved in mediated learning in humans (Jie, 2008) and the dorsolateral prefrontal cortex has been tied to violations of previously formed expectations (Corlett *et al.*, 2007). These so called mediated learning effects are of interest as they allow for an understanding of processes governing learning between representations of stimuli which are present and ones which are absent yet associatively invoked. Of cardinal significance are backward blocking (BB), unovershadowing/retrospective revaluation (UnOv) (Le Pelley & McLaren, 2001; Miller & Witnauer, 2016; Urushihara & Miller, 2010), and sensory preconditioning (SPC) (Brogden, 1939; Ward-Robinson & Hall, 1996). Explaining these effects with a simple associative learning rule has been elusive, as they seem to result from opposite learning directions between the cues, retrieved and present. Figure 1 depicts these seemingly contradictory mediated learning phenomena, succinctly describing their designs and the rules required to account for the observed results, to highlight the source of conflict in their interpretation.

**Figure 1:**
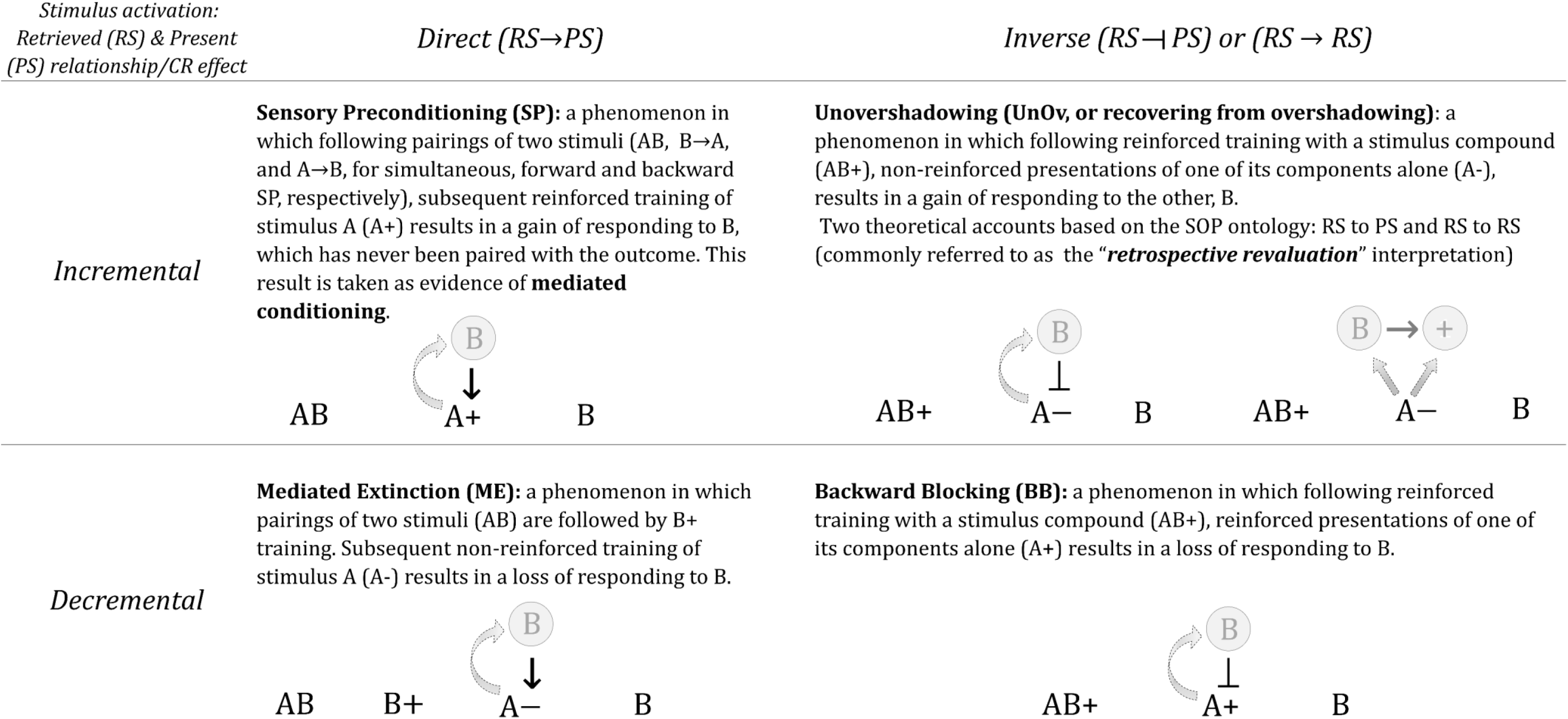
Critical mediated learning phenomena and putative learning rules required to account for Sensory Preconditioning (Mediated Conditioning, top-left); Mediated Extinction (bottom-left), Unovershadowing (top-right) and Backward Blocking (bottom-right). Relationship between the retrieved stimulus (RS, in grey circles) and the present stimulus (PS) activations is direct (→) or inverse (⊣) against the incremental or decremental effect on the CR. N.B. In some studies, mediated learning phenomena that involve an update of past learning experiences for non-present cues (UnOv and BB) and that that cannot, in principle, be explained by means of an associative chain, are referred collectively to as retrospective revaluation (right column).

For instance, in BB and UnOv designs, a compound of events, AB, is first associated to a motivational outcome. Once this association is established, presentations of one of its components alone, A, follows. While in a backward blocking procedure A is also reinforced, in a UnOv procedure, subsequent A training is not. Testing of the other element of the compound, B, afterwards, the BB treatment results in B losing associative strength compared to the first phase of learning, whereas in the UnOv treatment the excitatory strength of B increases. That is, strengthening the association between one of the elements of the compound and the outcome, results in a reduction in strength of the concomitant stimulus and the outcome, whereas lowering it rises the strength of its associated stimulus.

In a sensory preconditioning procedure, two non-reinforced stimuli, AB, are initially paired. In a second phase, A is reinforced. Subsequent test trials show that B, which has never been paired with the outcome, has nevertheless gained associative strength. Thus, an absent cue B undergoes a change in its associative strength as a consequence of the reinforcement treatment given to its associated cue. One possible mechanism for this phenomenon relies on the idea of mediated conditioning: during conditioning to A, an associative activation of stimulus B, which was linked to A during the initial compound training, is produced. This associatively activated representation of B acquires associative strength towards the present reinforcer. Of note is that learning between the absent cue, B, and the present outcome in SPC and BB proceeds in opposite directions. That is, while in SPC the absent cue acquires strength, in BB it loses strength.

Empirical research suggests that observing retrospective revaluation or mediated learning effects may depend on the degree of discriminability of the stimuli (Liljeholm & Balleine, 2009). Mediated learning will be preferentially obtained when the stimuli are poorly discriminated, whereas retrospective revaluation effects will be favored as generalization is reduced. There are other factors that are also known to influence the magnitude of UnOv: For example, the strength of the within-compound associations between the stimuli, so that the stronger the association, the higher the probability of observing a UnOv effect (Aitken, Larkin & Dickinson, 2001). In addition, UnOv is more readily observed with longer CS durations (Matzel, Schachtman & Miller, 1985). Moreover, a large number of revaluation trials seems to be required to produce a robust UnOv effect (Blaisdell, Gunther & Miller, 1999). Retrospective revaluation effects are also context dependent (Boddez, Baeyens, Hermans & Beckers, 2011). Finally, unovershadowing is more likely to emerge when the two cues hold differential saliences and the most salient member of the compound undergoes extinction (Liljeholm & Balleine, 2006).

Different models have been proposed to explain specific sets of mediated phenomena that can be ascribed to a single learning rule, either an increase or a decrease in strength in the same design conditions, but so far, no model can account for all of them. It would thus seem evident that a comprehensive model of learning should be able to predict these apparent contradictory results.

In the McLaren-Mackintosh model (McLaren & Mackintosh, 2000) stimuli are represented by sets of mutually overlapping elements in a connectionist network. Each node in the network has both an external (sensory) input, as well as a modifiable internal (associative) input, both of which contribute to its level of activity. The difference between these two inputs is considered the prediction error for that node, and thus determines the amount of associative learning from other nodes to it, as well as acting as a modulator of the associability from that node to other nodes. The internal input is precisely what allows for the retrieval of absent cues and therefore mediated conditioning in the model. McLaren and Mackintosh postulate that in sensory preconditioning, the preexposure of the compound AB results in the formation of bidirectional excitatory associations between the two CSs. Subsequently, when A is presented together with reinforcement, A associatively retrieves the representation of B. This associatively retrieved representation in return is capable of supporting learning towards the present outcome. Therefore, mediated conditioning is treated equivalently to learning between present stimuli. It is uncertain however whether the model can account for the revaluation effects of backward blocking and unovershadowing. For instance, in the case of backward blocking the delta rule employed would suggest that the retrieved cue B could potentially gain additional excitatory strength rather than losing it, as its prediction for the outcome would be lower than if it were directly present. That is, the delta error term would still support further excitatory learning, predicting an increase rather than the observed decrease in strength, thus the model would not be able in principle to account for BB, but would be able to predict UnOv.

Another partially successful model in accounting for mediated phenomena is SOP (Wagner, 1981). SOP postulates that inactive (I) stimulus representations are activated from long-term storage into short-term memory through either stimulus presentation or associative retrieval. The short-term memory system consists of an elemental network of processing units containing elements in primary (A1) and secondary, weaker (A2) states of activation for each stimulus center/unit, as seen in panel a) of Figure 2. Direct stimulus presentation results in the activation into A1 of a given proportion (p1) of its elements. Active elements (or associatively retrieved elements) decay in time into A2 according to a pd1 proportion. As with previous models such as RW, it is assumed that the total quantity of all stimulus A1 elements is limited. Therefore, increasing the number of A1 elements increases the speed at which these elements decay into the A2 state. Over time, elements in the A2 state of activation decay back into the inactive state. The model specifies that the direction of learning (excitatory or inhibitory) depends on the ratio of A1 to A2 elements of the outcome. Thus, a more novel outcome supports more excitatory learning, whereas an over-predicted outcome supports only inhibitory learning. These processes are summarized as learning rules in panel b) of Figure 2.

**Figure 2.**
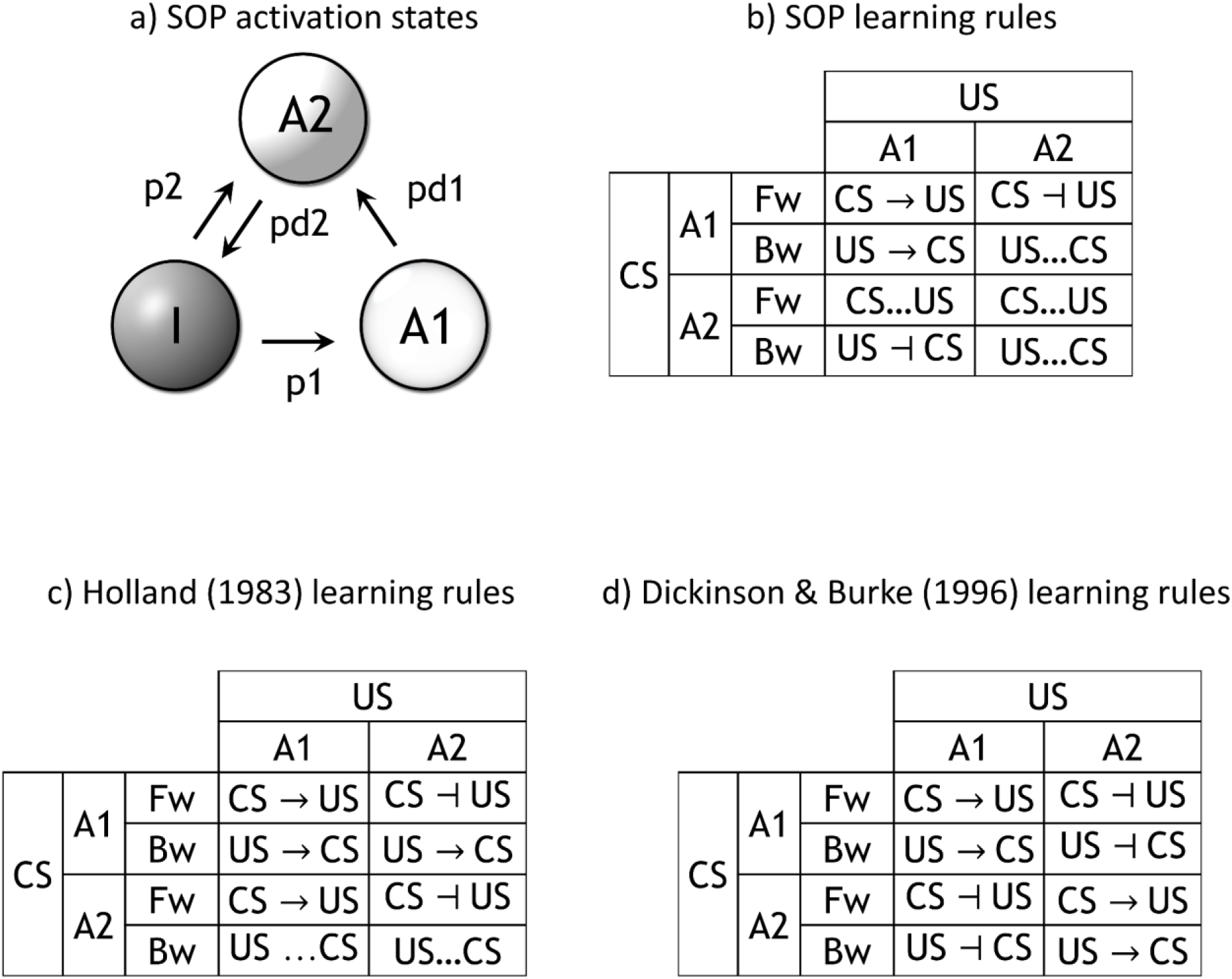
a) SOP activation states: I, A1, and A2 are respectively inactive, active, and associatively activated or decayed activation states; p1 and p2 are respectively rates by which inactive elements can be activated into A1 or A2 states; pd1 and pd2 are respectively rates at which A1 elements decay into the A2 states, and A2 elements decay into the inactive state. b) SOP learning rules. The arrow (→) describes the direction of an excitatory link, whereas (⊣) represents an inhibitory link vector, c) Holland SOP learning rules. d) Dickinson & Burke SOP learning rules.

SOP and its extensions (CSOP (Wagner & Brandon, 2001), and AESOP (Brandon, Vogel, & Wagner, 2003)) assume excitatory conditioning between fully active elements, and the formation of bidirectional inhibitory links between a present CS and an absent outcome (Figure 2, panel b), but do not offer further associative rules. Such rules are introduced by Holland (Holland, 1983), and Dickinson and Burke (Dickinson & Burke, 1996). The former (Figure 2, panel c) accounts for mediated conditioning and predicts that two stimuli in the A2 state undergo inhibitory learning. The latter (Figure 2, panel d) predicts backwards blocking (A2:A1 inhibitory) as well as mediated conditioning between two A2 stimuli. As is evident, these rules are in multiple instances (e.g., A2:CS A2:US and A2:CS A1:US rules) the reverse of one another. Indeed, experimental data has been collected to support each model (Dwyer, 1999), leaving the problem unresolved from the point of view of these two SOP extensions. That is, whether A2-A2 learning should be either excitatory or inhibitory is unclear. Hence, there exist conditions under which A2-A1 learning appears to lead to both excitation and inhibition.

Nonetheless, it is conceivable to explain mediated learning through mechanisms different from alterations in the associative learning rules. For instance, the comparator model produces retrospective revaluation through the competition of associative links during retrieval. That is, the B→US link is relatively larger than the A→US link after the latter has lost associative strength in the second phase. However, a formal analysis of various comparator models has cautioned that such revaluation effects might be harder to reproduce than originally conjectured (Ghirlanda & Ibadullayev, 2015, p. 18). Though the TD model does not presume learning between neutral stimuli, the Predictive Representations (Ludvig & Koop, 2008) extension of it accounts for mediated conditioning by proposing that present cues retrieve representations of stimuli with which they have been previously presented. These retrieved cues condition equivalently to present cues, save for having lower levels of activation. The formulation of the model however seems to preclude explaining backward blocking without further assumptions, as learning between a retrieved CS and a present outcome will tend to be excitatory. Hence the model conceptualizes mediated learning as a form of reasoning about causal relations between events, yet does not seem to capture reasoning occurring when prior learning is contradicted by subsequent learning in such a manner that previously formed links should be weakened.

A unique account of mediated learning is conceived of in the ‘replayed experience’ model of Ludvig, Mirian, Kehoe, and Sutton (2017). It in contrast postulates that the animal re-processes previous learning during the inter-trial interval (ITI), thereby consolidating contradictory learning contingencies. It can account for a wide variety of mediated learning effects including mediated conditioning and backward blocking in this manner. In a sense, it accounts for mediated learning by assuming that specific contingencies are intermixed as replay experiences in the mind of the animal, and for instance backward blocking is therefore explained in the same way as intermixed standard blocking. However, the model relies on the assumption that such post hoc revaluation indeed occurs during the ITI, and must assume that representations of different outcome-contingencies have already been learned by the animal for them to be replayed.

There are variations of the TD model and Rescorla-Wagner’s that can capture backward blocking and other revaluation phenomena based on the Kalman filtering algorithm. The Kalman Temporal Difference model (KTD) (Gershman, 2015) in particular, combines the Kalman-filter extension of Rescorla and Wagner’s model introduced in (Kruschke, 2008) and TD. The former, departs from Rescorla and Wagner’s (and most traditional models of classical conditioning) assumption of single reward predictions by accommodating distributions of beliefs that are updated according to Bayes’ rule; the latter renders an instantiation of Rescorla and Wagner’s in real-time and allows for predictions on long-term cumulative rewards. KTD can predict effects, such as latent inhibition, backward blocking and recovery from overshadowing, but, to our knowledge, the model does not offer a solution to other mediated learning phenomena which seem contradictory in nature such as mediated conditioning and mediated extinction.

The Latent Causes Theory (LCT) (Gershman & Niv, 2012; preliminary versions in Courville, Daw, & Touretzky, 2006; and Gershman, Blei, & Niv, 2010) is an alternative Bayesian model that diverges from the traditional assumption that associations are formed between conditioned and unconditioned stimuli, and postulates that animals infer models of the hidden (latent) structure of the environment over which beliefs are updated using Bayesian rules. Whereas LCT can predict a number of conditioning effects, early versions of the model failed to account for fundamental classical conditioning phenomena, such as blocking, and the authors themselves acknowledged that it was inadequate as a general model of classical conditioning (Gershman & Niv, 2012, p. 265). Integrating the rational theory of dimensional generalization (Navarro, Lee, Dry, & Schultz, 2008; Tenenbaum & Griffiths, 2001; Soto, Gershman & Niv, 2014) adapted LCT to account for several compounding effects. Nonetheless, their model fails to provide a mechanism that would explain such phenomena, instead it posits that learners infer what latent causes are more likely to be active from a pool of possible causes. LCT thus rely heavily on a potentially infinite capacity distribution of latent causes and may be susceptible to a combinatorial explosion of cause-stimuli configurations.

In summary, if mediated phenomena are to be explained purely through learning rules, what seems to be needed is therefore an operation that subsumes conflicting sets of rules. This supra-rule should reverse the direction of learning between present and absent cues in a principled manner depending on the prior reinforcement history. For instance, through a learning process, which can be justified as a form of approximate Bayesian inference. This is precisely the approach we follow in the DDA model.

To conclude, a plethora of models have been developed to account for wide varieties of classical conditioning phenomena. However, the sets of phenomena explained by models are quite distinct. Latent inhibition is predicted by diverse models, nevertheless the accounts given by the SLGK, McLaren-Mackintosh, and SOP models excel due to their ability to incorporate context modulation. Similarly, solving various non-linear discriminations, such as negative patterning, and modeling exposure effects on stimulus generalization, seem to necessitate a more elaborated stimulus representation than standard elemental and configural approaches. In terms of mediated learning, most of the models incorporating neutral stimulus associations can account for some but not all effects, with different models displaying different strengths.

The DDA model introduced next aims at accounting for all the highlighted classes of learning effects through a general framework of real-time error correction learning, instantiated as an elemental connectionist network. It integrates most prominently a unique second error term denoting the expectancy of an outcome predictor, a revaluation alpha which turns outcome uncertainty into a source of information, and critically, a dynamic asymptote of learning regulated by the similarity/discrepancy in the levels of activation of the elements of the association.

## THE MODEL

The DDA model introduced in this paper conceptualizes a formal, computational model of classical conditioning. It is instantiated as a connectionist network, with nodes representing clusters of elements belonging to stimuli. When two or more clusters of elements are activated, the elements of one node enter into association with other active clusters (including between non-US clusters, and clusters of the same stimulus). Unlike the element activity, which is binary (active, 1, or not, 0), the cluster’s activity is given by the mean number of its active elements. Elements in a cluster can be unique to a stimulus or shared between two stimuli. The model operates based on an error correction learning framework, thereby inheriting the properties of extant error correction models. Its associative learning rule includes a prediction error term for both the predicting and predicted stimuli, with both influencing the associability of a CS with an outcome. That is, a more novel predictor will undergo faster conditioning toward an outcome. Crucially, the asymptote of learning between any two clusters of elements measures the similarity of their activity. Hence, clusters of elements with similar activity patterns form stronger associative links. Finally, a process governing changes in attention-based associability is introduced and operates based on the time-discounted uncertainty in the occurrence of these stimuli being revaluated as a source of information.

The activity of elements is time-dependent, with groups of elements assembled into ‘temporal clusters’ allowing the model to predict differential responding early and late during a CS presentation. At a high level, learning occurs between motivationally neutral stimuli as well as between a neutral stimulus and a US, and the associations so learned modulate each other. Thus, the model accounts for preexposure effects such as latent inhibition. The predictor error term, which produces the so called ‘double error’ learning rule, along with the unique asymptote of learning used, endows the model with the capability of accounting for apparently contradictory mediated learning phenomena (e.g., BB and mediated conditioning). It does so by positing that the crucial factors influencing the direction of mediated learning are the strength of retrieval of the retrieved cue and the prior strength of the link between the retrieved cue and the outcome. Lastly, the attentional processes of the model contribute towards the model’s capability of predicting a wide range of phenomena including preexposure, non-linear discriminations, and contextual effects.

The model ontology is as follows: a stimulus, {A, B, C…, US, context}, can function either as a predictor or as an outcome (*p* ˅ *o*) and consists of a set of clusters {*i, j*,…}. Each cluster, in turn, is a collection of elements, such that each element *e* can be unique to a given cluster and stimulus or shared between clusters of two different stimuli. That is, *e* ∈ [(*i* ∈ A) ˅ ((*i* ∈ A) ˄ (*j* ∈ B))]. Elements are shared only between neutral stimuli. Although contexts and reinforcers could potentially share elements with other stimuli, their specific characteristics, the context ubiquitous distribution and its numerous and sensory diverse attributes, and the strong motivational value of the USs, would render their commonality contribution intangible, thus we have chosen to disregard it at this stage.

### Stimulus Representation and Activation

In the DDA model, a stimulus representation is constituted by a set of elements that are related to the physical attributes of the stimulus. These elements can be unique to the stimulus or shared with other stimuli. Shared elements (Figure 3) are sampled (in a binary fashion) whenever one of their ‘parent’ stimuli is active.

**Figure 3.**
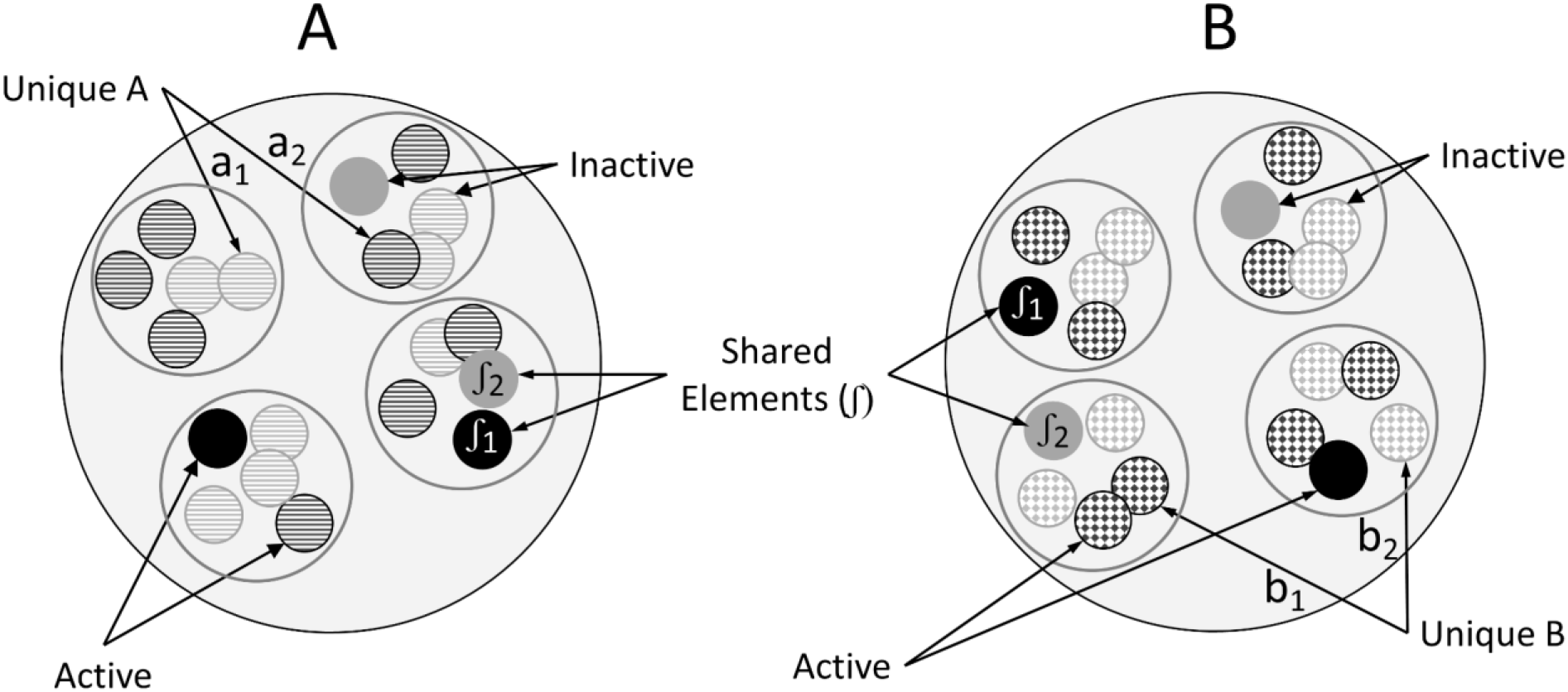
A stimulus consists of unique elements (e.g., a_1_, a_2_, represented with horizontally lined circles for A vs. b_1_, b_2_, spotted circles for B) and elements shared between two stimuli (∫_1_, ∫_2_, solid circles). The elements can be active (dark circles) or inactive (light circles).

Each element of a given temporal cluster, of a given stimulus, is capable of developing associative links to other temporal clusters (including temporal clusters within the same stimulus). Therefore, the model’s ontology presumes an elemental connectionist network structure (Figure 4) of the total stimulus representation, with links between elements of a given cluster and an ‘outcome cluster’ (i.e., elements modulate activity of clusters of elements). The DDA model postulates that elements learn to predict clusters of other elements, instead of forming direct element-element associations because the probability of sampling a given pair of elements (or the neural spiking behavior theorists attempt to approximate with them) is very low in comparison to the co-occurrence of an element and a cluster of elements. It is thus more plausible that a prediction would engender activation over a temporally stable cluster node than over a single element.^1^ In addition, learning is modelled in this form to represent the temporal structure of events which reconciles evidence for elements representing binary attributes of stimuli (that can hence be potentially shared in common between stimuli), together with evidence for associative activity operating in a continuous time interval. That is, learning is not modelled between elements due to their binary state of activity leading to a loss of the temporal information of the stimulus. Similarly, learning is not modelled between temporal clusters of stimuli, as this would disrupt the predictive temporal flow and render the contributions between elements unique to the cluster/stimulus and elements shared between stimuli indistinguishable. Finally, hypothesizing that temporal clusters as a whole could be shared between stimuli would impose unwarranted assumptions about stimulus commonality being dependent upon temporal dynamics.

**Figure 4.**
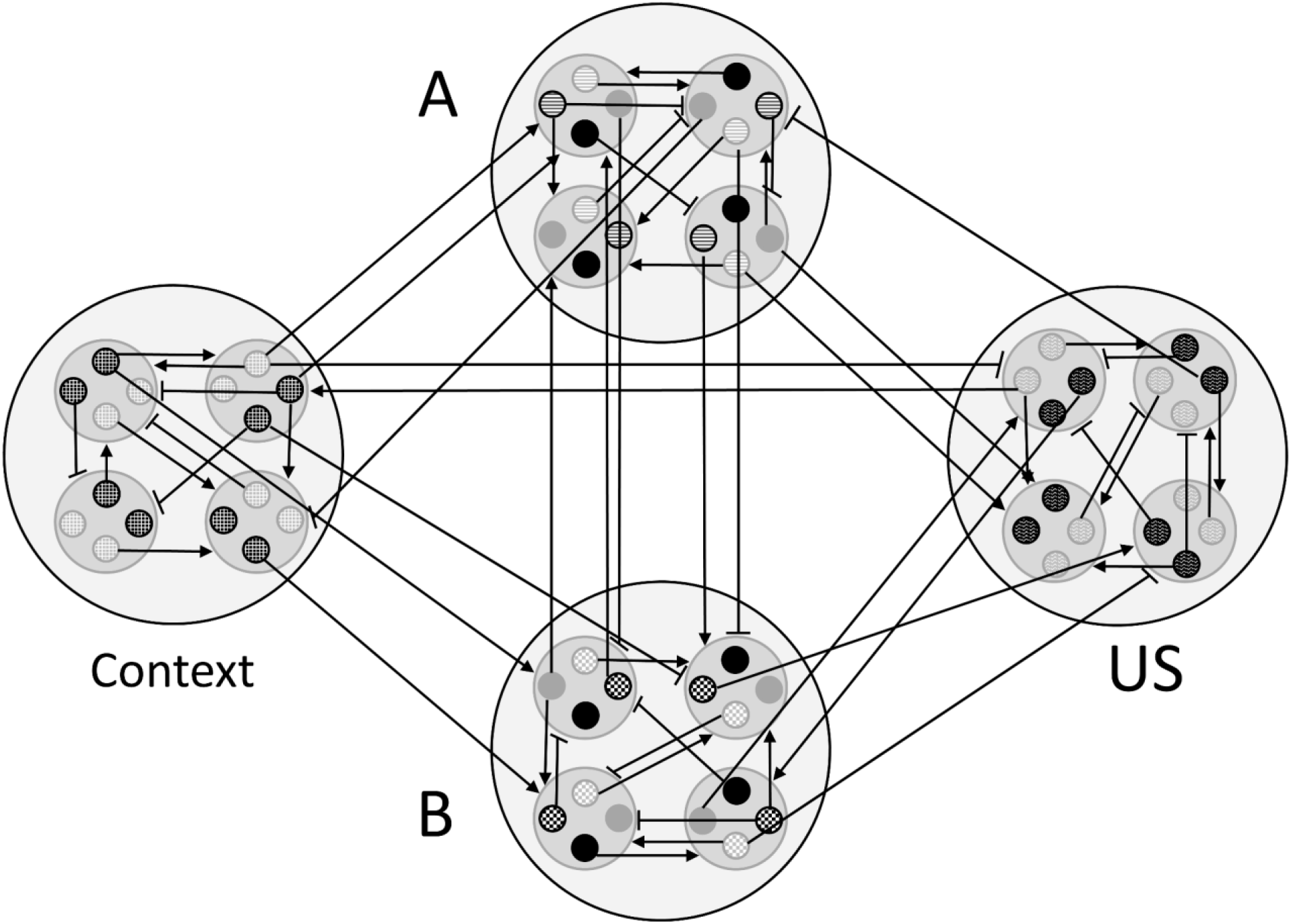
The elemental structure of the model as a connectionist network. Each stimulus (A, B, …, context, US) consists of a set of clusters of elements. Elements of a cluster can enter into association with any other cluster.

It is assumed that the stimulus representation varies through time, with some elements being differentially active early or late during the presentation of a stimulus. Hence, elements in a cluster are differentially active throughout the stimulus presentation: the probability of binary activation of each element in a cluster at a given time equals to that cluster’s temporal activation at that time (e.g., if the activation value of the cluster is 0.3 at time *t*, then each element has a 0.3 chance of being sampled at *t*). To clarify, the total activity of a temporal cluster is conceived of as resulting of a bottom up process of element sampling in which each element has a given probability of being sampled at a given moment in time. Elements that are correlated in their probability of being sampled will cluster temporally, and thus the total activity of the temporal cluster will correspond to the averaged binary activity of the elements that ‘belong’ to said cluster.

One temporal cluster is defined for each time-unit of the duration of the stimulus, with the maximal cluster’s activity occurring at time 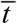. The probability with which an element within the cluster is activated follows an ‘approximately’ Gaussian distribution given by the cluster’s activation value at that time. Equation 3, denotes the direct activation distribution Φ for a cluster, *i*, acting as predictor *p* or outcome *o*, produced by physical stimulus presence.

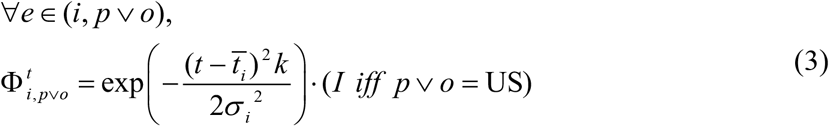

where 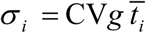 is the variance of the temporal cluster (i.e., temporal invariance is enforced, and CV, a coefficient of variation, and *g* are free parameters). We have assumed that 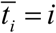, that is that the ‘*i*’ th temporal cluster peaks at time-point *t* = *i*. Additionally, *k* is a skew parameter that multiplies the enclosed term when *t* < *i*. If the cluster *i* belongs to a US, a scalar intensity value *I* multiplies the equation such that strong reinforcers are coded with high *I* values. The default value of *I* is set to 1.

The curve is not normalized by the usual 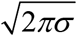 factor: we desire the peak of the curve to have a maximal value of 1, as this produces predictions interpretable as probabilities when combined with the asymptote used in the error term of the learning equation. For instance, with a 10 time-unit stimulus, the constituent temporal clusters of the stimulus follow the shape seen in Figure 5.

**Figure 5.**
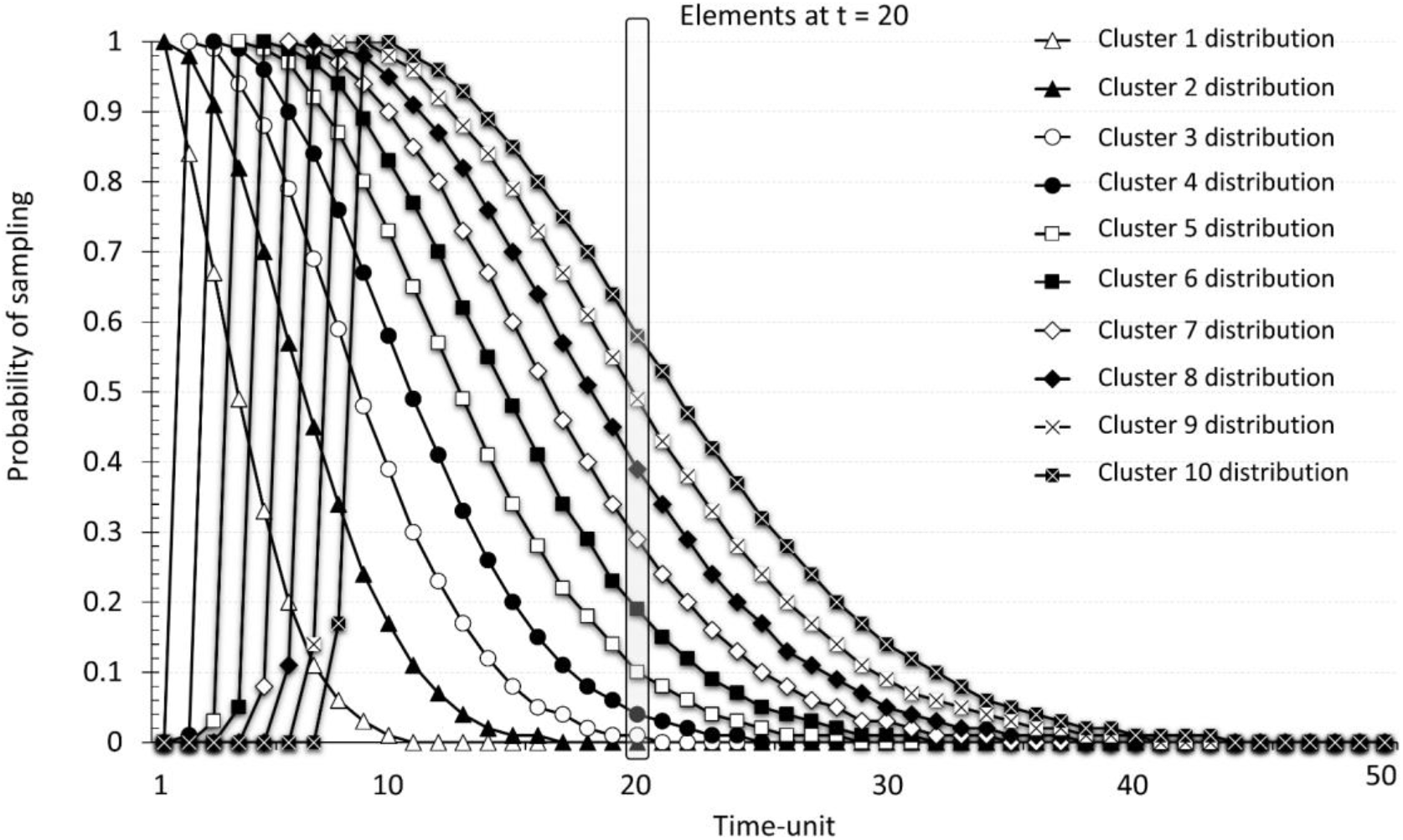
A stimulus consists of a set of clusters of elements. Each cluster is activated in time following an approximately Gaussian probability distribution (temporal cluster). Each temporal cluster peaks at its corresponding time-point of activation and its standard deviation is proportional to it. At any given time-step elements of different clusters coexist.

Temporal clusters are used to represent the idea that the stimulus representation varies in time, so that its elements are differentially active during the presentation of a stimulus. Thus, earlier temporal cluster’s elements can learn differential associations from later ones. Similarly, significant generalization through time occurs as the long tails of the temporal clusters’ activations overlap, allowing the model more flexibility than afforded by step-functions or non-generalizing stimulus activity functions as seen in the original TD model or its CSC extension. These temporal clusters are hence closely related to the ‘microstimuli’ representation of TD (Ludvig, Sutton, Verbeek & Kehoe, 2009). The persistence of temporal clusters’ activations also allows for effective trace conditioning. Thus, the DDA model does not rely on separate eligibility traces to be able to produce trace conditioning as in other models of learning, though it uses a different form of eligibility to counter-act intratrial extinction.

### Associative Activation

Whenever a given element is active, it produces predictions for clusters of elements in proportion to the weights from itself to these clusters. This amounts to a process of associative retrieval that engenders activity in the absence of a direct source of activation. Hence, at any given time there will be elements directly activated by a stimulus physical presence and elements associatively activated by prediction. Obviously, when no training has occurred all weights are presumed to be zero and no associative retrieval takes place yet. At a given time-point, the aggregate of the predictions for a cluster *i* functioning as an *o* of another cluster *j*, denoted by 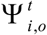, is the total prediction for it and consists of the contribution of each predicting element *e* (of a given cluster *j*, acting as a predictor *p*) active at that time-point 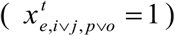. That is, the associative activation of a cluster is a function of the weight of the link from each active predicting element at that moment in time 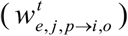, modulated by 0 < *ϑ* < 1, an associatively activated discount that is triggered if the predictor *j* is not directly activated but retrieved. Theta avoids infinite reverberating loops as discussed in (Wagner, 1981, p. 13). The associative activation of a cluster is thus calculated as per Equation 4, where Probability 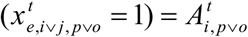

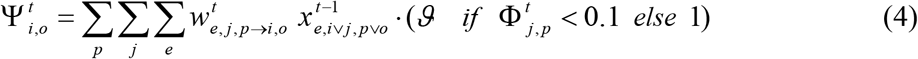

The notation 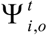 is used here to underline that associatively activated elements are predicted (*o*, outcomes).

### Overall Stimulus Activation

The overall activation of a cluster, either a predictor or an outcome (*i, p* ˅ *o*), is taken to be whichever is larger: its direct activation 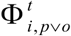, or its associative activation 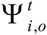, an indirect activation caused by being predicted (output) by other present clusters of a stimulus, as given by Equation 5.

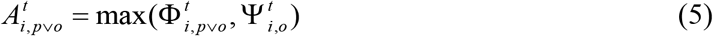

When distinguishing between predictors and outcomes (each stimulus can serve as either), we will use 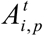 and 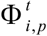, and 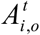 and 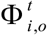, respectively.

That is, the associative and direct activations of a given cluster do not compete, but rather act complimentarily in terms of invoking activity of the elements of a temporal cluster. Hence predictions formed by one stimulus to another do not inhibit the activation of the predicted stimulus as in models such as SOP (Wagner, 1981) or McLaren-Mackintosh (2000), but rather simply reduce the novelty of the predicted stimulus.

### Spread of Activation in the Network

As described in the previous section, a cluster of elements of a stimulus can be activated in two ways: directly from the activity engendered by external sensory stimulation, and associatively from a secondary source of activation produced by an active internal element linked to it. As a consequence, absent but associatively active stimuli can undergo learning. In other words, a given configuration of active stimulus representations can activate other representations through associative links, which have formed due to prior learning. This ‘associative chain’ can propagate to one further cluster (forwards or backwards) at each subsequent time-step. As learning occurs in the model between any concurrently active sets of elements, this activity propagation equips the model with the capability of producing mediated and higher-order learning between stimuli (for instance, A and C in the example in Figure 6). The propagation of activation is however attenuated by dissipation produced by the associative/prediction discount parameter *ϑ*, which multiplies predictions from one node to another if the predicting node is not directly activated by an external source, and by the weight between nodes tending to have a value lower than 1, which renders a weaker activation than that generated by stimulus presence.

**Figure 6:**
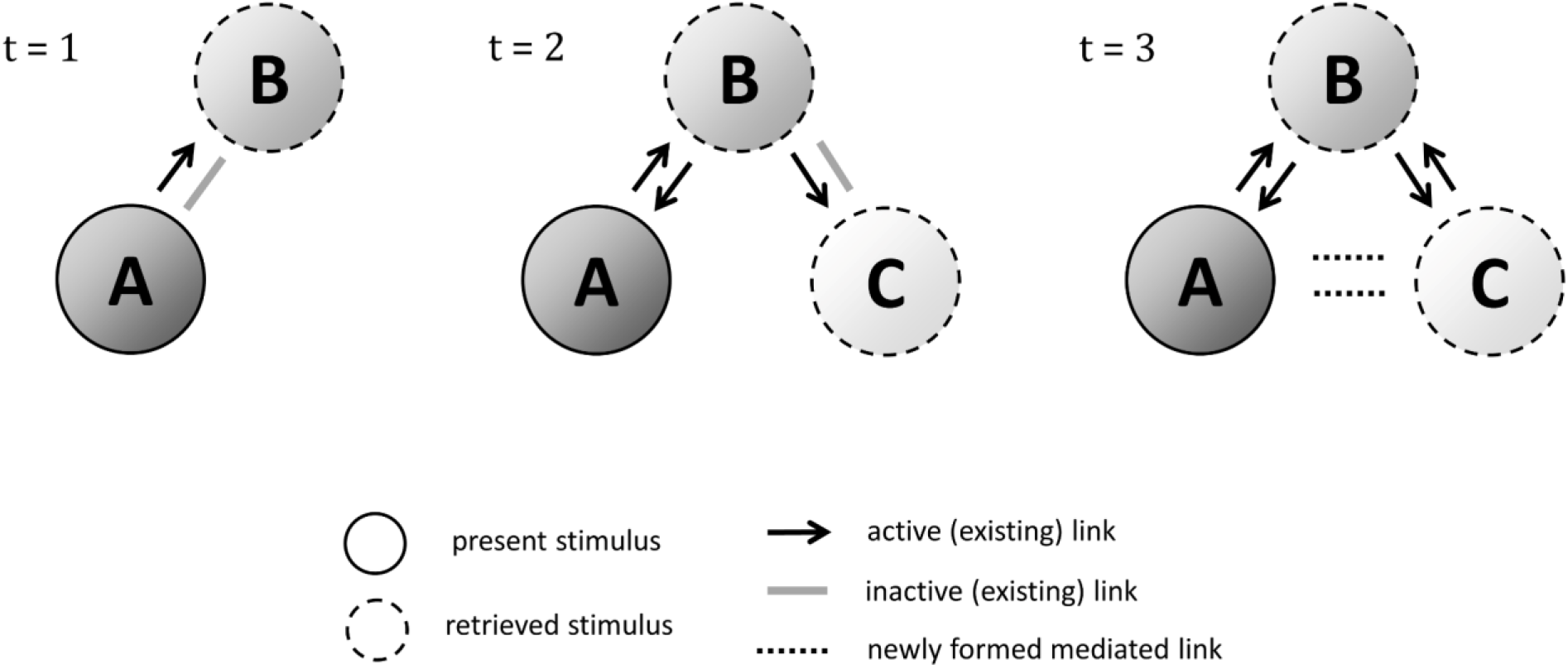
Example of the spread of activation in the network. A time *t* = 1 a cluster node activated by stimulus A’s presence (direct activation Φ) engenders Ψ activity (black arrow) into a node of an absent stimulus B, that is, A retrieves an associative representation of B (an outcome of A) through a pre-existing but inactive link (grey line). Next, at *t* = 2 the now associatively active B spreads activation into an associate C (by means of an existing inactive link). At *t* = 3 A can form new mediated links (dotted lines) to the third-order active C.

### Learning and the Dynamic Asymptote

Learning occurs in the model whenever two clusters are concurrently active either through sensory experience or associative retrieval. Associations develop between an element and a cluster, but since the element’s probability of activation is given by its cluster’s temporal activity distribution, the cluster’s activity is used (instead of the element’s absolute binary activation) to calculate the asymptote of learning so that element-to-cluster learning is modulated continuously in time. The direction of learning (excitatory or inhibitory) between clusters is dependent on the dynamic asymptote, which is a measure of closeness between the aggregate activation of the clusters, as well as the associative strength of other cues in the error term. That is, if two temporal clusters have similar activities, associations between the elements of the predicting cluster and the outcome cluster will develop more rapidly and eventually reach higher asymptotic strength. Hence, two temporally overlapping present stimuli will undergo strong excitatory learning, while a retrieved and a present cue will support a significantly lower maximal level of conditioning. This mechanism derives from the idea that elements (seen as attributes of stimuli) with a similar cluster activation level are equally probable of being present at that moment (with the cluster serving as a more persistent abstraction), and hence belong in the same ‘causal modality’. The power of the dynamic asymptote is that it predicts the same A1→A1 and A2→A2 learning (though it does not presume SOP activation states) as Dickinson and Burke’s SOP learning rules in most cases, while being able to produce A2→A1 learning governed by both Dickinson and Burke’s and Holland’s rules depending on the preceding training (i.e., the A2 stimulus’ associative strength towards the A1 stimulus prior to A2→A1 conditioning). To reiterate however, the model’s behavior in terms of excitatory and inhibitory learning derives from the error term of the outcome; thus no ‘learning rules’ are used. As the retrieved A2 stimulus will usually have a lower activation than if it was directly present, its dynamic asymptote towards the A1 stimulus will have a lower value as compared to A1→A1 conditioning. Therefore, its associative strength will approach this intermediate value. If its prior associative strength is lower than this intermediate value (e.g., as in a mediated conditioning design) it will gain associative strength, and if its prior associative strength was higher (e.g., as in a backward blocking design) it will lose associative strength. Hence by presuming that the maximum supported extent of learning is in fact dynamic and proportional to the degree to which two stimuli are equi-present either directly or through associative retrieval, various learning phenomena can be explained in a parsimonious and emergent manner. Further, the strength of a retrieved association is crucial for the extent of excitatory learning between a retrieved cue and a present stimulus. Neural data for stronger retrieved representations tending to be more associable has been gathered by Zeithamova, Dominick, and Preston (Zeithamova, Dominick, & Preston, 2012). They found that the degree to which a cue was associatively reactivated was correlated with subsequent performance on a predictive inference task involving the retrieved cue and a present outcome.

The asymptote of learning used in the outcome error term is an inverse measure of the distance in activity between the predictor cluster and the predicted cluster (Equation 7). For each type of cluster, the asymptote is estimated using a constrained overall stimulus activation, 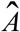, such that if the stimulus direct activation is above a threshold (0.1) its value is set to the maximum direct activation (Equation 6).

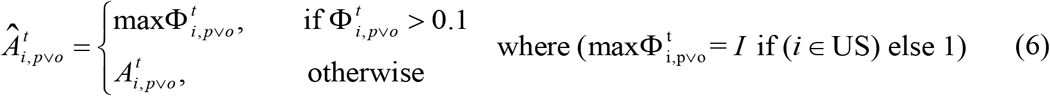

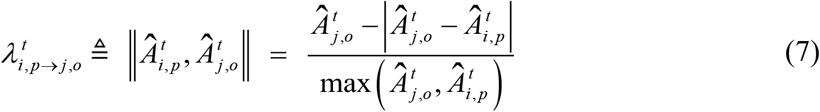

The result is an asymptote based on a linear distance function, with two cues with highly dissimilar activations supporting less learning than if their activations at a given time-point were more similar. This rule is not a strict inverse distance function however, as the absolute value being subtracted from the outcome activity breaks symmetry and allows for differential learning between a present predictor and absent outcome (extinction), and an absent predictor and present outcome (mediated conditioning). As seen in Figure 7, the asymptote’s values range between −1 and 1. The former is produced when the outcome representation is completely inactive, while the predictor has an activity level of 1. The latter is produced when the outcome and predictor have the same activity level. As the absolute value is subtracted from the outcome’s total activity level, this dynamic asymptote is anti-symmetrical, i.e., the outcome activity is more determinant of whether the asymptote is positive or negative.

**Figure 7.**
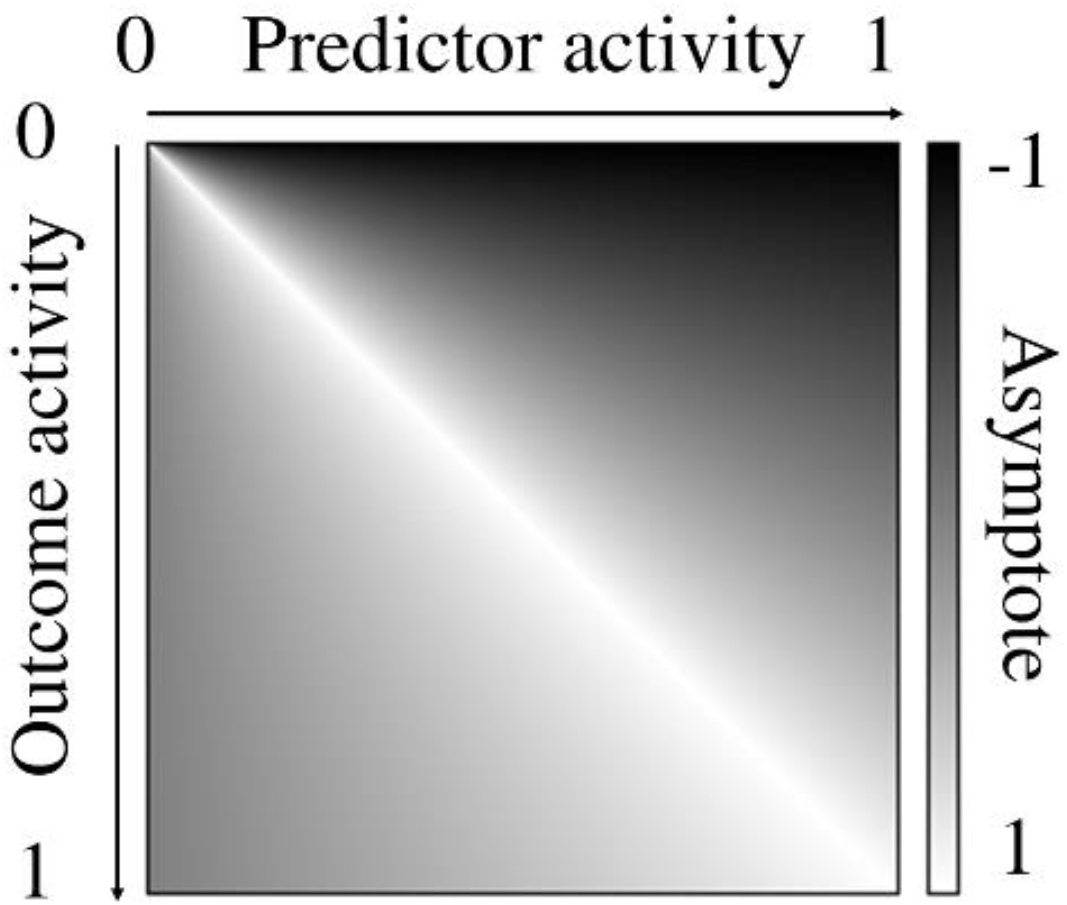
Asymptote of learning as a distance measure of the activities of the predictor and the outcome.

### The Double Error Term and Weight Update

The outcome-error (i.e., prediction error) from an element *e* of a temporal cluster *i* of a given predictor *p* to a temporal cluster *j* of a given outcome *o* is calculated, per Equation 8, by the discrepancy between the asymptote of learning 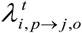 and the total prediction for the outcome 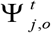.

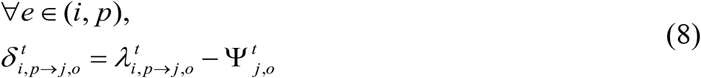

As the asymptote of the model operates through a distance function of cluster activities, and these activities persist through time, the model does not utilize the form of gradient learning incorporated by the error term of the TD model. In the DDA model, the weights of elements from one cluster to another cluster linearly encode the degree to which activity of the predicting element predicts the activity of the outcome cluster. That is, no information about the change in activity over time is encoded in the weights directly, as occurs in the TD model.

Uniquely in the DDA model, per Equation 9, the predicting cluster itself has an error term, the predictor’s error, denoting how expected the predictor stimulus is. The notation →*i, o* is used to emphasize that the predictor’s prediction takes the predictor as an outcome of other elements. This error term is used both directly to modulate learning in the model, as well as to define the revaluation alpha update.

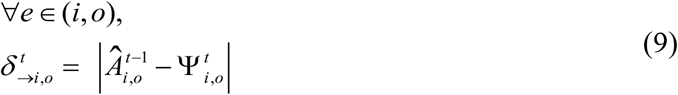

These predictor and outcome error terms are then used in the weight update, Equation 10, for a given (sampled) element of a given temporal cluster, along with the saliences, *s*_*e*,*i*_ and *s*_*j*_, of the predicting element and predicted cluster respectively^2^, the clusters’ activations, 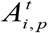 and 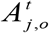, the element binary activity, 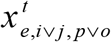, and eligibility term, 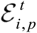, used to counteract extinction before the outcome occurrence (defined in Equation 13), as well as the adaptive revaluation rate, 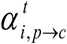(defined in Equation 15). A backward discount *b* multiplies learning from 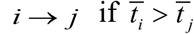, that is if the cluster *i* occurs after cluster *j*.

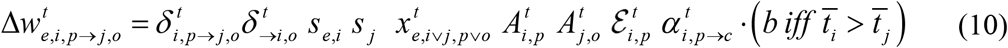

where the values of saliences *s*_*e*,*i*_ and *s*_*j*_ are each a fraction of the stimulus’, including the context’s salience, and are defined in Equation 11. The stimulus saliences are displayed in Table 2 as *S*_S_ in which `S` stands for stimulus A, B, etc.

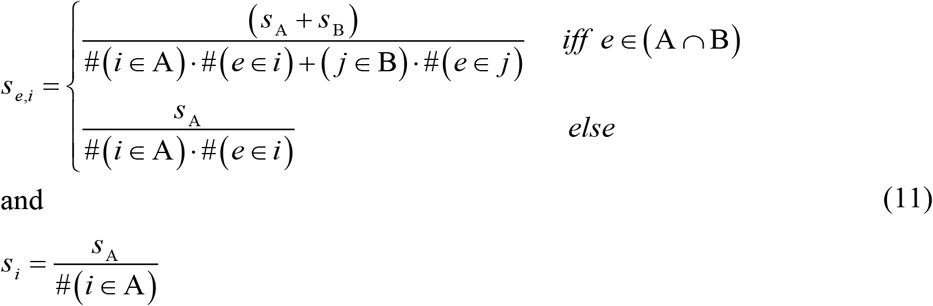

The direction of learning is determined by the outcome prediction error and its variable asymptote, while the prediction error for the predictor itself, along with other modulating factors, influence the extent and speed of learning. As such, the novelty of the temporal cluster the predicting element belongs to plays a crucial role in determining the rate of learning. That is, more novel cues are more readily associated with an outcome. The source of this novelty, namely the extent to which other cues have formed associative links to it, is fully accounted for by the model. This critically allows the DDA model to explain preexposure effects such as latent inhibition in a parsimonious manner.

### Eligibility modulation

As the DDA model learns in real time, significant extinction occurs before the onset of an outcome due to it being predicted yet absent. To counter-act this trend, we have introduced an eligibility factor, which is given by the current prediction for the outcome divided by the maximal prior observed prediction for the outcome in previous trials (calculated using a moving average), raised to a fixed exponent *z* (a value of 2/3 was used in the model). Equation 12 gives the formulation for the observed prediction from one cluster to another (a sum over the elements’ predictions), while Equation 13 formulates the eligibility. For the eligibility, we do not use the overall prediction defined in Equation 4 for all possible predictors of an outcome, but rather eligibility modulation is defined as operating on the basis of cluster-to-cluster temporal predictions. When the predictor cluster is not present but associatively retrieved, Equation 12 is multiplied by *ϑ*.

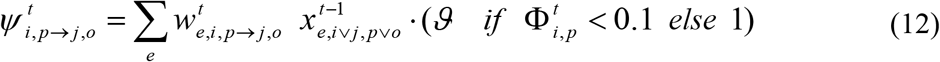

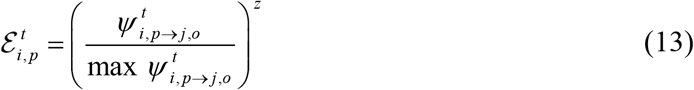

For a given predictor, *p*, while it is active, max 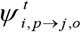 for each outcome is updated toward the maximal prediction for that outcome cluster in the current trial (*T*). The rate of this update is determined by the eligibility discount *γ* as follows:

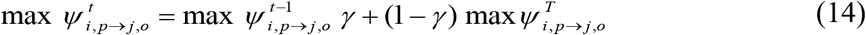

### Attentional Modulation: The Stimulus Associability

The revaluation alpha of the model works by the principle that attention to cues increases when there is uncertainty in the occurrence of the outcome. It further postulates that if the uncertainty remains high over a sufficient period (as tracked by a moving average), the individual uses this persistent uncertainty as a source of information and reduces its level of attention to the cue.

The model states that attentional variables are specific for links to reinforcer and to non-reinforcer stimuli. Formally, they are defined according to Equation 15 which is algorithmically implemented as *α*_*r*_ and *α*_*n*_ depending on the class of outcome, a reinforcer or a non-reinforcer. In the latter case, when the predictor is a US the attentional variable is implemented with the initial value of *α*_*+*_ as given in Table 2. Thus, Equation 15 is defined for *i, p*→*c*, where *c* stands for all active clusters of that class. When the occurrence of a reinforcer is uncertain, the speed of acquisition of excitatory or inhibitory links from a stimulus to that reinforcer increases. Likewise, uncertainty in the occurrence of a non-reinforcer similarly increases the learning speed towards it. If this uncertainty however remains high, the level of attention decays again. The change in the attentional modulation is in proportion to the time-dependent activation 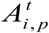 of the element as well as *ρ*, which is a fixed adaptation rate parameter determining how quickly the revaluation alpha changes. The overall direction towards which *α* changes is determined by the overall moving-average error of its respective class of cues.

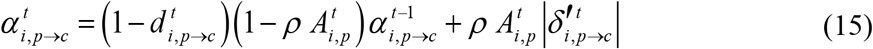

where 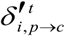 is the overall error of the class of outputs calculated as a moving average in time, and 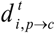 is a decay that kicks in if the moving average crosses a threshold, *ξ*. For instance, during partial reinforcement it leads to *α*_*r*_ decaying after sufficiently many trials when the learner realizes that the contingency is inherently random and therefore warrants less attention. Thus, the crux of the introduced alpha is that an individual can turn a lack of certainty over the occurrence of cues into a source of information in itself. The overall error of a class of cues, 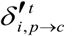, is updated following Equation 16 and Equation 17, where errors are aggregated differentially depending on the class (reinforcer or non-reinforcer) of outcome cluster they belong to.

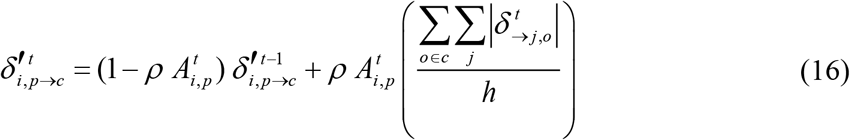

where *h* is the number of reinforcer or non-reinforcer temporal clusters.

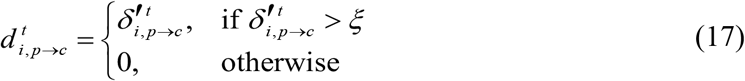

Hence, the DDA model assumes that selective attention, and therefore associability, towards reinforcers and non-reinforcers is proportional to the overall uncertainty of each type of outcome. The variable *α*_*r*_ is thus similar to the Pearce-Hall rule, especially during an acquisition and extinction procedure (left panel, Figure 8), but the DDA model goes significantly beyond it by postulating a similar and independent process for non-reinforcing cues (using *α*_*n*_), as well as assuming that sustained attention as produced in a partial reinforcement schedule declines if uncertainty remains high long (right panel, Figure 8). It further predicts that for each class of outcome, changes in associability induced by uncertainty in one outcome will influence the associability of cues towards other outcomes. In psychological terms this implies that attention (and hence the rate of learning) toward classes of outcomes is proportional to their time-averaged uncertainty. This attentional process operates separately on reinforcers and non-reinforcing classes of outcomes, that is a higher rate of attention to reinforcers does not decrease the absolute rate of attention to non-reinforcers. Hence, during an acquisition procedure, for instance, the speed of learning between a CS and the US is influenced by three factors: 1) the prediction error for the US decreasing over trials; 2) the CS becoming predicted by the context and itself, reducing its rate of learning about the US; and 3) the CS associability to the US reducing in proportion to the US error shrinking. In addition, unlike the Pearce-Hall rate, DDA’s associability is generalized over many trials and its value is dependent upon a moving average of uncertainty that conveys an end of the sustained attention when a prolonged consistent high error endures after long training. Thus, the effects of these different approaches to associability only overlap partially.

**Figure 8:**
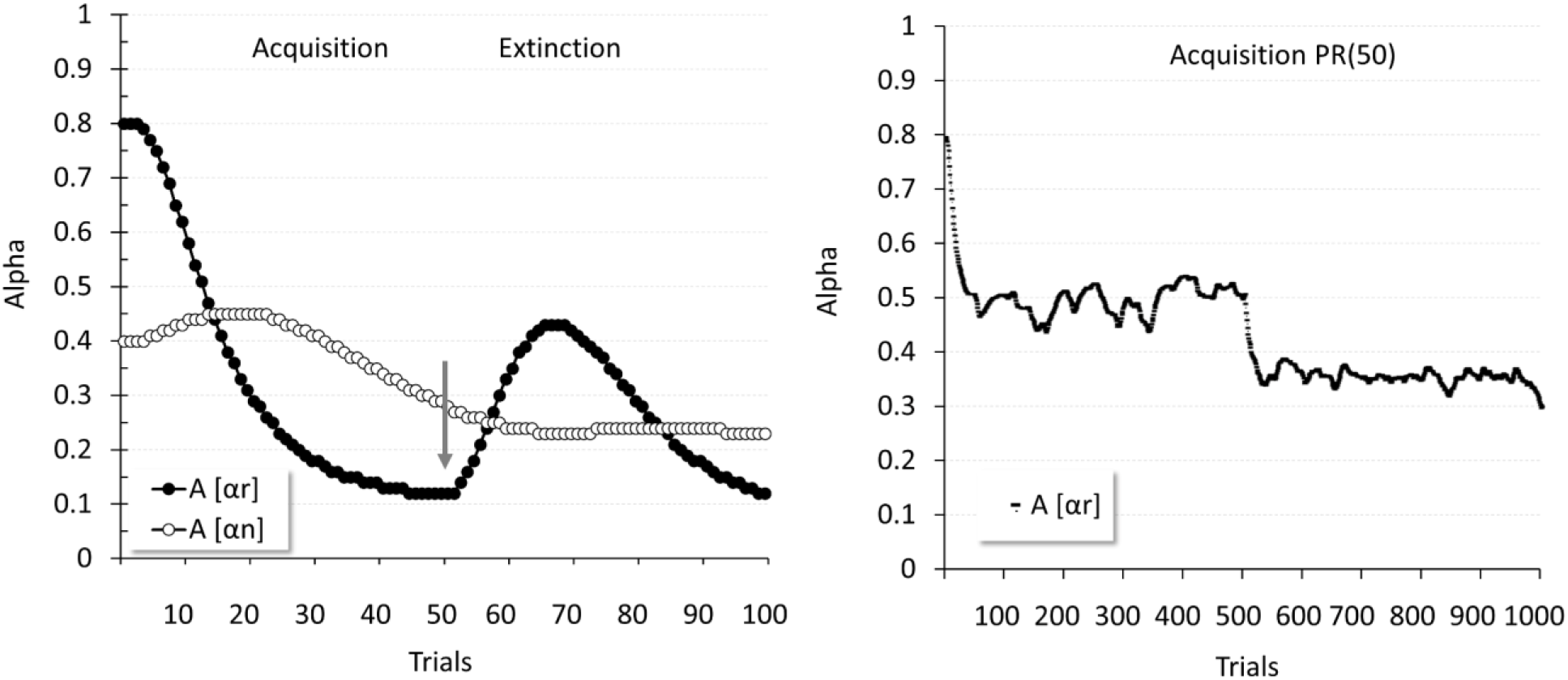
Left panel: Dynamics of the attentional variables of a stimulus A for reinforced and nonreinforced elements (*α*_*r*_ and *α*_*n*_) throughout acquisition and extinction. Right panel: Flow of *α*_*r*_ during partial reinforcement training, showing the revaluation effect produced by the moving average, which reduces attention with persistent uncertainty.

### Computation of Trial Values

To calculate the response elicited on a given trial 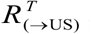, the cumulative predictions of the US^3^ for each and all trial time-points, which is equivalent to the associative activation of the US clusters divided by their number, is averaged over each time-step (Equation 18), and only positive values are taken.

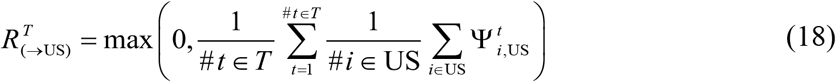

Similarly, to calculate the per-trial prediction made by a given stimulus A for another stimulus B, the cumulative predictions over time for each outcome cluster of the outcome stimulus made by predictor clusters of the predictor stimulus are summed and divided by the number of outcome clusters and time-points in the trial (Equation 19).

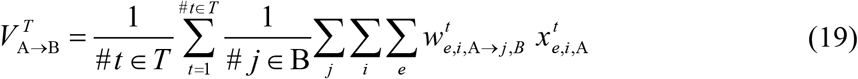

Finally, the per-trial associative link strength from a predictor stimulus A to an outcome stimulus B is calculated equivalently to Equation 19 above, except that the activity of the element is omitted:

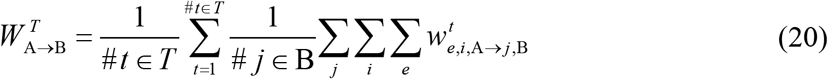

### Model Summary

In summary, the DDA model instantiates a connectionist network consisting of elements, which belong to individual stimuli or are shared between pairs of stimuli, and which are temporally clustered. There are two sources of cluster activation, direct (sensory) activation and associative (cued) activation. At each time-point within a trial or ITI period, the activity distribution of a cluster of elements in time is represented by a semi-Gaussian function with a fixed mean per cluster. The momentary probability of sampling of each element within a cluster equals its cluster’s activity distribution function at a given moment in time. Thus, elements are binary sampled with a given probability that is time-dependent. The associative activation of an cluster is produced by the predictions generated for that temporal cluster at the previous time-point by all other elements of all stimuli. The overall activation of a cluster is then taken to be whichever is larger: the direct activation or the discounted associative activation, and this is used as the resulting probability of sampling elements.

The model next calculates the revaluation alphas individually for each temporal cluster. This occurs in proportion to the activity of the cluster (such that more active clusters experience faster changes in their alpha values). The value that the alphas change towards is the time-averaged mean error of all reinforcers (*α*_*r*_) or non-reinforcers’ (*α*_*n*_) clusters. If this time-averaged mean error value crosses a threshold, the respective alpha value decays at each time-point on which this condition remains true.

Finally, the DDA model calculates the learning between elements and clusters. First, the asymptote of learning is estimated. This asymptote is higher for two clusters with more similar overall levels of activation. This dynamic asymptote enters into the error term of the predicted element, along with the summed predictions from all other elements for the predicted element’s cluster. Similarly, the predictor error term is calculated as the discrepancy between the predictor element’s cluster’s overall activity and predictions made for it by other elements.

The weight from the predictor element to the predicted element’s cluster is then calculated as the product of the two error terms mentioned (with the absolute value of the predictor error term being taken), along with the saliences of the respective stimuli, the overall activations of the two clusters, the revaluation alpha from the predictor to the outcome (i.e., *α*_*r*_ or *α*_*n*_ if the outcome is respectively a reinforcer or a non-reinforcer), the eligibility, and the element’s binary activity.

The pseudo-code and a flow-chart depicting the dynamics for the various processes involved in the DDA model are presented as Table 1 and Figure 9, respectively. Although the DDA model may seem formally complex, its essence is rather simple. Elements of the CS, context, and US are treated for the most part equivalently in that their learning of associations at each time-point is calculated using the same equation. The overall activation of a cluster is simply a factor of its sensory input and predictions made for it by other elements. The revaluation alphas of a given cluster track the overall uncertainty of reinforcing and non-reinforcing events in a straightforward manner. Finally, the direction of learning in the model between two clusters is dictated by the activation similarity of the two clusters along with cue competition; with other factors merely modulating the magnitude of this learning.

**Figure 9:**
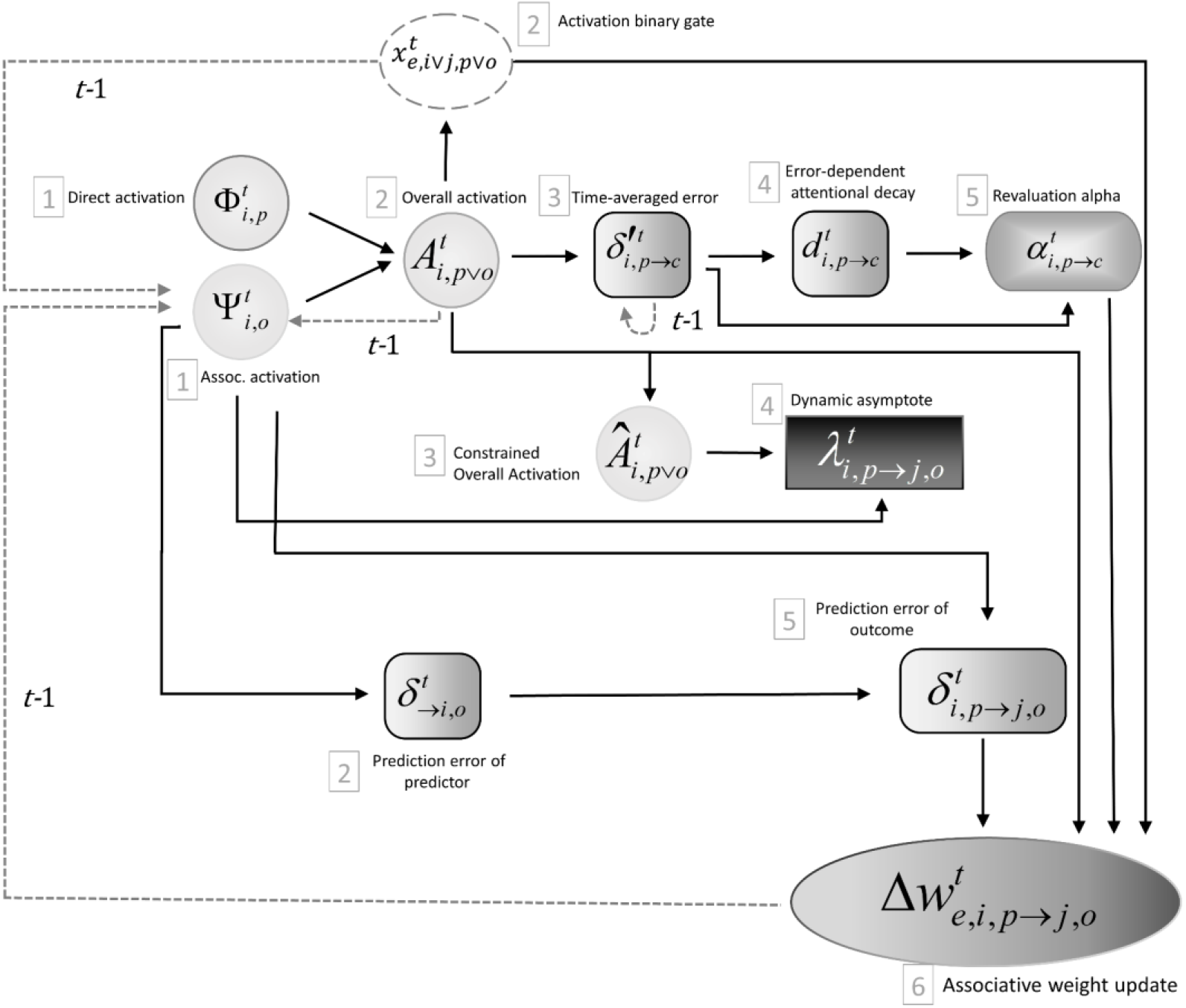
Flow-chart of the dynamics of the DDA model’s equations. Here ‘*t*-1’ denotes that a given update is performed based on values of the previous time-step. Grey numbers show the order in which operations are performed. At each time-point, the DDA algorithm computes both the direct and associative activations of each node (1). The latter takes as inputs the dot product of all weights and activations from the previous time-step (prediction for the node). These two types of activations are used to calculate the overall activation (2) which operationally sets the probability of the binary activation of the element (which determines the cluster activation in a bottom up manner) and modulates the weight update. The overall activation is thence fed into calculating the time-averaged error of reinforcers and non-reinforcers, as well as being passed through a threshold to define the constrained overall activation (a presence value) of a given temporal cluster (3). In parallel, the error term for the predictor (2) is calculated using the associative activation (or prediction) for the predictor. The time-averaged error (3) is used to decide whether a decay in the revaluation alpha is to be triggered (4), and both are used to calculate alpha (5). The constrained overall activation (3) also feeds into calculating the dynamic asymptote (4), which combined with the associative activation of a given outcome is used to calculate the error-term of the outcome in relation to its predictor (5). Finally, the revaluation alpha, total activation, and two error terms input into updating the strength of the weight between any two nodes (6).

**Table 1:**
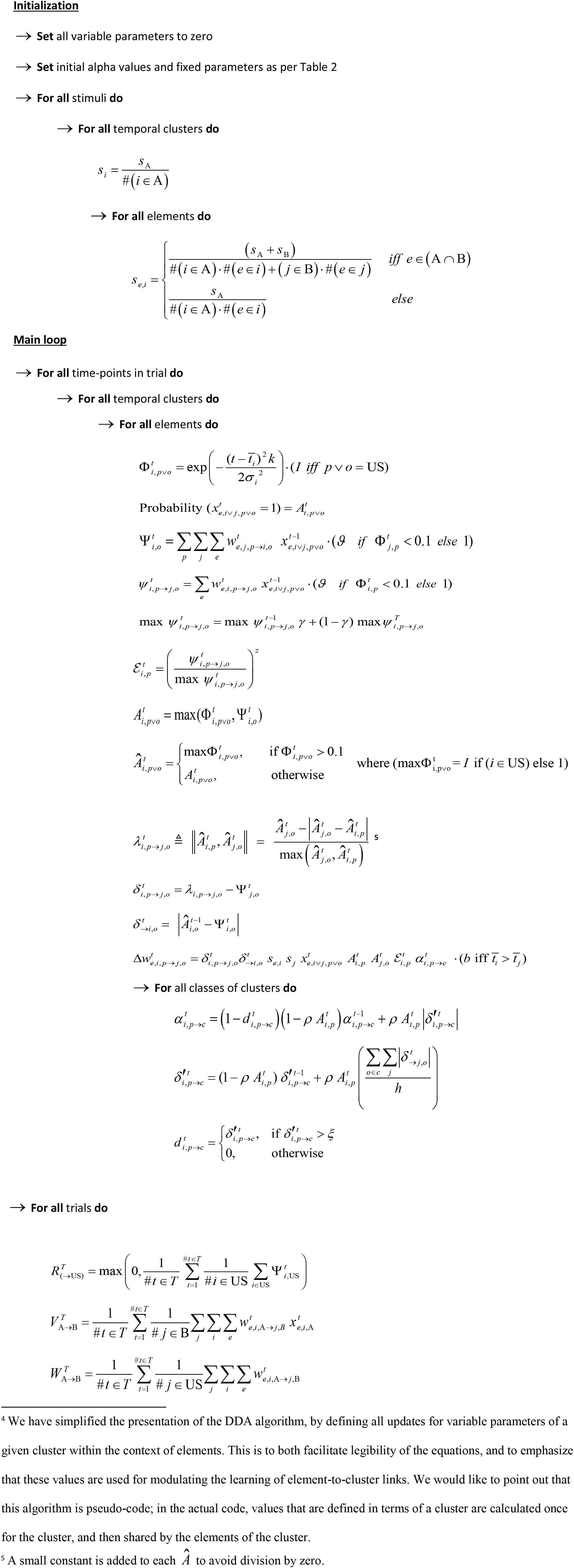
Algorithm of the DDA model with step by step pseudo-code^4^.

Before proceeding to the Results section, which focuses on presenting specific experimental designs, critical to reveal the predictive power of the model and its comprehensive nature, a summary of simulations of some fundamental phenomena of conditioning are presented in Figure 10. These comprise acquisition and extinction of a CR, blocking and unblocking produced by an increase or decrease in the US intensity, ABA renewal, superconditioning, spontaneous recovery along the lines of Bouton’s conjecture (Rosas & Bouton, 1998; Bouton, 2004), contexts effects in latent inhibition, as well as timing effects including the asymptotic level of responding during acquisition being influenced by the ITI length (ITI effects), and superimposition of timing responses (scalar invariance).

**Figure 10:**
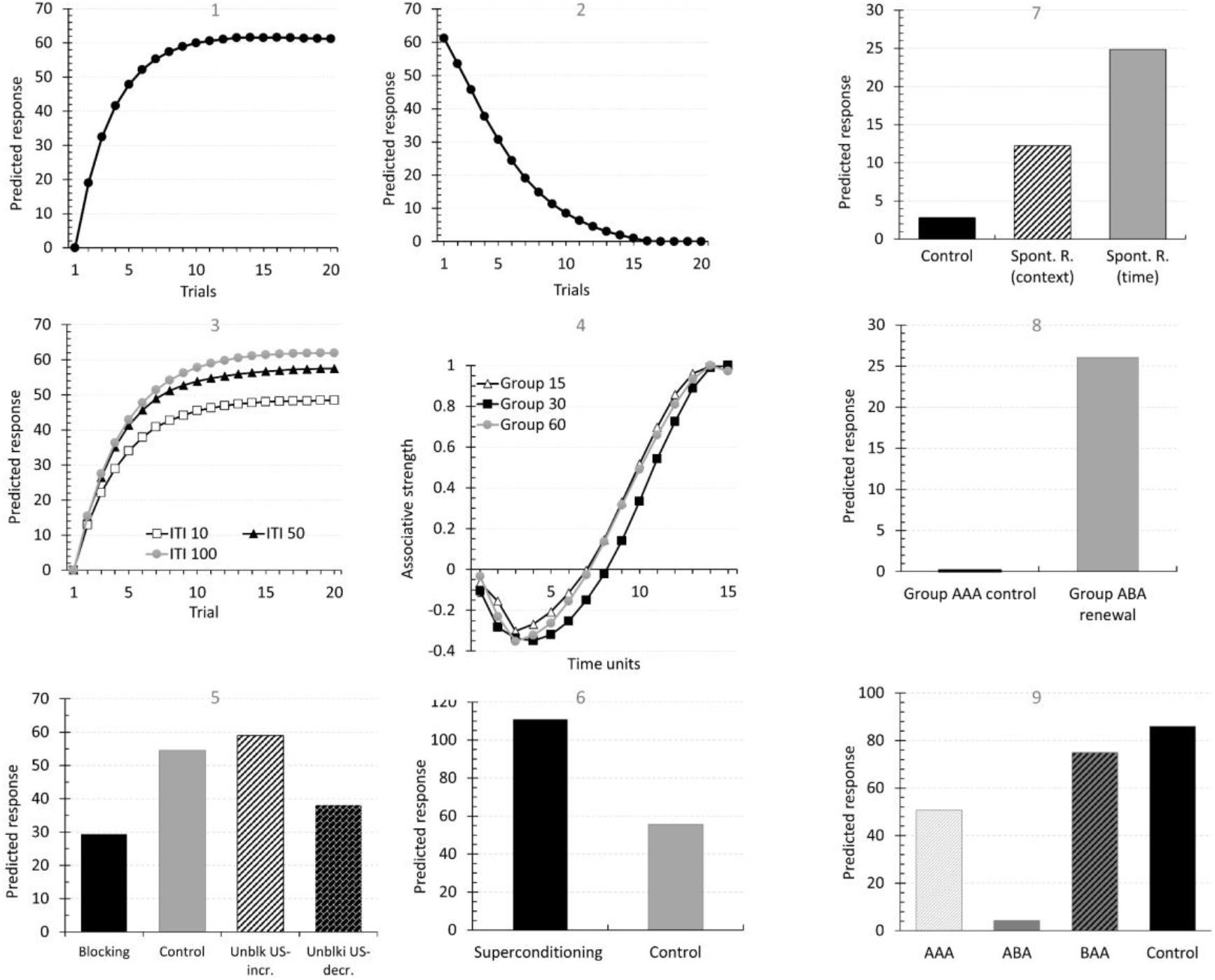
DDA model simulations of conceptual designs of fundamental conditioning phenomena. Panel 1 to 9 are respectively: Acquisition, extinction, inter-trial effects, CR timing superimposition, blocking effects (blocking, unblocking by a US increase in intensity, unblocking by a decrease in the US intensity), superconditioning, spontaneous recovery (by assuming a context change, or by an interval of time in a non-neutral context), ABA renewal, and context effects in latent inhibition.

#### Acquisition and extinction

In explaining the mechanisms of the DDA model involved in accounting for different empirical results, and for the sake of clarity, a set of clusters belonging to a stimulus would be simply referred to as their representation node (A, B, …).

Whenever there is concurrent activity between two stimulus representation nodes, A and B, two directional associations are formed, from A to B and from B to A (with the roles of ‘predictor’ and ‘outcome’ determined by the direction of the link). The change in the weight of each connection depends on the weights and activations of all other active predictor elements of the same outcome (global error term), the predictor’s prediction weight (double error), and the proximity of their respective levels of activation which sets the current asymptotic value (variable asymptote). The associative strength of a connection is the product of the link weight and the predictor’s activity. The strength of a CR is determined by the net associative strength value of all, present or associatively activated, predictors of the US.

During simple conditioning (Figure 10, panel 1), (context) CS→US links are formed between the context and both the CS and the US and between the latter two. Each type of association impacts on the total acquisition response. The context→CS (and also the CS→context) association contributes to the predictor’s error, and the CS→US link and the context→US link both compete and add to the total US prediction, which in turn determines the strength of the CR. However, the context→US undergoes extinction during the ITI. With extensive training and, importantly, with long ITIs the weight of the context→US link becomes slightly negative.

Extinction training (Figure 10, panel 2) results in a loss of the strength of the CS→US weight, but in standard conditions this loss is incomplete because during extinction training the context acquires inhibitory properties towards the US, preventing the CS from total extinction. Full response suppression is thus due to the resulting low weight of the CS→US link in conjunction with context→US inhibition.

#### Timing effects

A well-known time effect in Pavlovian conditioning is that acquisition is stronger when trials occur spaced than when they are close together in time (Figure 10, panel 3). The DDA model predicts this effect due to the differential amount of extinction that the context undergoes during the ITI. The longer the interval, the more extinction occurs, and therefore, the less the context would compete with the stimulus to acquire associative strength.

In timing studies, another principle phenomenon is the scalar invariance effect that refers to the observation that the standard deviation of the response rate distribution in time increases proportionally to the time interval. As a result, superimposition of responses is observed when their rate is normalized in relative time (Gibbon, 1977). As the DDA model theorizes semi-Gaussian curves which preserve the ratio between the variance of the curve and its mean, the model is capable of producing approximate scalar invariance as shown in Figure 10, panel 4.

#### Blocking and unblocking

Figure 10, panel 5 displays simulations for blocking and unblocking procedures. Blocking occurs when prior conditioning of a stimulus A with a US comes to attenuate subsequent responding to another stimulus B in a test that follows compound training of A and B with the same outcome, in comparison to a control condition in which the initial conditioning to A is omitted. Error correction models assume that this reduction in response reflects a deficit in learning acquisition to B during the conditioning in compound with A, as a result of cue competition.

As an error correction model, the DDA model also accounts for the blocking effect in this manner. The blocking stimulus A diminishes the ability of B to entering into association with the US by reducing the overall error term.

When in a blocking procedure the magnitude of the reinforcer is changed from the initial stimulus training to the compound training phase, blocking of responding to B is reduced, an a more vigorous CR is observed. This phenomenon is known as unblocking (e.g., Holland, 1984).

Predicting unblocking when the shift in the reinforcer is produced by an increase in magnitude is straightforward for error correction models such as the DDA model because it leads to a higher supported asymptote of learning for the US, thus permitting additional excitatory conditioning.

Standard error correction models however fail to predict unblocking when the reinforcer is reduced in magnitude. In such case, B is expected to acquire inhibitory properties towards the US as a consequence of the decreased asymptote. Despite being based on an error correction rule, the DDA model can reproduce unblocking by a decrease in the US magnitude. When during the compound conditioning, the reinforcer is reduced, an inhibitory link is indeed progressively formed between B and the US, which fosters stronger conditioning of the context in relation to that observed when the reinforcer remains unchanged (set to either a high or low value across training). Since in the DDA model the asymptote of learning varies according to the levels of activation, and the weight update includes an error term for the predictor itself, cue competition does not sum linearly, and the amount of excitation that the context can acquire may lead to a net US prediction higher than that attained when the US value is not altered. A further source of secondary excitation contributes to the effect when compared to a group in which a weak US is used through all phases. During the initial training, A reaches a higher asymptotic level of conditioning to the strong US in the unblocking condition in comparison to the one resulting from training with a lower intensity US. On test, the associatively retrieved A adds up to the US prediction by means of an associative chain.

#### Superconditioning

Superconditioning (Figure 10, panel 6) refers to an enhancement or facilitation of conditioning between a CS A and a US produced by pairing them in the presence of a previously established conditioned inhibitor B. In the model, the presence of an inhibitor within the error term enlarges the discrepancy, thus enhancing conditioning over an otherwise normal asymptote. Since all other cues paired with the inhibitor do not have an initial excitation that would counteract the growth of the asymptote, a net increase in the conditioning levels of all excitors will be observed.

#### Renewal (ABA) and spontaneous recovery

Both effects refer to a recovery of a conditioned response that was previously extinguished. In the ABA renewal paradigm (Figure 10, panel 7) extinction is carried out in a context other than that in which conditioning took place. When the stimulus is tested back in the conditioning context, a revival of the CR is observed. As commented above, extinction in the DDA model does not presume a total loss of the stimulus associative strength. Extinction training renders the context with inhibitory properties that protect the CS→US link from total extinction. This effect is particularly strong when extinction occurs in a context other than that of conditioning because the context is unpredicted by the stimulus, and according to the model’s provision surprising predictors are learned about more readily. The context inhibitory value towards the outcome summates to the observed CR, which, as a consequence, is totally suppressed, and full extinction becomes apparent. If the extinguished stimulus is tested in a context other than that in which the stimulus was extinguished, such as in the conditioning context, responding will be recovered due to the removal of the contribution of the inhibitory power conveyed by the context of extinction, provided that the testing context has not undergone similar levels of extinction itself.

In spontaneous recovery (Figure 10, panel 8), the critical manipulation following extinction is simply a time elapse between conditioning and test. Following Bouton’s (1993) suggestion that spontaneous recovery could be understood as a case of temporal context modulation, we carried out two simulations: One in which a novel context was used during test, and another one in which conditioning, extinction and test were conducted in the same context but that included a long retention interval between extinction and test in a different context (during the retention interval, a few conditioning trials to other stimuli and free US delivery were given, as it could be assumed to happen in real experimental conditions). In both scenarios, the DDA model is able to predict the results. If the stimulus is tested in a novel context, responding will increase because the main contributor to the net US prediction will be the CS, protected from extinction by the negative value of the extinction context, whose contribution to the total prediction is no longer present. The model, on the other hand, explains recovery of the conditioned response after a retention interval due to generalization from casual experiences of conditioning to other similar stimuli and from the predictive value of the context in which the retention interval occurs towards similar outcomes.

#### Context effects in LI

Simulations of context manipulations in latent inhibition are shown in Figure 10, panel 9. When preexposure is programmed to occur in a context different from the conditioning and test context (BAA), latent inhibition is attenuated, that is, conditioned responding to the latent inhibited stimulus during test is more robust than that resulting from a training condition in which all phases occur in the same context (AAA). However, if latent inhibition is tested back in the context in which preexposure took place (ABA), latent inhibition is restored, in some cases more vigorously (Westbrook, Jones, Bailey &. Harris, 2000)

The DDA model can account for these effects in a similar way in which it explains context effects in the extinction paradigm. During preexposure 1) both the stimulus and the context lose associability and 2) a strong association is formed between the stimulus and the context. If conditioning takes place in the same context as preexposure, acquisition of the context→US association is reduced by the context loss of associability during preexposure and by the contribution of the CS→context prediction (the DDA model’s double error reduces the speed of learning for expected events). On the contrary, if conditioning occurs in a different context from that of preexposure, a more reliable context→US association is formed due to the combined effect of associability and that the context is unexpected (not predicted by the stimulus). If the conditioning context is novel, its associability would remain intact, allowing for fast and strong conditioning. If the context was preexposed, by itself or with a different stimulus, strong context→US conditioning, although less pronounced, will still be observed because the context is not predicted by the stimulus. Cue competition will warrant that the association between the preexposed stimulus and the US will be more effective when carried out in the same context than if it occurs in a different context. Responding, however, is driven by the total prediction of the US, to which the context strength is added. A larger difference between the strengths of the contexts than between the strengths of the stimulus predictions in both treatments would result in a net higher US prediction following a change of context during conditioning than when the context is unchanged. Crucially, when following a change between the preexposure context and the conditioning context the stimulus is then tested in the same context in which preexposure was delivered (ABA), the contribution to excitation from the context is removed, and thus the total US prediction is given by the stimulus strength which underwent “blocked” conditioning earlier. Hence, conditioned responding in the ABA condition will be lower than in all other treatments.

To conclude, the DDA model does not need to postulate a special type of association between stimuli and “potential consequences”, or to appeal to any specific mechanism to operate in latent inhibition as complementary to those underlying extinction manipulations. (see, Westbrook and Bouton, 2010 for a review).

## RESULTS

This section presents a set of simulations aimed at demonstrating that the DDA model’s unique features enable it to account for a wide variety of the learning effects discussed in the introduction. Simulations were carried out with a universal design simulator DDA Simulator Ver.1. Executable files for the simulator are publicly available at https://www.cal-r.org/index.php?id=DDA-sim, and the code and the data input files for the experiments are deposited in GitHub, https://github.com/cal-r/DDA_model.). Results presented in this section are grouped in three different blocks. The opening block tested the contribution of attentional revaluation and the formation of neutral associations to model preexposure effects. First, latent inhibition and its context specificity are covered, and we build a case for both attentional (Pearce & Hall, 1980) and context mediated learning processes (McLaren & Mackintosh, 2000; Wagner, 1981) being involved in attenuating learning during the acquisition phase of the procedure, as predicted by the DDA model. Next, compound stimulus preexposure during latent inhibition is studied to show the explanatory power of the combination of context mediated learning with a fully connected associative network in predicting phenomena anticipated by some controversial modifications of the PH model (Hall and Rodriguez, 2010). Proceeding from there, simulations of the Hall-Pearce effect are introduced, which we contend is founded on attentional and error-correction processes alike to those operating during latent inhibition, the essence of which the DDA model captures through its revaluation alphas and dynamic asymptote. We continue presenting a simulation in which a mediated negative correlation is observed in a learned irrelevance procedure, which showcases the competence of DDA’s dynamics within a fully connected network to account for phenomena in which seemingly unrelated stimuli, never paired together, can become negatively linked, and how mediated learning in the model can lead to reproduce negative retrospective learning in a procedure formally identical to that of mediated conditioning, but that results in an opposite learning value. Finally, perceptual learning in a within-subject preparation, which poses a challenge to current associative models, is simulated.

The second block of experiments assessed the impact of the model’s dynamic asymptote in predicting the individuals’ capability to revaluate past associations through retrieved representations of cues. Thus, mediated learning procedures were simulated, specifically, backward blocking, unovershadowing, mediated conditioning in a backward sensory preconditioning procedure, and mediated extinction.

Lastly, a third block of simulations examined the competence of the model in coping with complex stimuli and non-linear discriminations such as negative patterning and biconditional discriminations, along with experiments that tested stimulus generalization decrement. This block of simulations was intended to make evident that, with only the assumption of shared elements, the DDA model can learn to approximate so called configural learning. For each phenomenon, we present both the design and the trial-by-trial response values of interest. Simulations are presented with a response measure matching the experimental output. Thus, when required to parallel experimental results, that is when the analyzed response increased inversely to the associative strength, such as in suppression of baseline responding in a conditioned emotional response procedure, or suppression of fluid consumption in a taste aversion learning paradigm, a simulated suppression ratio (*r*) was computed following (Mondragón *et al*., 2014), as seen in Equation 15.

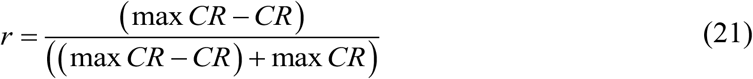

Here, the baseline normalization constant, max *CR* rounded to the nearest five, denotes the maximal level of responding elicited during the test condition, that is, normalized to a scale commensurate to the axis units of the empirical data. Values of *r* closer to 0.5 indicate poor conditioning whereas values near 0 correspond to high levels of conditioning.

The parameters of the simulations are displayed in Table 2 and the design for each experiment is displayed in Table 3. Simulating Experiment 6 requires, due to the large number of cues involved in the design, more RAM than is available on most personal computers. As such, simulating this design needs either the manual disabling of the code for many data arrays normally stored during the running of the simulator or that it be run on a machine with substantially more RAM (e.g., a super-computer).

### Preexposure Effects

The following set of experiments involves the presentation of one or multiple stimuli prior to conditioning. Stimulus preexposure treatments are said to entail variations in the processing of the stimulus, which results in a slow rate in subsequent conditioning of the preexposed stimulus to a reinforcer (e.g., latent inhibition effects, Hall-Pearce negative transfer). When the treatment encompasses purportedly non-correlated CS and US presentations, conditioning is retarded more than would be expected just by the combined effects of the preexposed stimuli (learned irrelevance). Moreover, evidence suggests that, under certain conditions, unrelated presentations of a CS and a US, without an explicit negative correlation, can render the stimuli with an inhibitory relationship, by means of a retrospective revaluation (Baker, Murphy, & Mehta, 2003). Additionally, exposure treatments often result in a reduction of generalization between stimuli (perceptual learning). As such, these phenomena offer the ideal testing ground to demonstrate the model’s mechanism of revaluating the associability rate as well as the modulatory effect fostered by neutral cue associations, namely, context→CS and CS→CS associations on subsequent conditioning to a US.

**Table 2:**
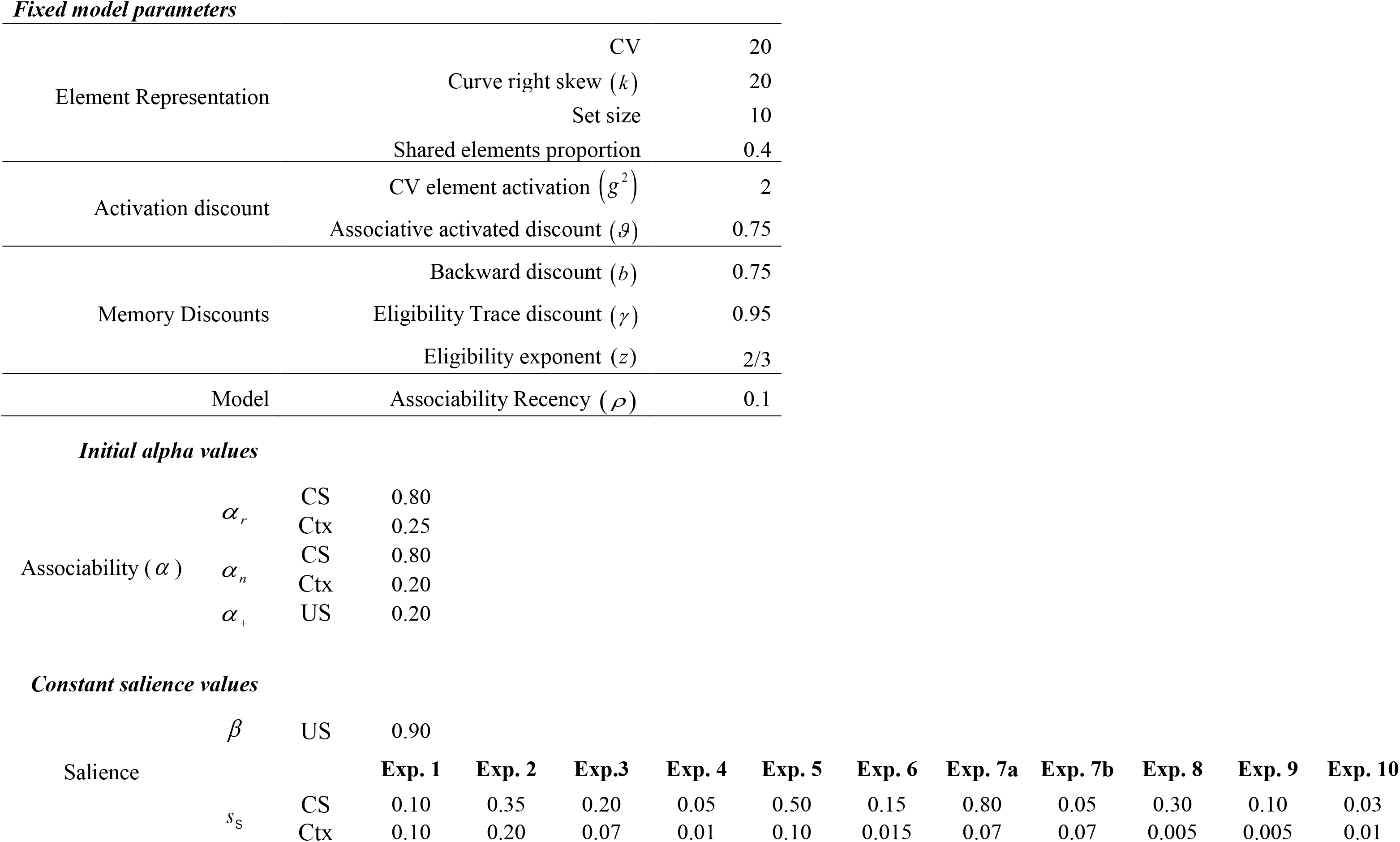
Simulation Parameters. Parameters are not optimized, however salience values differ to adjust for empirical differences of the stimuli used in each experiment as reported in the literature.

#### Experiment 1: Latent Inhibition and Context Specificity

An arbiter of a model’s ability to fully explain the effect of latent inhibition (LI) is whether the model can also reproduce the fact that LI shows context specificity. Models such as Pearce and Hall (Pearce & Hall, 1980) can account for the deceleration observed in latent inhibition, yet fail to explain the effect that latent inhibition is weaker when subsequent conditioning takes place in a different context than that in which preexposure occurred.

In Experiment 3 (Channell & Hall, 1983) two groups of rats were given preexposure training to a stimulus in a distinctive context (Φ) and then in a subsequent phase received appetitive conditioning trials to this stimulus. For half of the subjects (Group Exposed Same) conditioning occurred in the same context that was used in preexposure whereas for the remaining animals conditioning training was given in a different context (ψ)(Group Exposed Different). Two further groups of animals (Group Control Same and Group Control Different) received identical conditioning training but did not receive preexposure to the stimulus. Table 3, Experiment 1, shows the design.

**Table 3:**
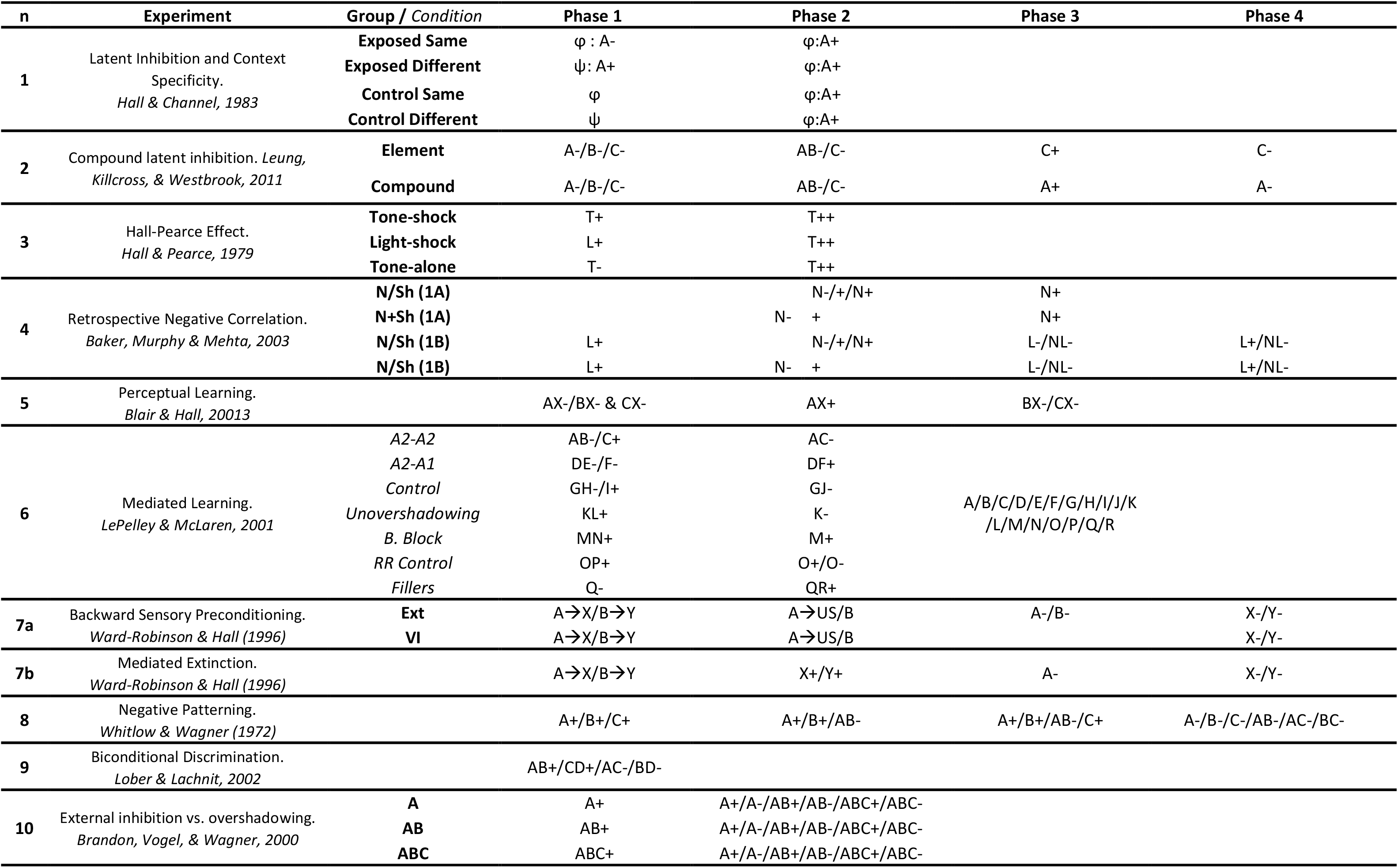
Experimental designs.

| n | Experiment | Group / Condition | Phase 1 | Phase 2 | Phase 3 | Phase 4 |
| --- | --- | --- | --- | --- | --- | --- |
| 1 | Latent Inhibition and Context Specificity.<br>*Hall & Channel, 1983* | Exposed Same | $\varphi$ : A- | $\varphi$:A+ | | |
| | | Exposed Different | $\psi$: A+ | $\varphi$:A+ | | |
| | | Control Same | $\varphi$ | $\varphi$:A+ | | |
| | | Control Different | $\psi$ | $\varphi$:A+ | | |
| 2 | Compound latent inhibition. *Leung, Killcross, & Westbrook, 2011* | Element | A-/B-/C- | AB-/C- | C+ | C- |
|  |  | Compound | A-/B-/C- | AB-/C- | A+ | A- |
| 3 | Hall-Pearce Effect.<br>*Hall & Pearce, 1979* | Tone-shock | T+ | T++ |  |  |
|  |  | Light-shock | L+ | T++ |  |  |
|  |  | Tone-alone | T- | T++ |  |  |
| 4 | Retrospective Negative Correlation.<br>*Baker, Murphy & Mehta, 2003* | N/Sh (1A) |  | N-/+/N+ | N+ |  |
|  |  | N+Sh (1A) |  | N- + | N+ |  |
|  |  | N/Sh (1B) | L+ | N-/+/N+ | L-/NL- | L+/NL- |
|  |  | N/Sh (1B) | L+ | N- + | L-/NL- | L+/NL- |
| 5 | Perceptual Learning.<br>*Blair & Hall, 20013* |  | AX-/BX- & CX- | AX+ | BX-/CX- |  |
| 6 | Mediated Learning.<br>*LePelley & McLaren, 2001* | A2-A2 | AB-/C+ | AC- | A/B/C/D/E/F/G/H/I/J/K<br>/L/M/N/O/P/Q/R |  |
|  |  | A2-A1 | DE-/F- | DF+ |  |  |
|  |  | Control | GH-/I+ | GJ- |  |  |
|  |  | Unovershadowing | KL+ | K- |  |  |
|  |  | B. Block | MN+ | M+ |  |  |
|  |  | RR Control | OP+ | O+/O- |  |  |
|  |  | Fillers | Q- | QR+ |  |  |
| 7a | Backward Sensory Preconditioning.<br>*Ward-Robinson & Hall (1996)* | Ext | A→X/B→Y | A→US/B | A-/B- | X-/Y- |
|  |  | VI | A→X/B→Y | A→US/B |  | X-/Y- |
| 7b | Mediated Extinction.<br>*Ward-Robinson & Hall (1996)* |  | A→X/B→Y | X+/Y+ | A- | X-/Y- |
| 8 | Negative Patterning.<br>*Whitlow & Wagner (1972)* |  | A+/B+/C+ | A+/B+/AB- | A+/B+/AB-/C+ | A-/B-/C-/AB-/AC-/BC- |
| 9 | Biconditional Discrimination.<br>*Lober & Lachnit, 2002* |  | AB+/CD+/AC-/BD- |  |  |  |
| 10 | External inhibition vs. overshadowing.<br>*Brandon, Vogel, & Wagner, 2000* | A | A+ | A+/A-/AB+/AB-/ABC+/ABC- |  |  |
|  |  | AB | AB+ | A+/A-/AB+/AB-/ABC+/ABC- |  |  |
|  |  | ABC | ABC+ | A+/A-/AB+/AB-/ABC+/ABC- |  |  |

The results of this experiment are displayed on the left panel of Figure 11. The conditioning rate was retarded displaying a sigmoidal acquisition shape when the stimulus was preexposed (Group Exposed Same and Group Exposed Different) in comparison to non-preexposed animals (Group Control Same and Group Control Different). However, this retardation effect was attenuated when conditioning occurred in a context other than that used during preexposure. Thus, Group Exposed Same displayed significantly more latent inhibition, i.e., slower conditioning, than Group Exposed Different, which in return showed slight attenuated learning compared to the control groups.

**Figure 11:**
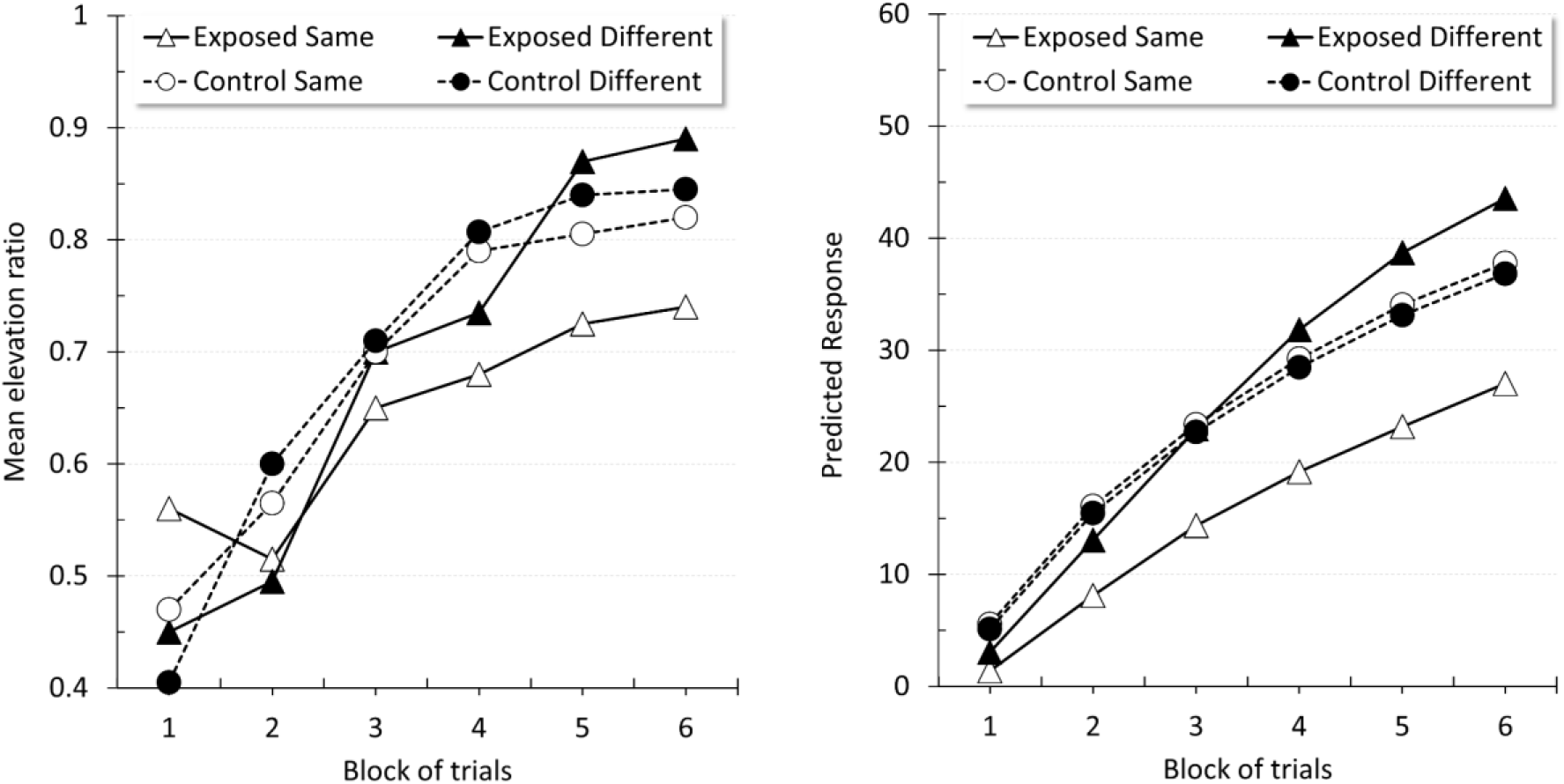
Latent inhibition and context specificity. Empirical (original measurement units) and simulated results during the conditioning phase of Experiment 3, Channel and Hall (1983). The left panel is an adaptation of the published data showing acquisition of a CR in groups Exposed Same, Exposed Different, Control Same and Control Different. The right panel displays the corresponding simulated results generated by the DDA model.

The results of the corresponding simulation are displayed on the right panel of Figure 11. The duration of the CSs, US and ITI was 5, 1 and 50 time-units, respectively, the US intensity was 0.25. The values of the remaining parameters were set as per Table 2, Experiment 1. Preexposure consisted of 100 trials in groups Exposed Same and Exposed Different. Group Control Same and Group Control Different received no training in Phase 1. Phase 2 consisted of 66 trials in all four groups, with the Different groups’ training programmed to occur in a different context. The results of this simulation (Figure 11, right panel) closely matched the empirical pattern. Conditioning in Group Exposed Same was delayed in comparison to the other groups. In particular, the effect of preexposing the stimulus, latent inhibition, although initially evident was considerably reduced with training when the context was changed from conditioning (Group Exposed Different).

During preexposure, repeated presentations of the exposed stimulus in isolation results in a loss of its associability to the reinforcer, *α*_*r*_. As no outcome is expected, there is no uncertainty, which, following the DDA model, would be critical in sustaining the associability of a stimulus. Consequently, during subsequent acquisition trials, the stimulus associability to the reinforcer is higher in groups Control Same and Control Different, which have not undergone preexposure and therefore keep their initial associability intact, than in the preexposure groups for which *α*_*r*_ has decayed. In other words, preexposure reduces the associability of cues to future reinforcers as they have a history of not being causative of, or correlated with reinforcement. Hence, the control groups display faster acquisition partly due to the stimulus’ relatively higher associability towards the US. The model also postulates that the error in predicting the outcome is reduced when the CS is predicted by other stimuli. Thus, in the groups in which preexposure and conditioning occur in the same context, the associations between the context and the stimulus and between the CS elements (unitization), which are formed during preexposure, reduce the speed of the CS→US learning; a decrease which is proportional to the loss of novelty of the CS. As the CS is predicted more strongly in Phase 2 in Group Exposed Same than in Group Exposed Different in which conditioning occurs in a novel context, the rate of learning is further reduced in the former in comparison to the latter, thus resulting in a greater latent inhibition effect when compared to the corresponding non-preexposed control group (Group Control Same). In fact, the degree of latent inhibition is postulated to be proportional to the net effect of unitization, prediction by the context, and loss of selective attention due to lack of reinforcement.

In summary, the DDA model’s associability accounts for the decelerated learning rate observed in LI. Just as importantly, the predictor error term in the model’s learning equation endows it with the ability to predict context modulation of learning, because the CS becomes expected through training in a single context in comparison to when this context is no longer present.

Given that these effects can be disassociated, the model can further predict that latent inhibition would be attenuated (yet not completely abolished) should the preexposure treatment be followed by exposure to the context alone, as this would extinguish the context to CS link. Some evidence supporting this prediction has been reported by Aguado, Symonds & Hall (1994). In their Experiment 3, a retention interval (in home cages) was introduced between preexposure and conditioning, which, in this particular procedure, could be taken as context exposure. This effect would be slightly mitigated since changes in the preexposure regime would raise the uncertainty of neutral cues, thereby increasing *α*_*n*_ of both the CS and the context. As a consequence of the increase in the neutral cues’ associability, stimulus unitization and context to CS learning would presumably be facilitated, thus lessening the attenuation of latent inhibition. Following the same argument, the model can advance a further prediction: Preexposing the CS in multiple contexts should result in stronger latent inhibition, as this would reduce the variable associability of the CS, while retaining the strength of the context to CS links.

#### Experiment 2: Compound Latent Inhibition

Recent results by Leung, Killcross, and Westbrook (2011) have found evidence of a novel prediction derived from Hall and Rodriguez’s model (2010), according to which, when a preexposed target stimulus is further exposed in compound with another stimulus, latent inhibition accrued by the target is larger than if the target is additionally preexposed in isolation. Hall and Rodriguez’s prediction derives from the assumption that preexposure to a CS leads to an association between the CS and an unspecific noUS center. This association would interfere with the formation of links between that CS and any given US. Unlike isolated CS exposure, additional compound preexposure would result in a reduction of the error due to summation of the two CS→noUS predictions, causing a decline in attention to the target CS. As a consequence, conditioning would proceed more slowly. Simulating Leung et. al.’s results is of interest for two main reasons. First, these results challenge accounts, such as the DDA model, that postulate the intervention of the context in producing the effect. Under this assumption, it could be advanced that preexposing a stimulus compound would be expected to lessen, rather than to potentiate, the LI effect due to competition between context-to-CS learning and CS-to-CS learning. Second, it is a matter of theoretical significance to find an alternative mechanism to Hall and Rodriguez’s which requires assuming a controversial unspecific non-US center to predict the result.

Experiment 3 in Leung *et al.* (2011) used an aversive conditioning procedure in rats. In Phase 1, the two groups of animals were presented with non-reinforced random presentations of A, B, and C. Phase 2 consisted of non-reinforced presentations of a compound AB and C. In Phase 3, Group Element received a single reinforced presentation of C (the elemental control), while Group Compound received a reinforced presentation of A (the target). The experiment found that preexposing A in compound with B led to a more pronounced attenuation of subsequent conditioning, that is, to a more robust latent inhibition effect between A and the outcome when compared to stimulus C, which was preexposed in isolation. Figure 12, left panel, shows that animals in Group Element displayed a higher mean percent of freezing, i.e., faster acquisition, than animals in Group Compound during the test phase.

**Figure 12:**
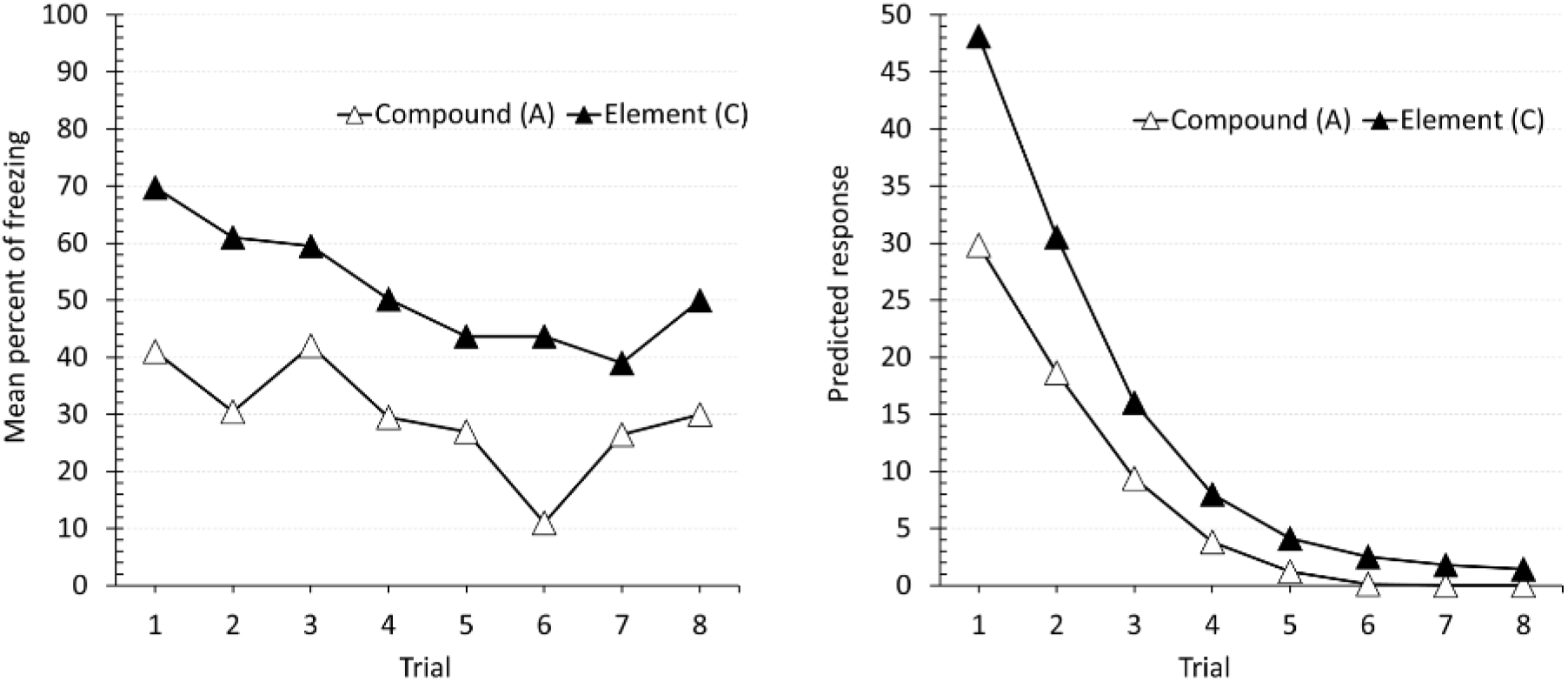
Stimulus compounding in latent inhibition. Empirical (original measurement units) and simulated results of Experiment 3 in Leung *et al.* (2011). The left panel is an adaptation of the published data showing the mean percent freezing in the test phase. The right panel displays the corresponding simulated results generated by the DDA model.

The temporal parameters used to simulate this experiment were a 5 time-units CS, 1 time-unit US (intensity 2.5), and 50 time-units ITI. The remaining parameters were set to the values in Table 2, Experiment 2. Both groups of the design were programmed to receive 10 randomly presented non-reinforced trials of each stimulus (A, B, C) in Phase 1. In Phase 2, both groups were programmed to receive 10 randomly presented AB and C trials. In Phase 3, Group Compound received a reinforced presentation of stimulus A, while Group Element received a reinforced C trial. The programming of the subsequent test phase presented respectively 8 non-reinforced A and C trials for groups Compound and Element. The full experimental design is presented in Table 3.

The simulation reproduced the pattern observed in the experiment. Figure 12, right panel, shows that the predicted responding strength to A in Group Compound was lower than to C in Group Element during the test trials of the last phase, thereby displaying more latent inhibition.

The DDA model’s account of this effect is simple and does not require extra assumptions as those in (Hall & Rodriguez, 2010). Neutral associations between the constituent elements of the compound AB are formed during preexposure, so that the two CSs become predictors of one another. This leads to A retrieving B elements during conditioning in Group Compound, which in return produces a second-order prediction for A elements. Thus, A in Group Compound is predicted by both the context and the retrieved B. Since the DDA model assumes a predictor error term within the error determining the change in the associative weight of a stimulus, it predicts that A would condition at a slower rate in comparison to the speed of acquisition of C in the Group Element, which is predicted only by the context.

This is of considerable interest, as it implies that the DDA model does not require to postulate acquisition of an arbitrary and unspecific CS→noUS association during preexposure (Hall & Rodriguez, 2010) to account for these effects. The DDA model’s unique learning rule, which incorporates the unexpectedness of a predictor (in this case the CS) in the learning equation, thus modulating the rate of conditioning between a predictor and the outcome, suffices.

The suggested mechanism permits the model to make a further prediction: The potentiation of the LI effect observed in this experiment should be considerably attenuated should B undergo a series of non-reinforced presentations after being paired with stimulus A, but before the reinforcement of A. This treatment would weaken the associative connection between B and A, thereby reducing the proportion of A elements cued by B during conditioning, accelerating the rate of acquisition.

#### Experiment 3: Hall-Pearce Effect

The acquisition of a CS→US link does not always proceed monotonically in relation to only the pre-existent associative strength. For instance, preexposing a CS with a weak US has been proved to attenuate subsequent acquisition with a stronger US (the Hall-Pearce effect). The error correction and revaluation alpha processes of the DDA model can reproduce this observation.

In Hall and Pearce (1979) Experiment 2 two groups of rats received presentations of a tone (Group Tone-shock) or a light (Group Light-shock) followed by a weak shock. A third group, Group Tone-alone, received non-reinforced presentations of the tone in isolation. In Phase 2, all three groups of rats received presentations of the tone followed by a strong shock. The design of this experiment is displayed in Table 3. The experiment found that reinforcing a CS with a weak shock retarded subsequent acquisition towards a more intense shock. This attenuation of learning was however less pronounced than that produced by preexposure of a CS. The results of this experiment are displayed in the left panel of Figure 13. Group Tone-alone showed a higher suppression ratio than Group Tone-shock, which in return produced a higher suppression ratio than Group Light-shock.

To simulate this experiment, the temporal parameters used were a 5 time-units CS, 1 time-unit US, and 50 time-units ITI. Aside from these values, the parameters corresponded to those in Table 2. In Phase 1, each trial-type was programmed to occur 66 times with a US intensity of 0.01. Likewise, Phase 2 consisted of 66 reinforced trials, with a US intensity of 0.75.

The simulated results (Figure 13, right panel) paralleled empirical data. Group Tone-alone showed the largest suppression ratio over the second phase, followed by Group Tone-shock. Group Light-shock produced the lowest suppression ratio, thus displaying faster learning.

**Figure 13:**
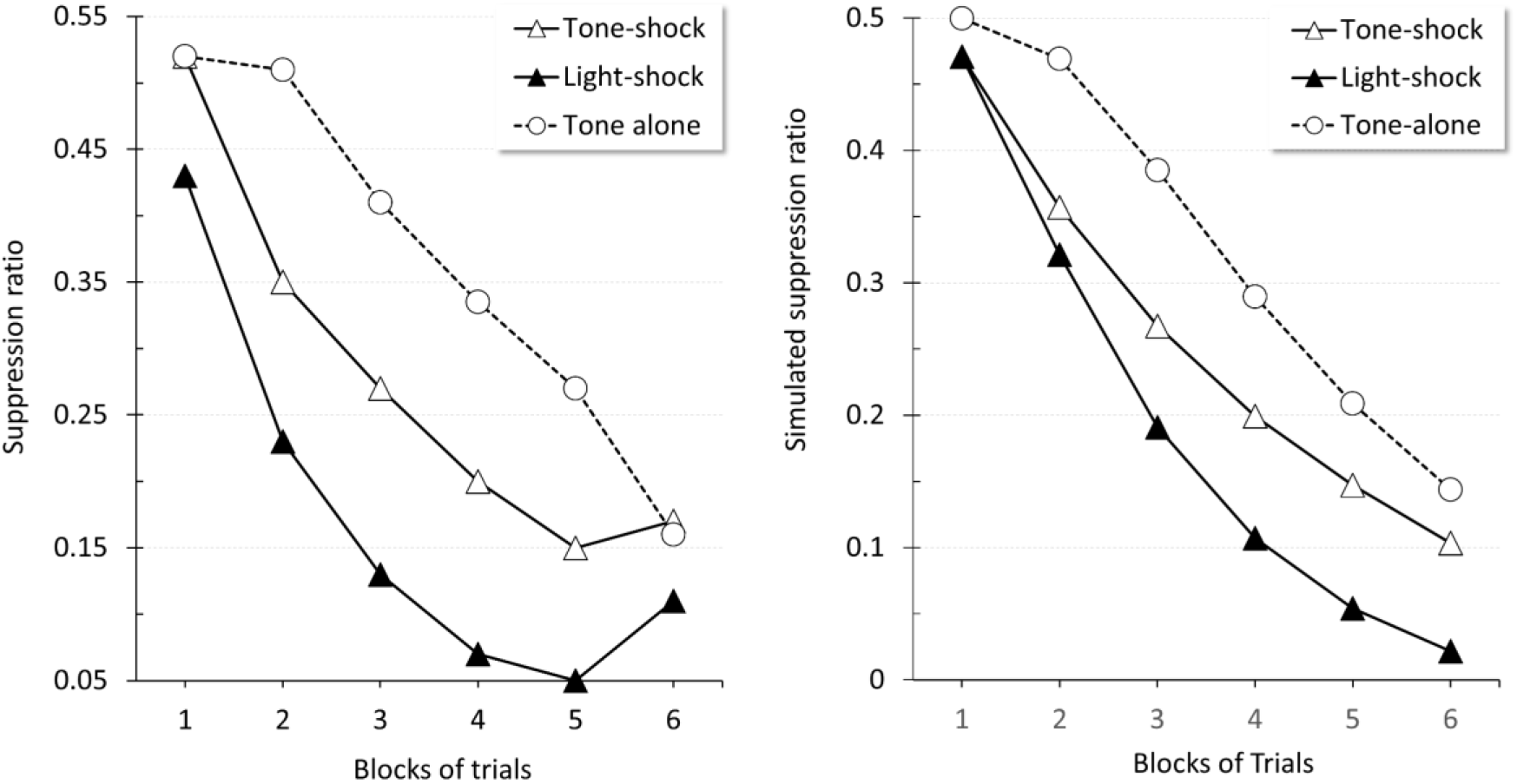
The Hall and Pearce effect. Empirical (original measurement units) and simulated results of Experiment 1 in Hall and Pearce (1979). The left panel is an adaptation of the published data showing the acquisition of a CR in groups Tone-shock, Light-shock, and Tone-alone. The right panel displays the corresponding simulated suppression ratios generated by the DDA model.

The DDA model accounts for this result in the following manner: In Phase 1, exposure to the target stimulus results in a loss of the tone’s novelty in Group Tone-alone and Group Tone-shock. This loss of novelty has a two-folded effect. On the one hand, the tone is predicted by the context at the time of conditioning, which reduces the predictor error term and therefore, the rate of conditioning to the US; on the other, there is a decline in the tone’s associability rate to the reinforcer. Due to the presence of the shock in Group Tone-shock, this decline in the tone’s associability rate is less pronounced than in Group Tone-alone. In contrast, in Group Light-shock the tone has not been preexposed by itself and therefore is neither predicted nor affected by a loss of associability.

Since the effect of the delay in conditioning under these conditions is, according to the DDA model, a product of a decrease in the error and a loss of the stimulus associability, one could predict that would the experiment be modified to have the second phase treatment occur in a novel context, the Hall-Pearce effect would be significantly attenuated (Swartzentruber & Bouton, 1986).

#### Experiment 4: Retrospective Negative Correlation

So far, we have seen effects that can potentially be ascribed to a simple loss in the associability of a stimulus. Next, we are describing an experiment aimed at proving an inhibitory relationship between an absent neutral stimulus and a present reinforcer. The original experiment (Baker *et al.*, 2003, Experiment 1) intended to assess the adequacy of some controls in studying learned irrelevance (Bonardi & Hall, 1996; Mackintosh, 1973). Testing for learning irrelevance has been difficult due to the fact that control treatments involving independent exposure to the CS and the US often entail a veiled negative CS–US contingency. Importantly, the experiment suggests that negative correlations could also appear in schedules that may require revaluating retrospectively the correlation between the CS and the US, such as in blocked stimulus presentations, in which the CS precedes the US. In this section, we simulate Baker *et al.*, (2003), Experiment 1 which shows inhibitory properties between a CS and a US developed in a treatment in which the conditioned inhibitor was cued by the context but not physically present.

The experiment was split into two sub-experiments. Experiment 1a was designed to assess inhibitory properties by means of a retardation test. Experiment 1b was intended to evaluate inhibition with a summation test. During Phase 1 of Experiment 1a animals did not receive any treatment. In Phase 2, Group N/Sh received uncorrelated presentations of a noise and a shock. In Group N+Sh a block of noise trials followed by a block of shock trials were scheduled. In Phase 3, both groups received conditioning trials to the noise followed by the shock (retardation test). In Experiment 1b a light was conditioned to the shock in Phase 1. Group N/Sh and Group N+Sh received identical treatment as their counterparts in Experiment 1a. In Phase 3, all animals were given non-reinforced trials with the noise and a compound noise-light (summation test). During Phase 4 a saving test consisting of reinforced noise trials intermixed with the compound noise-light was carried out.

The retardation test in Experiment 1a showed that uncorrelated presentations of the noise and the outcome (Group N/Sh) produced a smaller deficit in subsequent acquisition than the blocked scheduled presentations in Group N+Sh did. The left panel of Figure 14 shows the retardation test results in Experiment 1a: Group N/Sh displayed a lower suppression ratio (and hence faster learning) than Group N+Sh.

**Figure 14:**
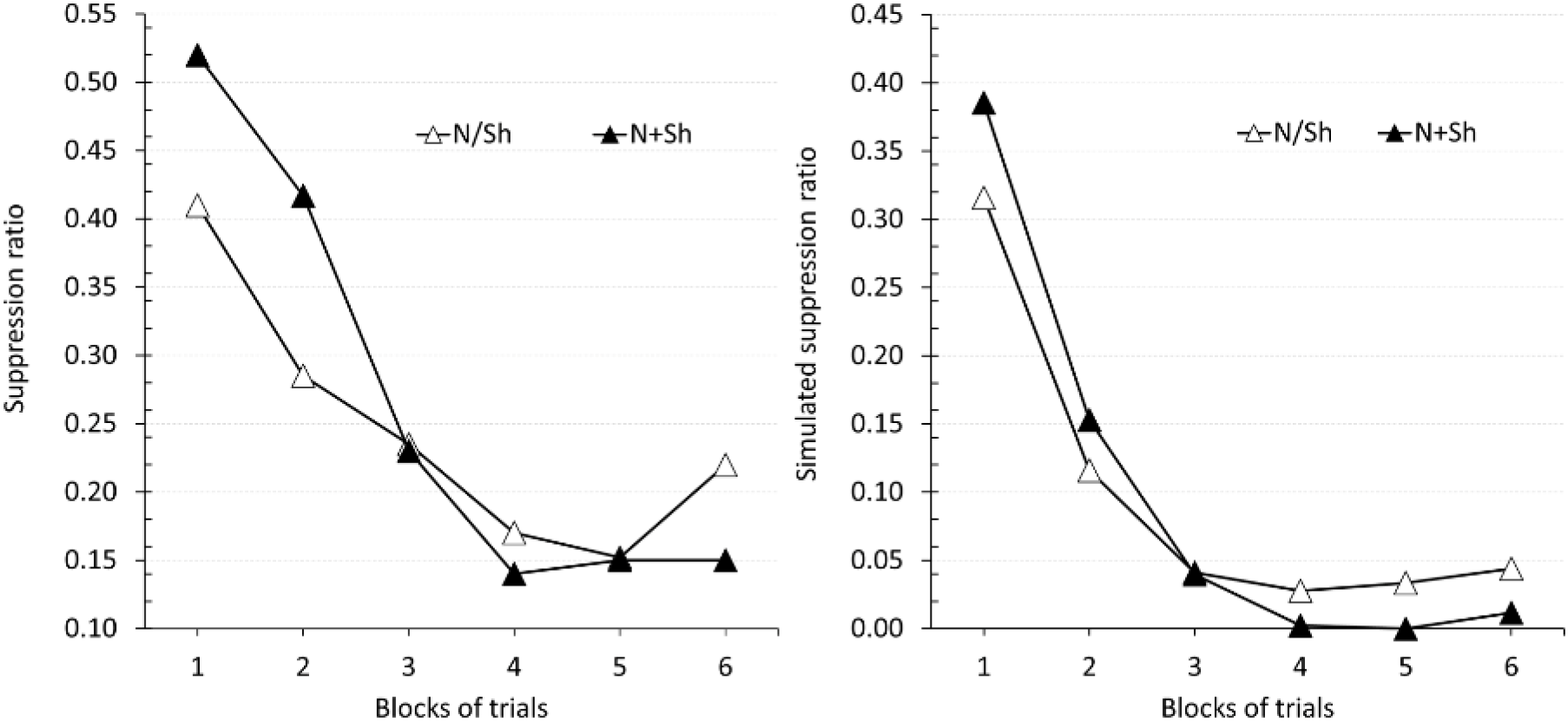
Retardation test in an uncorrelated group (N/Sh) and a retrospective negative correlation group (N+Sh). Empirical (original measurement units) and simulated results of Experiment 1a in Baker *et al.* (2003). The left panel is an adaptation of the published data showing suppression ratios for the retardation test trials to the noise in Phase 3. The right panel displays the corresponding simulated suppression ratios generated by the DDA model.

Figure 15 left panel shows the results of the summation test. Animals trained under a blocked schedule of presentations (Group N+Sh) showed a stronger summation effect, visible as a larger differential responding to the stimulus and the compound. The summation effect was also evident, but to a lesser degree for animals trained in the uncorrelated treatment of Group N/Sh. These results, together with those of the retardation test, suggest the formation of an inhibitory association between the noise and the shock in Group N+Sh.

**Figure 15:**
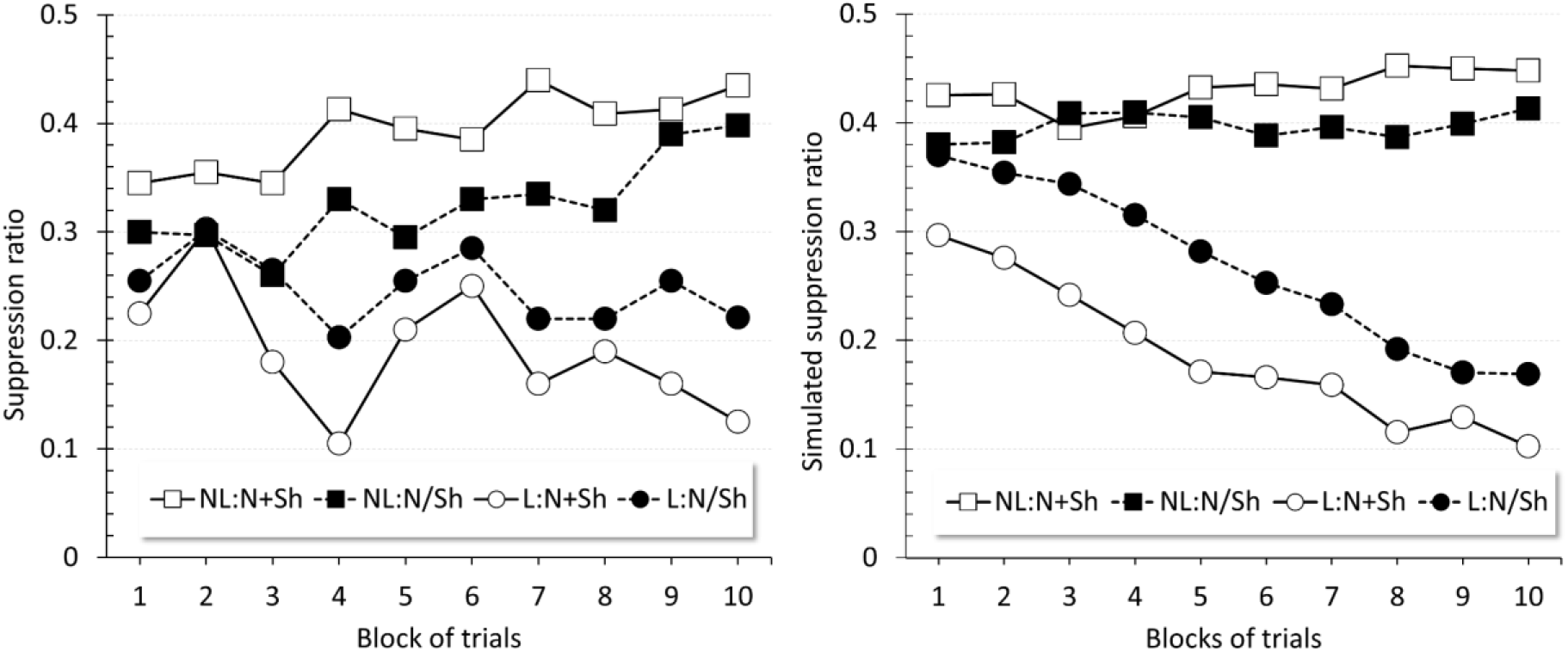
Summation test in an uncorrelated group (N/Sh) and a retrospective negative correlation group (N+Sh). Empirical (original measurement units) and simulated results of Experiment 1b in Baker *et al.* (2003). The left panel is an adaptation of the published data showing suppression ratios for the summation test trials. The right panel displays the corresponding simulated suppression ratios generated by the DDA model.

For the simulation, temporal parameters consisted of a 5 time-units CS, 1 time-unit US (intensity 1), and 50 time-units ITI. The remaining parameters were set to the values in Table 2. For Experiment 1a no event was programmed to occur in Phase 1. In Phase 2, Group N/Sh received random presentations of N and of the US, such that both stimuli coincided on 16 trials. Half of these US presentations occurred before N, and half following it. Group N+Sh received a block of 24 N trials followed by a block of 24 US trials. In Phase 3 both groups received 36 reinforced N presentations. In Experiment 1b, Phase 1 consisted of 8 reinforced L trials. Phase 2 was programmed identically to that in Experiment 1a. In Phase 3, non-reinforced randomly presented trials of a compound NL and two of L were programmed in both groups. Finally, Phase 4 was programmed to deliver 40 non-reinforced NL and 20 reinforced L trials randomly presented. The full experimental design is depicted in Table 3.

The simulation results match empirical data. In the retardation test (Figure 14, right panel), Group N+Sh showed a lower suppression of responding than Group N/Sh.

Results for the summation test (Figure 15, right panel) closely matched the experimental results. Group N+Sh showed a larger difference in simulated suppression between the NL and L trials than Group N/Sh, suggesting that the noise had become a conditioned inhibitor of the outcome in Group N+Sh.

The DDA model accounts for this result by assuming that an animal learns to approximate the CS-US relationship, which in the case of the N+Sh groups would imply retrospectively assessing the information in the following manner: In this group, there will be many US trials in which N is predicted by the context, yet is not present. Thus, the context associatively retrieves N but its activation is weak. The discrepancy between the activation levels of N and the US would yield a minute or negative asymptote, engendering inhibitory learning between N and the US, which combined with competition by the context would allow the model to replicate the result. In contrast, N in N/Sh groups sometimes coincides with the US, therefore preventing the formation of an inhibitory link, or rendering it weak. Additionally, the difference in performance between the groups is further enlarged by the fact that in N/Sh groups the random ordering of the trials bounds the maximal extent to which the context predicts the US in comparison to the prediction following the blocked presentations in N+Sh groups, thus limiting context competition. In other words, a further source of difference between the groups is due to context-mediated inhibitory learning between the CS and the US. This experiment exemplifies that mediated learning in the DDA model can predict a negative (inhibitory) relationship between a cued neutral stimulus and a reward, in an arrangement that is formally equivalent to that of mediated conditioning, which in principle relies on a positive correlation between the events. Moreover, it does so with the context, which bears low attentional load, acting as the retriever.

#### Experiment 5: Perceptual Learning

Perceptual learning (PL) is an effect whereby exposure of stimuli reduces generalization between them or, equivalently, improves subsequent discrimination. The phenomenon is highly relevant in learning theory because it is in apparent conflict with latent inhibition: Whereas stimulus exposure delays acquisition, it also facilitates discrimination learning. Amelioration in discrimination has been found to be more profound when the preexposed cues are intermixed (strictly alternated), as compared to being presented in blocks of trials (e.g., Hall & Honey, 1989; Mackintosh, Kaye, & Bennett, 1991; Mondragón & Murphy, 2010; Symonds & Hall, 1995).

Blair & Hall (2003), Experiment 1a, employed a within-subjects design to further control for a differential effect of common stimulus features in assessing the influence of the schedule of exposure. Their experiment, in a flavor aversion preparation, tested PL in a generalization test. In Phase 1, rats received non-reinforced exposure to three flavors, compound stimuli AX, BX, and CX. The first half of trials consisted of alternated presentations of AX and BX, followed by a block of CX trials in the second half. This schedule was counterbalanced across animals, such that half of the animals experienced first a block of CX and then the alternated AX and BX trials. In Phase 2, AX trials were followed by a LiCl injection to induce flavor aversion to AX. In the test phase thereafter (Figure 16, left panel), consumption of BX was higher than consumption of CX, implying that the aversive learning to AX generalized more to CX than to BX, that is, that the animals discriminated better between the alternated stimuli AX and BX than between AX and the blocked CX. The complete design is displayed in Table 3.

**Figure 16:**
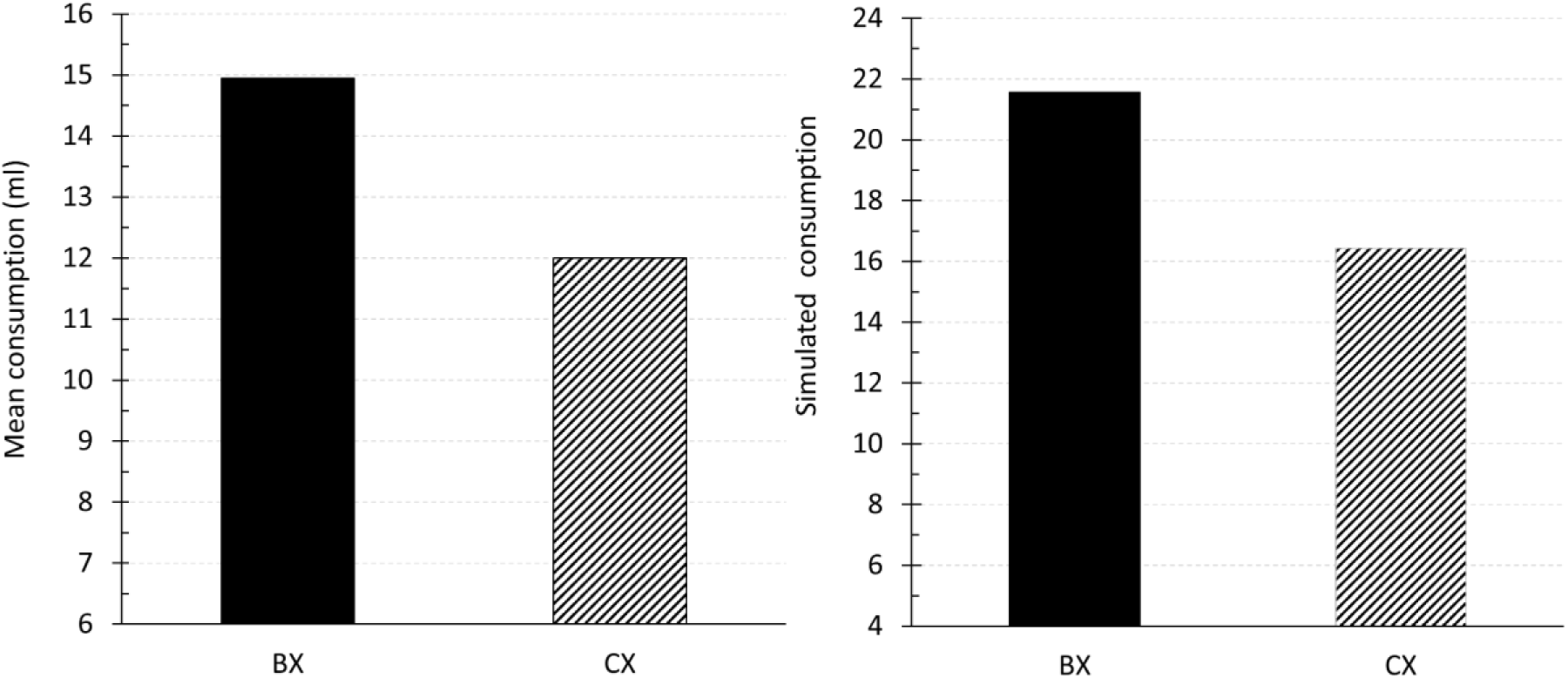
Within-subjects perceptual learning effect. Empirical (original measurement units) and simulated results of Experiment 1a in Blair and Hall (2003). The left panel is an adaptation of the published data showing mean fluid consumption. The right panel displays the corresponding simulated mean consumption for BX and CX test trials.

A simulation of this experiment was conducted with the following temporal parameters: 5 time-units compound CSs, 1 time-unit US (intensity 1), and 75 time-units ITI. All other parameters are shown in Table 2. During Phase 1, each compound was presented 10 times in accordance with the experimental schedule. Conditioning in Phase 2 consisted of 10 trials. In Phase 3, 10 generalization trials to BX randomly interspersed with 2 CX trials were programmed.

The simulated results (Figure 16, right panel) reproduced the empirical data. The simulated consumption (calculated by Equation 15) was higher on BX than CX test trials. Consequently, the model was able to predict that intermixed exposure facilitates discrimination when compared to equally exposed blocked presentations.

According to the DDA model, intermixed AX and BX preexposure results in: 1) slightly higher associability in C than B, such that both neutral associations and subsequent reinforcer associations develop at a higher rate; and 2) a stronger A→C association than the A→B association. Thus, during the AX acquisition trials, mediated conditioning to B will be weaker than to C. Lastly and importantly, intermixed presentations lead to weaker B→A than C→A links. Thus, during the test phase, the associative chain B→A→US is weaker in comparison to C→A→US, contributing towards the disparity between the intermixed and blocked conditions.

### Mediated Learning

A key feature of the DDA model is its ability to account for the way associations change between cues when one or both may be associatively retrieved yet physically absent. It accomplishes this through both its fully connected network and its dynamic asymptote. The latter works on the assumption that the similarity in the level of activation of the predictor and predicted cue dictates the maximal strength of the association that will form between them. Accordingly, a discrepancy in the levels of activity of the stimuli limits their ability to enter into association. This simple idea of adaptability of the asymptote of learning is critical in explaining the apparent contradictory results of mediated related effects that have given origin to conflicting models.

For instance, in a retrospective revaluation experimental setting, two stimuli, a paired-CS and a target-CS, undergo reinforced training in compound. After this training is completed, the paired-CS is further trained (either reinforced or non-reinforced) independently. As a consequence of this treatment, the associative strength of the paired-CS is adjusted, but more importantly, the strength of the target stimulus that does not receive further training is also modified.

Backward blocking and unovershadowing (often referred to as retrospective revaluation) are exemplary cases of mediated phenomena. In a backward blocking procedure, following reinforced compound training, the paired cue is subsequently trained with the same reinforcer. As a result, the target cue losses some of its initial strength. In an unovershadowing design, the paired cue is presented in extinction instead. Following the extinction trials, the associative strength of the non-present target is found to increase when subsequently tested. At a face value, these results seem to suggest an inverse (or inhibitory) relationship between active and retrieved cues.

Formally identical treatments such as sensory preconditioning (SPC) and mediated extinction (ME), in which the initial compound training is giving in the absence of a reinforcer, have produced results that oppose the hypothesis above. That is, they suggest a direct or excitatory connection between retrieved and present cues. For instance, in SPC subsequent reinforced training of the paired cue results in the target acquiring associative strength, rather than losing it as in the backward blocking procedure. In a ME procedure, following non-reinforced training of a compound pair-target, the target is conditioned. In a subsequent phase, the paired stimulus receives extinction training. When the target stimulus is next tested, a reduction in strength compared to that attained earlier is observed. This result contradicts the predictions of theoretical approaches that are able to account for retrospective revaluation experiments, unovershadowing designs in particular, in which an increase in strength is obtained instead.

#### Experiment 6: Unovershadowing and Backward Blocking

In Experiment 3 (Le Pelley & McLaren, 2001) mediated learning effects were studied in a causal judgement task with human participants, and comprised of a series of mediated learning conditions and controls (see Table 3 for details). All participants received the whole set of conditions. In condition A2-A2, Phase 1 consisted of non-reinforced presentations of compound AB, followed by reinforced presentations of C. In Phase 2 the subjects received non-reinforced presentations of a compound AC. In condition A2-A1, Phase 1 consisted of non-reinforced presentations of a compound DE followed by nonreinforced F presentations. Phase 2 consisted of reinforced DF presentations. In condition Control non-reinforced presentations of a compound GH were followed by reinforced presentations of I in Phase 1, whereas in Phase 2 non-reinforced GJ trials were given. The Unovershadowing condition consisted of reinforced KL presentations in Phase 1 followed by non-reinforced presentations of K in Phase 2. In the Backward Blocking condition reinforced MN presentations were given in Phase 1, and Phase 2 consisted of reinforced presentations of M. In the RR Control condition subjects received reinforced OP trials in Phase 1, and O was partially reinforced in Phase 2. In condition Fillers, Phase 1 consisted of non-reinforced Q trials, and reinforced QR trials were given in Phase 2.

The most interesting results of this experiment (see Figure 17, left panel) show that responding to the target cue in the Unovershadowing condition L was higher than that of the Backward Blocking condition N. The difference in ratings between L and P, indicative of unovershadowing, was larger than that between N and P. However, no differences were found in the ratings between P and N, failing to replicate backward blocking, found in their previous experiments. Additionally, no differences were found either between the target cues in the A2-A2, the A2-A1 and the Control conditions (cues B, E, and H respectively) showing no evidence of mediated learning. These results pose a challenge to models such as Holland (1983) and SOP, which cannot predict unovershadowing or backward blocking. These can be considered as post-acquisition effects resulting from memory interference rather than new learning (McLaren, 1993).

**Figure 17:**
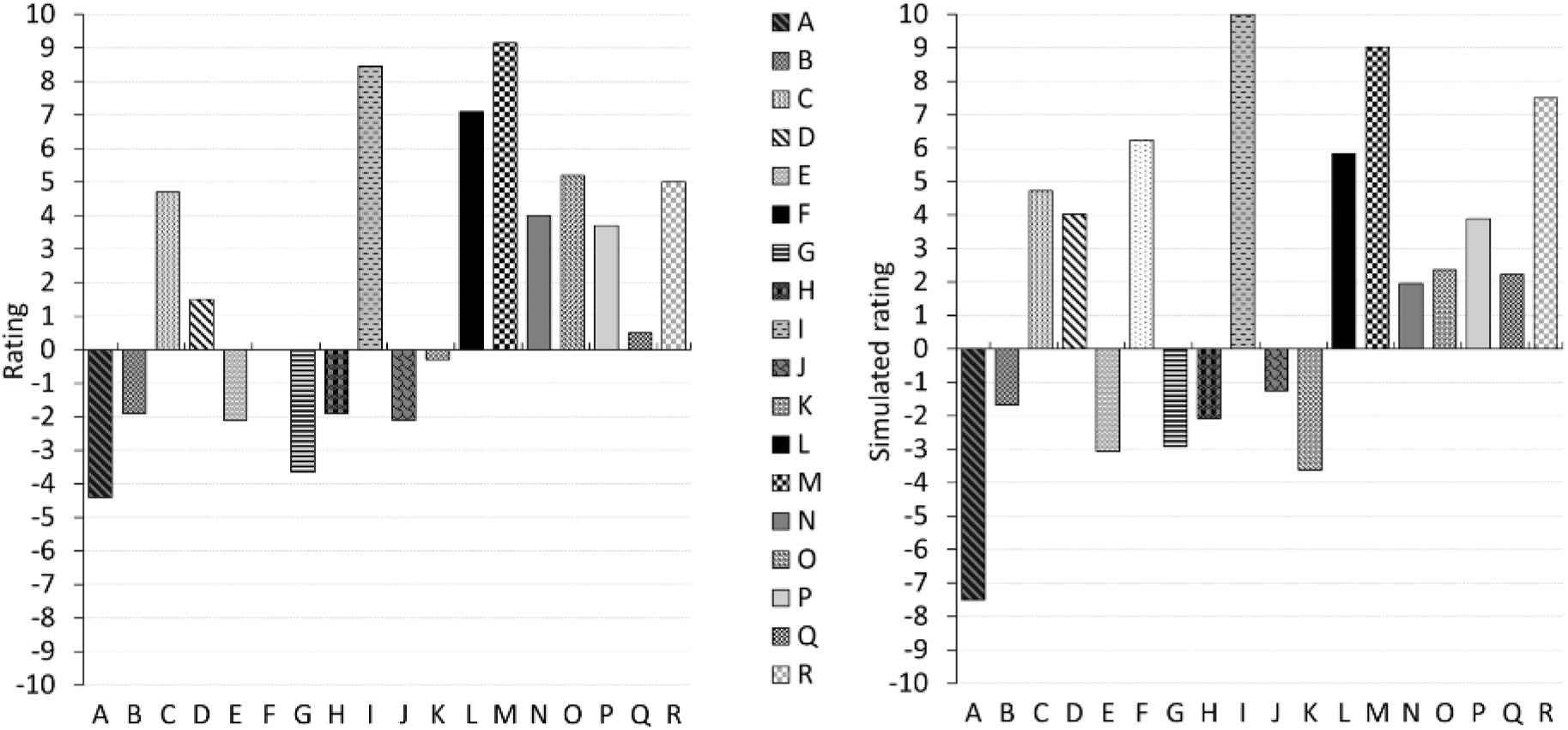
Unovershadowing and backward blocking. Empirical (original measurement units) and simulated results of Experiment 3 in Le Pelley and McLaren (2001). The left panel is an adaptation of the published data showing mean ratings in a food allergic reaction task during test. Positive ratings indicate high likelihood of a given food resulting in an allergic reaction. Negative ratings indicate that the food would prevent allergic reaction. The right panel displays the corresponding simulated ratings for each cue.

The temporal parameters of the simulation used a 1 time-unit CS, 1 time-unit US (intensity 1), and 20 time-units ITI. All other parameters were set as per Table 2. Phase 1 of the experiment was programmed to consist of 8 trials of each trial-type. Phase 2 was conducted with 8 trials for each condition. In Phase 3 each cue was tested in a single trial in a random order. To calculate simulated ratings, the weights were normalized such that the highest weight corresponded to a rating of 10.

The simulations matched the empirical data closely (Figure 17, right panel). Crucially, simulated ratings to the Unovershadowing target cue L were higher than those observed for the Backward Blocking condition N. Simulated data showed differences in the ratings between P and N, thus exhibiting backward blocking, and between the target cues in the A2-A2, the A2-A1 and the Control conditions (cues B, E, and H, respectively) which were however rated negatively as in the published experiment.

The DDA model predicts that the associative links of cues B, E and H have such low ratings towards the outcome, because during Phase 1 they are presented in the context of reinforced trials of other conditions, resulting in the context acquiring a mild excitatory strength thus fostering inhibition to B, E and H, therefore counteracting mediated acquisition. Furthermore, the low amount of trials in Phase 1 does not allow for significant within-compound links to form between the CS compounds, which impedes Phase 2 mediated learning by leading to a lower level of retrieval of the target cue.

In the Unovershadowing Condition, reinforcement of K and L is followed by nonreinforcement of K. This leads to excitatory revaluation of the L link, as the relative probability of K being a predictor of the outcome diminishes. Specifically, the links K→L and K→+ from Phase 1 of the experiment result in K retrieving L and the US to a similar level of activation in Phase 2. As the K→+ link extinguishes over these presentations, and as the asymptote in the model supports a high level of learning from L to the US due to their similar level of activity, the net result is a growth of the link L→+. In the case of Backward Blocking, a similar revaluation takes place, but in the opposite direction. The link M→N that forms in Phase 1 of the design leads to M retrieving N in Phase 2. As the asymptote of learning in the model between the retrieved N and the present US would be lower than when both cues were present in Phase 1 (due to the disparity in the level of activations), this would result in a decrease of the strength of the association between N and the US.

Further, since this decrease is dependent on how strongly N is retrieved by M, as well as on the extent to which N contributes towards predicting the outcome, the model predicts that a larger number of compound trials in the first phase should result in more backward blocking being observed.

#### Experiments 7a and 7b: Backward Sensory Preconditioning (BSP) and Mediated Extinction (ME)

In SPC, pre-training of a target-paired stimulus compound (XA) is followed by conditioning being given to the paired stimulus (A) and by a subsequent test of the strength of the target (X). With a preparation of aversive conditioning in rats, Experiment 2 (Ward-Robinson & Hall, 1996) aimed to uncover the mechanism underlying SPC, that is, whether sensory preconditioning operated by means of an associative chain X→A→US or whether a direct link X→US was formed through mediated learning. To do so, they employed backward serial presentations of the compound stimuli during the initial training, that is A→X, with the intent of preventing the chain of association from occurring. Simulating the result of this experiment is an important validator of the capability of the DDA model to reproduce conflicting mediated phenomena. The model postulates that although weak, an excitatory link would indeed be formed between X and A during backward conditioning. Thus, during test, X would still be able to associatively activate the chain X→A→US. However, during conditioning of A, the target X would also be active, and mediated conditioning A→US would occur. Despite that the DDA model advances two sources able to produce the effect, mediated conditioning would nonetheless be weak given that the discrepancy between the levels of activation of the stimuli involved would bear a low asymptote of learning. The conjunction of both mechanisms would therefore be needed to produce the effect.

In Experiment 2 (Ward-Robinson & Hall, 1996) two groups of rats received random presentations of two serial compound trial-types, A followed by X, and B followed by Y in Phase 1. In Phase 2, presentations of A were followed by an outcome whereas presentations of B were not. Thus, B and its paired Y acted as a within-subjects control for BSP. In Phase 3, Group Ext received non-reinforced presentations of each A and B, intended to extinguish their association between A and the US, while Group VI received none. The test phase, Phase 4, consisted of X and Y trials for both groups (see Table 3). If the association between A and the US were critical to the effect, BSP would only to be expected in Group VI.

Results (Figure 18, left panel) showed that in Group VI, suppression to X, paired with A, which was reinforced in Phase 2, was greater than to Y, paired with B, which in turn was presented without reinforcement. That is, a BSP effect was shown in Group VI but not in Group Ext, for which extinction training following conditioning was given. In this group, suppression to X did not reliably differ, indicating that the effect in Group VI relied on the strength of the association between A and the reinforcer. In other words, the extinction treatment to A abolished the BSP effect as observed in Group VI.

**Figure 18:**
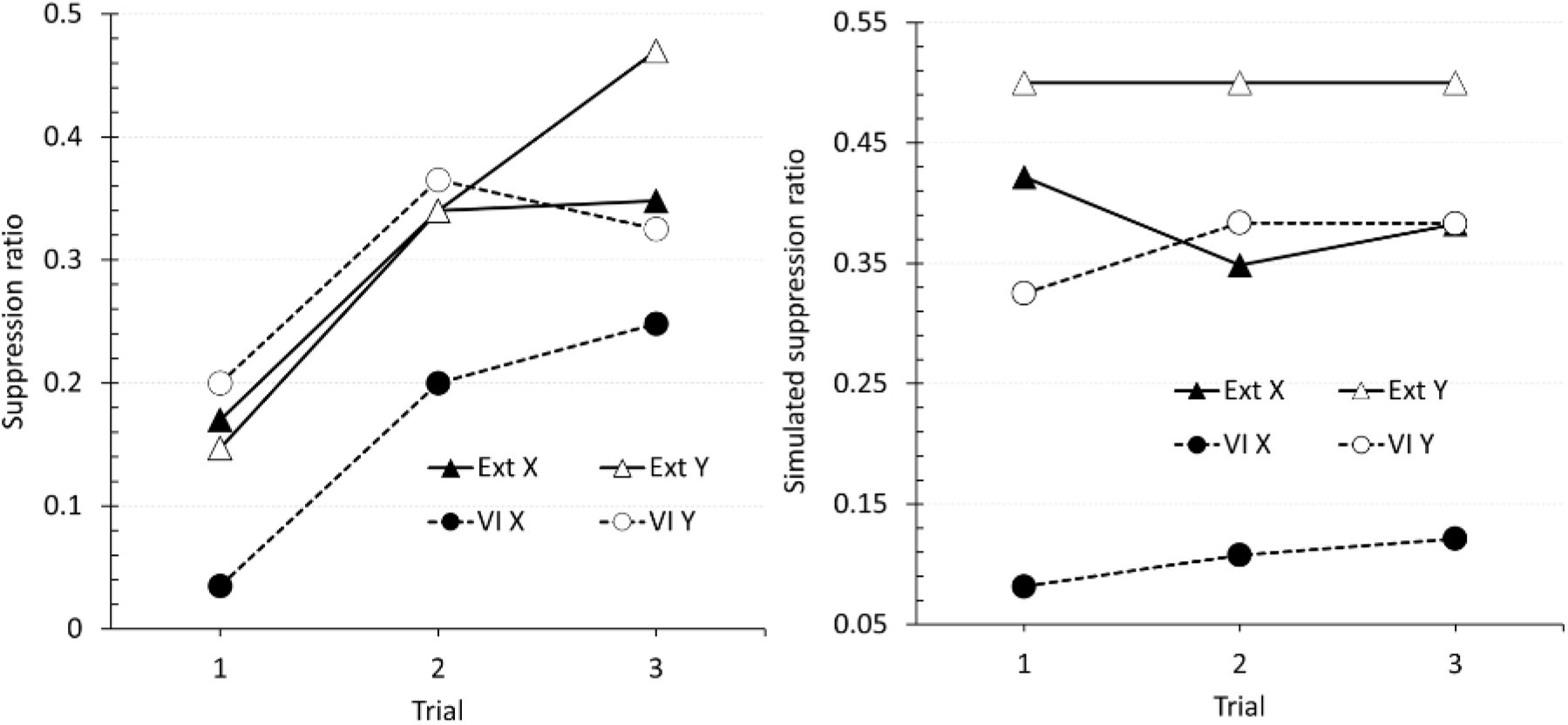
Backward sensory preconditioning. Empirical (original measurement units) and simulated results of Experiment 2 in Ward-Robinson and Hall (1996). The left panel is an adaptation of the published data showing mean suppression ratios to X and Y during the test phase for Group VI and Group Ext. The right panel displays the corresponding simulated suppression ratios.

The temporal parameters used for this simulation were a 5 time-units CS, 1 time-unit US (intensity 1), and 50 time-units ITI, with remaining parameters presented in Table 2. In Phase 1, all groups were programmed to receive 12 random presentations of serial-compounds A→X and B→Y. In Phase 2, both groups received 4 presentations each of trace conditioning A→+ and non-reinforced B trials, presented in the following order MNNM. In Phase 3, Group Ext received 44 random non-reinforced presentations of cues A and B, while Group VI was programmed to receive no training. Finally, both groups were programmed to receive 3 random non-reinforced presentations of cues X and Y in Phase 4.

Simulated results (Figure 18, right panel) replicated the empirical pattern of responses. In the test phase, suppression to X in Group VI was greater than suppression to Y. However, unlike the empirical results, a smaller but clear difference between X and Y was also present in Group Ext. This difference could just be attributed to a more complete extinction of the association between A and the US during the simulated Phase 3 in comparison to the observed empirical levels, in conjunction to a lesser stimulus generalization. It is worth noticing that, despite not being significant, a similar tendency can be observed in the empirical data in the last test trial, after further extinction occurred.

Ward-Robinson and Hall’s Experiment 3 (Ward-Robinson & Hall, 1996) tested (forward) mediated extinction in a within-subjects design that paralleled the one above (see Table 2). Animals received serial A→X and B→Y trials in Phase 1, followed by conditioning to X in Phase 2 in which Y was also presented but in extinction. Phase 3 consisted of presentations of A in extinction and finally, during Phase 4, test trials to X and Y were given.

If mediated extinction occurred during Phase 3, conditioning to X associatively retrieved by A should be reduced in comparison to that of Y. Results of this experiment (Figure 19, left panel) showed that indeed this was the case. During test, suppression to X was lower than to Y.

**Figure 19:**
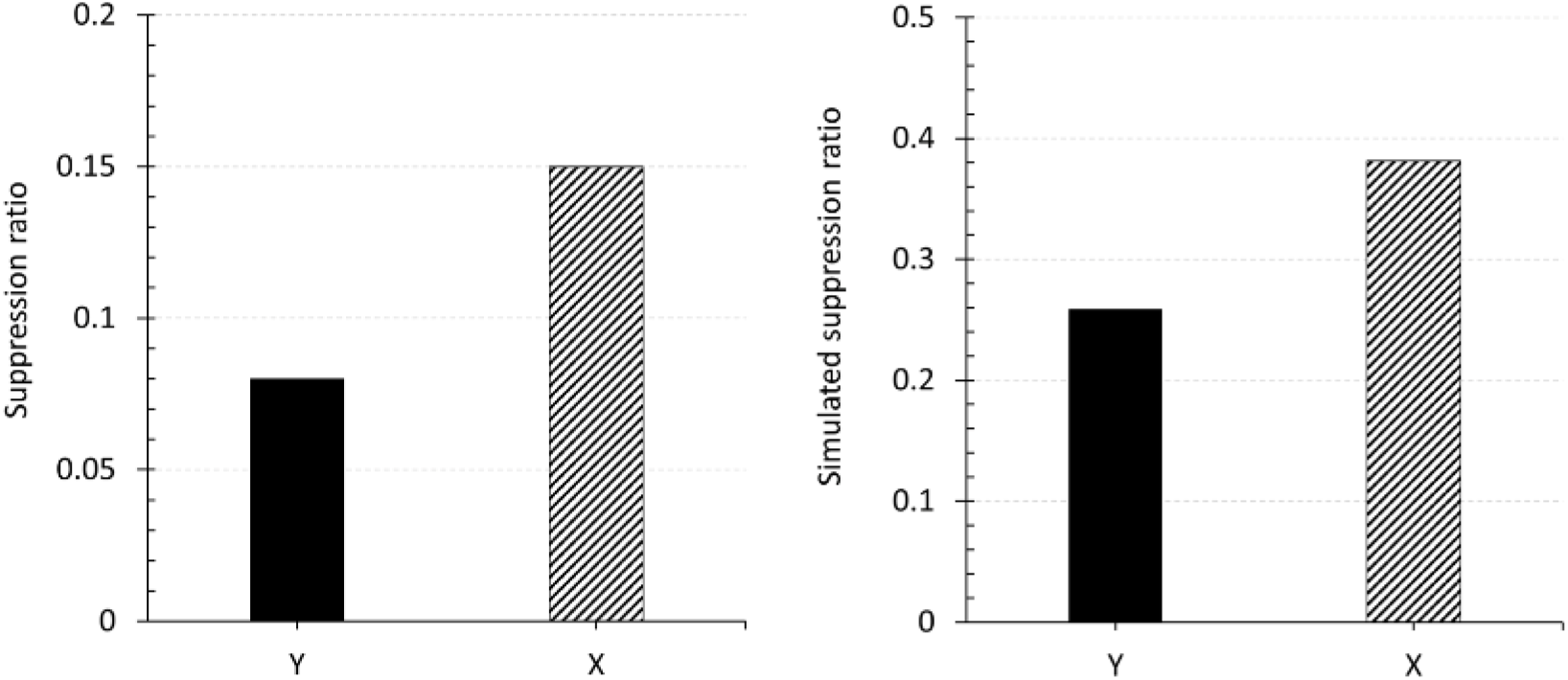
Mediated extinction. Empirical (original measurement units) and simulated results of Experiment 3 in Ward-Robinson and Hall (1996). The left panel is an adaptation of the published data showing mean suppression ratios to X and Y during the test phase. The right panel displays the corresponding simulated suppression ratios.

The parameters and design used in this simulation are displayed in Table 2. The simulation used a US intensity of 2. During Phase 1, animals received identical training to those in Experiment 7a. In Phase 2 however, X and Y (rather than A and B) were both reinforced. In Phase 3, A was extinguished and responding to X and Y tested in Phase 4.

The simulated results showed mediated extinction of X, that is, a loss of the associative strength of X as a consequence of extinguishing its paired stimulus A, when compared to the strength attained by Y whose paired stimulus B did not undergo extinction (Figure 19, right panel).

The DDA model predicts that in Experiment 7a, presentations of A→X and B→Y in Phase 1 would result in bidirectional excitatory learning between the two CSs of the serial compound, as the CS that is presented first is presumed to leave a persistent memory trace even after its offset and excitation occurs whenever two similarly active elements co-occur. In the subsequent conditioning to A in Phase 2, A retrieves X. Although the lesser activation of X implies a lower asymptote of learning towards the present US, this nevertheless increases the associative strength of X. The A extinction trials in Group Ext lead to: (1) a decrease in the association between A and X; (2) a reduction in the association between A and the reinforcer; and (3) a mediated loss in the US expectation elicited by X, as A retrieves the US more strongly than X. This discrepancy in the levels of activation would foster the development of inhibitory learning between the weakly retrieved X and the strongly retrieved US on the basis of the dynamic conceptualization of the asymptote in the DDA model. The absence of extinction trials in Group VI would preclude the effects described above from occurring, and thus X would still sustain the level of prediction of the US attained during conditioning to A. However, since the DDA model assumes that bidirectional links will be formed despite the serial stimulus presentation, the contribution of the associative chain X→A→US cannot be diminished. Nonetheless, the extinction treatment in Group Ext would prevent A eliciting the US representation, thus countering the BSP effect. Further, if the model is accurate, it would imply that should the first and second phases of the experiment be respectively longer and shorter, it could in fact reverse the effect such that mediated conditioning would take place in Group Ext between X and the US. This would occur as elongating the first phase would strengthen the A→X link to a level rivalling the more speedily acquired A→US association, while shortening the second phase would achieve a similar result by weakening the A→US association. This would subsequently lead to X and the US being retrieved to an equally active level on the A-trials, in which case the dynamic asymptote of the model would predict a strong excitatory association from X to the US. The mediated extinction hypothesis was further validated by the results of Experiment 7b. The experiment consisted of an equivalent Phase 1 treatment as experiment 7a, and hence the model predicts equivalent X→A and Y→B links forming in this phase. During the second phase, the DDA model predicts that, due to the aforesaid links, both X and Y would strongly retrieve a representation of A and B respectively. The high level of activation of these representations would produce a high asymptote of learning from these cues to the present US, and thus would undergo mediated conditioning. During the crucial third phase, nonreinforced presentations of A would retrieve X and the US, but the retrieved representation of the US would be weaker than that of X (reflecting the link strength from A to these cues). Therefore, the asymptote of X toward the US would be smaller than its prediction, and hence the error of X would become negative, resulting in X losing strength (mediated extinction). Additionally, the associative chain X→A→US would weaken due to extinction of the A→US link in Phase 3. Hence, in the test phase, X would elicit less responding than Y, which did not undergo such mediated extinction.

### Non-linear Discriminations

Non-linear discriminations are those in which a linear summation of the associative strengths of the constituent CSs is insufficient to accurately predict when reinforcement will or will not occur. Solving these discriminations therefore requires the use of additional representational, learning, and attentional processes to introduce non-linearity in the system. Despite its elemental nature, the DDA model is able to solve complex non-linear discriminations primarily through its assumption of elements being able to be activated by multiple stimuli. This effect is accentuated by both the alpha and the predictor error term of the model. The former leads to elements active on reinforced trials having higher associability. Further, the persistently high error, produced by partial reinforcement, eventually triggers a decay mechanism of the revaluation alpha that disproportionately affects shared elements. The latter operates through the network of neutral element associations to reduce the learning rate of cues that are predicted well by other cues, thereby allowing stimuli that are predictive of reinforcement or non-reinforcement to learn faster.

#### Experiment 8: Negative Patterning (NP)

Whitlow and Wagner’s (1972) Experiment 1, studied a NP discrimination in eyeblink conditioning (Table 3). In Phase 1, rabbits were presented with reinforced presentations of A, B, and C. Phase 2 consisted of random presentations of reinforced stimuli A and B, as well as the non-reinforced compound AB trials. In Phase 3, training continued as in Phase 2 with the addition of reinforced C trials. Finally, in Phase 4 isolated stimulus presentations and all combinations of the three cues were tested in a random order, that is it consisted of A, B, C, AB, AC, and BC presented in extinction. Stimulus C was used to assess the contribution of simple summation in the responding elicited by a compound in comparison to putative configural components emerging from training a stimulus compound.

Results of this experiment showed that the animals learned to discriminate between the stimuli and the compound AB (Figure 20, left panel), withholding responding on AB trials. Also, test showed that this suppression of the response was only evident for AB trials, that is, the animal’s response to the other compounds tested, i.e., AC and BC was sustained (Figure 21, left panel).

**Figure 20:**
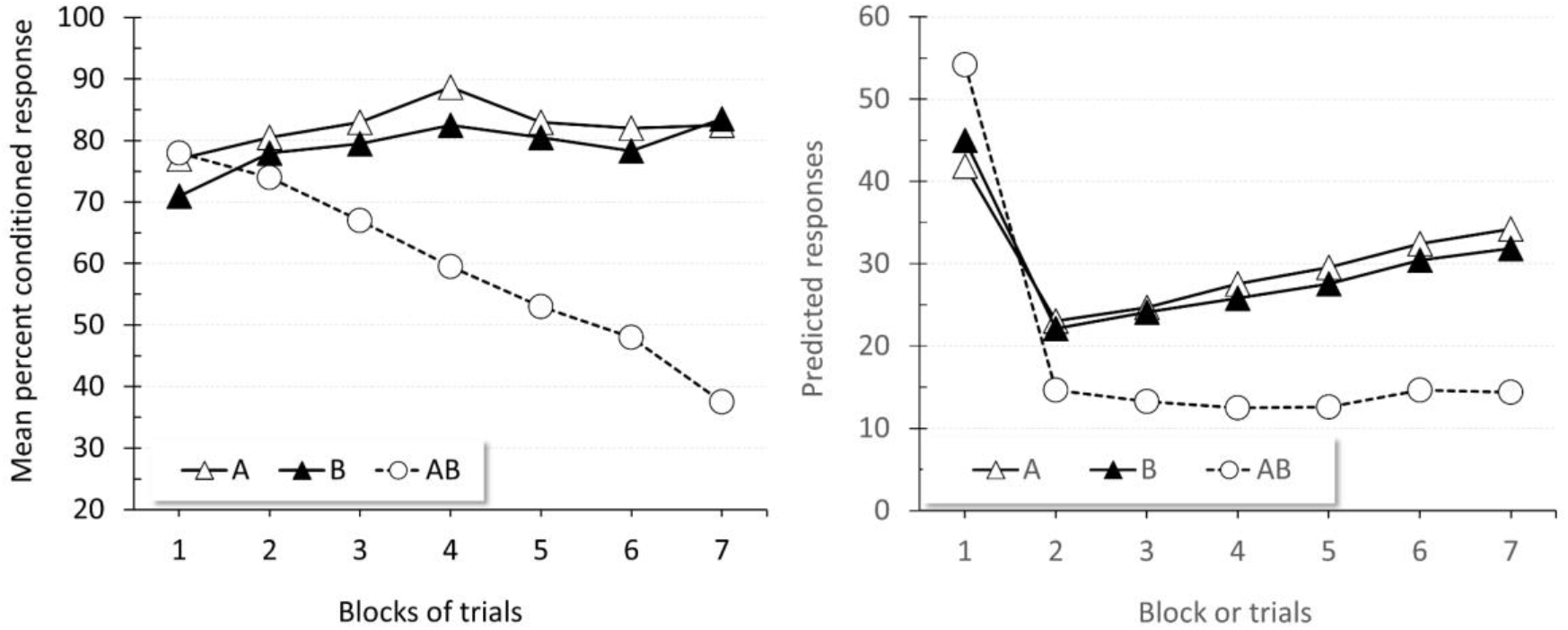
Negative patterning acquisition. Empirical (original measurement units) and simulated results of Experiment 1 in Whitlow and Wagner (1972). The left panel is an adaptation of the published data showing responses to A, B, and AB during a negative patterning discrimination. The right panel displays the corresponding simulated responses.

**Figure 21:**
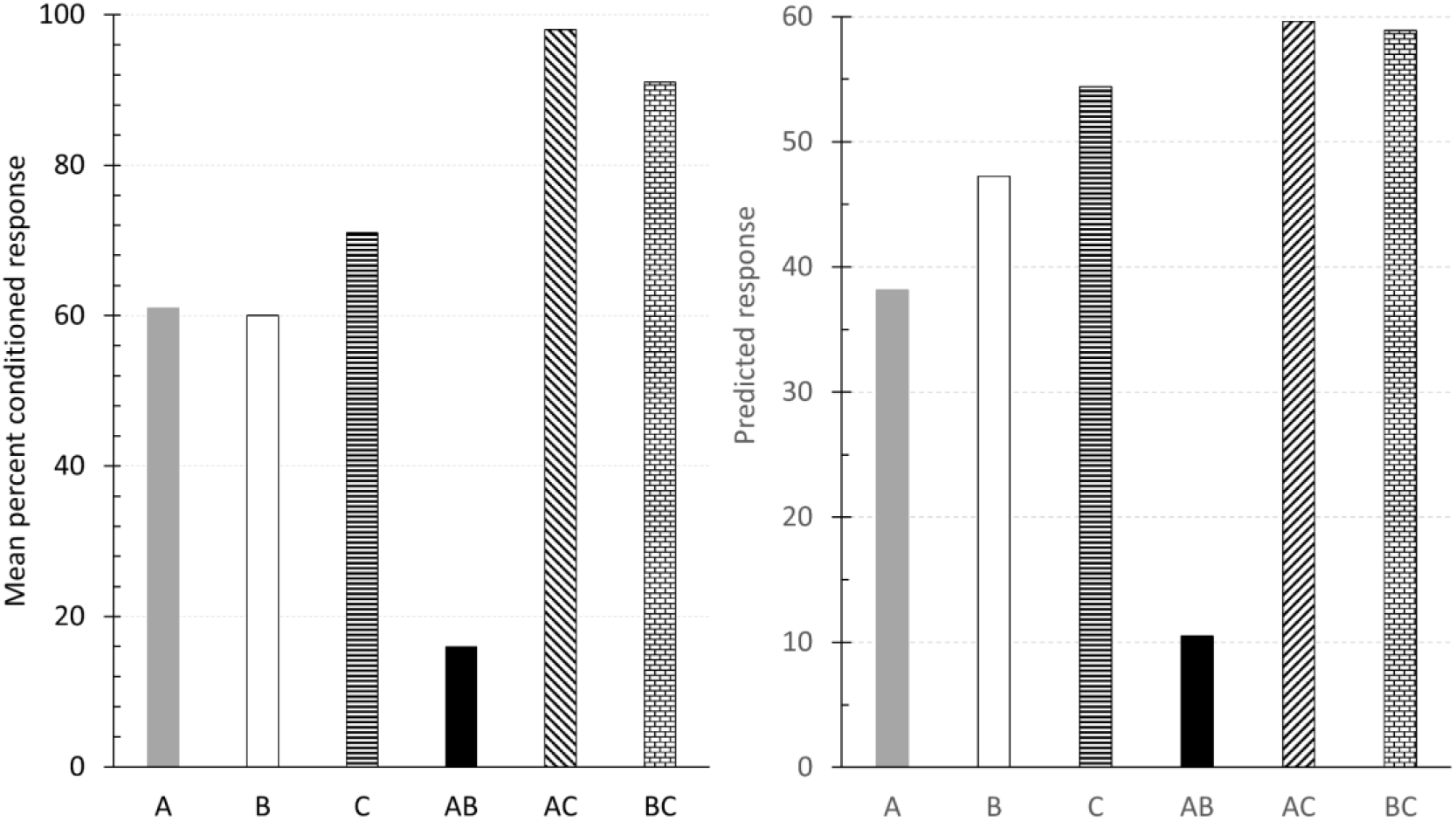
Negative patterning test of the contribution of simple summation. Empirical (original measurement units) and simulated results of Experiment 1 in Whitlow and Wagner (1972). The left panel is an adaptation of the published data showing test data following a negative patterning discrimination. The right panel displays the corresponding simulated responses.

The temporal parameters for this simulation were a 1 time-unit CS, 1 time-unit US (intensity 0.5), and 20 time-units ITI. The parameters of the experiment are found in Table 2. Phase 1 consisted of 180 reinforced trials of A, B, and C presented in a random manner. Phase 2 consisted of random 560 presentations of reinforced A and B trials and 1,120 nonreinforced AB trials. In Phase 3, 40 reinforced random presentations of A and B were programmed, together with 20 reinforced presentations of C, and 80 non-reinforced presentations of the compound AB. Finally, the test phase consisted of a single, nonreinforced presentation of trial-types A, B, C, AB, AC, and BC.

The simulation visibly reproduced the experimental data. Figure 20, right panel, shows the acquisition of the NP discrimination. Following an initial drop due to the fluctuation of responding across the first block of 80 reinforced and 160 non-reinforced trials, responding to stimuli A and B progressively increased across trials, while responding to AB remained steadily low. The right panel of Figure 21 displays the test results. These parallel the empirical data, that is, suppression of responding to AB was larger than to all other stimuli or compounds.

According to the DDA model, during discrimination training redundant cues (the elements shared between A and B as well as the context) become excitatory toward the US, while elements unique to A or B become highly inhibitory. Moreover, on AB trials the shared elements between A and B can only be sampled once. Thus, they contribute relatively less to responding than in A or B trials, thereby leading to a lower response being elicited on such trials by these elements than would be expected by linear summation. Two unique mechanisms of the DDA model strengthen this effect. First, unique elements of A and B in AB trials strongly predict the shared elements, thereby delaying the speed of learning by decreasing their associability. This would contribute to sustain a high expectation for the reinforcer thus leading to a high negative error term for the unique elements. Secondly, the associability rate to the reinforcer, which is initially sustained due to the inconsistent outcome, is revaluated. The DDA model predicts that when training under inconsistent outcomes is prolonged (e.g., A being reinforced and non-reinforced through training) and the error term remains high, animals come to learn that the outcome is inconsistently attained and a mechanism to cease sustained attention is activated. This mechanism would become active sooner for the shared elements than for unique elements because these elements are active in more trials, therefore accruing evidence faster for the contingency being inherently random. As a consequence, the associability of the shared elements is further reduced, pushing the unique elements to become more inhibitory. The net effect is that in compound trials, the inhibitory strength of the unique A and B elements is of higher magnitude than the excitatory strength of shared elements, thereby leading to a with-holding of response. When the stimuli are experienced in isolation, the contribution of the inhibitory strength of the unique elements is reduced by half, yet the same quantity of excitatory strength is provided by the shared elements. Hence, generating a higher net reinforcer prediction, and thus a higher rate of responding.

#### Experiment 9: Biconditional Discrimination

In a biconditional discrimination four compounds of two stimuli are reinforced in a manner such that each individual cue receives both reinforced and non-reinforced training (AB+, CD+, AD-, BD-). This discrimination, from the elemental perspective, is difficult to solve due to each stimulus receiving equal partial reinforcement. It therefore seems to require the existence of configural nodes, which enable the discrimination to be solved through nonlinear means. However, the assumption of pairwise shared elements in the stimulus representation in the DDA model, a mild assumption, circumvents this difficulty, enhancing the power of this and other elemental models to predict ‘configural’ learning without the assumption of configural representations.

To prove it, we simulated the biconditional discrimination experiment in (Lober & Lachnit, 2002), which used an aversive skin-conductance conditioning procedure in human participants. For the biconditional treatment, participants received random presentations of reinforced (AB+, CD+) and non-reinforced (AD-, BD-) compounds. The participants learned the discrimination, progressively increasing responding on the reinforced trials and withholding responding on non-reinforced trials (Figure 22, left panel).

**Figure 22:**
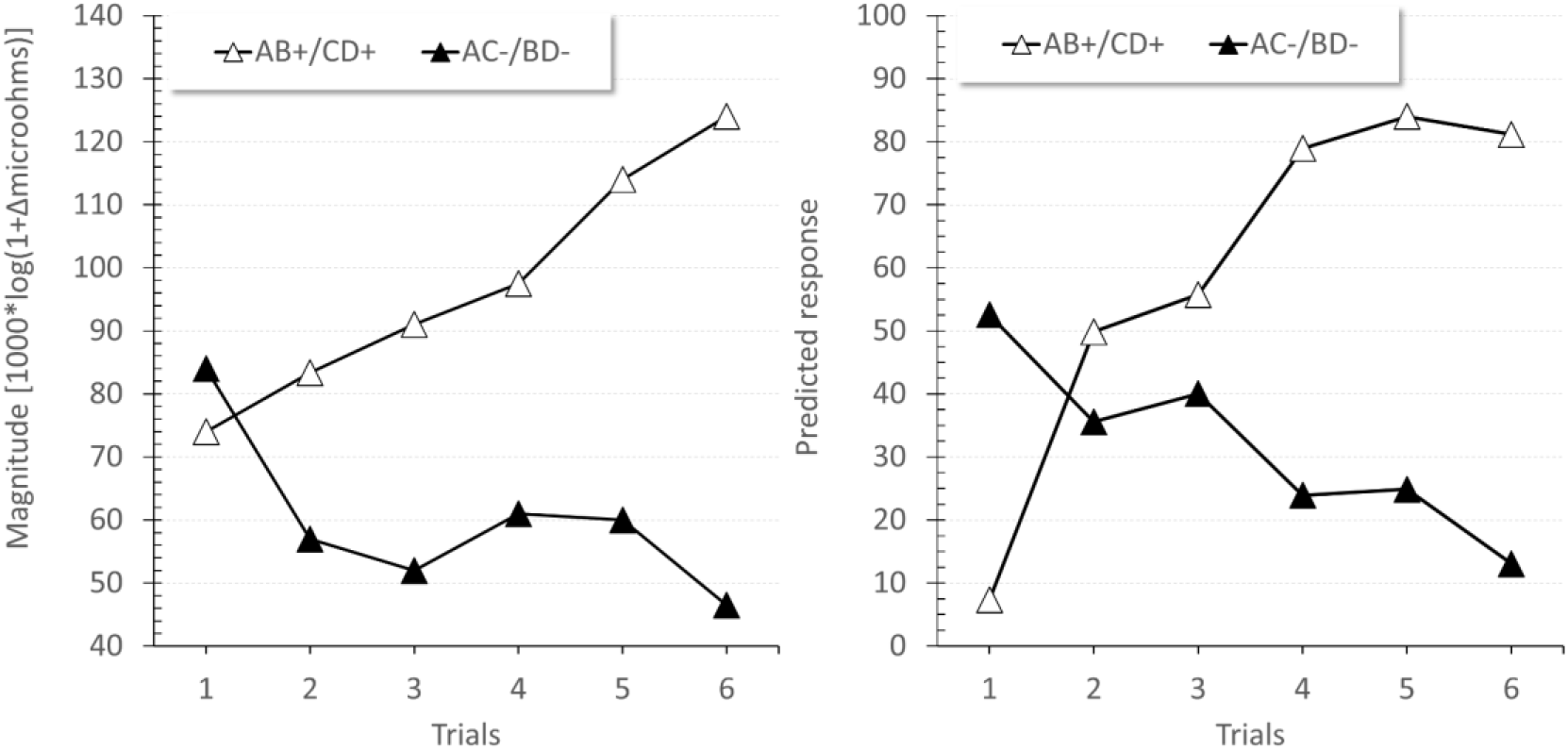
Biconditional discrimination. Empirical (original measurement units) and simulated results of a biconditional experiment in Lober and Lachnit (2002). The left panel is an adaptation of the published data showing second-interval responses (SIR) for combined AB+/CD+ trials and AD-/BC-. The right panel displays the corresponding simulated responses.

The simulation’s temporal parameters consisted of a 1 time-unit CS and US (intensity 2), and a 50 time-units ITI. The remaining parameter values are presented in Table 2. Training consisted of 50 randomly presented trials of each reinforced compounds AB+ and CD+, and of the non-reinforced compounds AC- and BD-. The full design is in Table 3.

The simulation matched the empirical pattern of responding. Figure 22, right panel, shows an initial increase in responding on both reinforced and non-reinforced trial-types. This was followed by a gradual decrease in responding on non-reinforced trials, with responding on reinforced trials remaining high.

The DDA model predicts that initially, the context and the unique elements of the four compound stimuli would gain some excitatory strength. In parallel, the elements shared between the stimuli would also gain modest excitatory strength. However, as training progresses the elements shared between the two pairs of reinforced trials (i.e., the elements shared between A and B and between C and D) would progressively become inhibitory, promoting super-excitation of their unique elements. On reinforced trials, e.g., AB, both A and B would recall their shared elements, but they could only be sampled once and therefore their activation would be comparatively lower than that of the unique elements and the US. Thus, due to the dynamic asymptote mechanism, the discrepancy in the level of activation would preclude the shared elements from gaining excitatory strength. On the next nonreinforced trial, the prediction error would turn negative (as the US is not present but there is a general excitatory prediction), and all elements would lose strength, but because these shared elements would have weaker strengths than other elements, they would reach negative values earlier. At that point, they would foster acquisition of the unique elements excitatory strength on subsequent reinforced trials (by increasing the prediction error). On the next nonreinforced trial, these boosted excitatory elements would produce a larger negative prediction error, engendering more inhibition in the shared elements. This cycle would recur, adjusting the strengths through training.

Given that in each reinforced trial (AB and CD) their respective inhibitory shared elements are only sampled once the inhibitory impact would be less significant than in the non-reinforced trials (e.g., AC) in which A would retrieve the elements shared with AB, and C would retrieve the inhibitory elements shared with CD. Thus, duplicating the contribution of the inhibitory strength.

#### Experiment 10: Generalization Decrement Overshadowing vs. External Inhibition

A significant hurdle for configural models of learning is the observation that removing a cue from a previously reinforced compound stimulus (overshadowing test) produces a greater decrement in learning than when a novel cue is added to the compound (external inhibition test). As an elemental model, the DDA model inherits an advantage in explaining this effect: Training a stimulus compound, e.g., AB, and then testing a single stimulus (e.g., A) should lead to a strong generalization decrement because the summation rule would imply that the total associative strength of the compound is distributed in the individual stimulus strength. Thus, testing a single stimulus removes one source of predictive value and necessarily reduces the total prediction. In contrast, training a single stimulus A implies that the total amount of available prediction is accrued by A. Consequently, testing a compound AB should result in no decrement, provided that B has no inhibitory value itself.

Precisely this difference between external inhibition and overshadowing was studied in Brandon *et al.* (2000), using a rabbit eyeblink conditioning procedure. In their experiment, Group A, Group AB, and Group ABC received respectively reinforced A, AB, and ABC trials. Subsequently, all groups received test trials of randomly presented reinforced and non-reinforced presentations of A, AB, and ABC (see Table 3). The experiment found that adding a cue to a previously reinforced compound produced a smaller decrement in responding than removing a cue from a previously reinforced compound. As it is apparent in the left panel of Figure 23, the decrement of generalization produced by adding one or two cues (AB and ABC) in Group A was comparatively smaller, although clearly present, than that observed when B was removed in Group AB, and when one or two cues were omitted in Group ABC.

**Figure 23:**
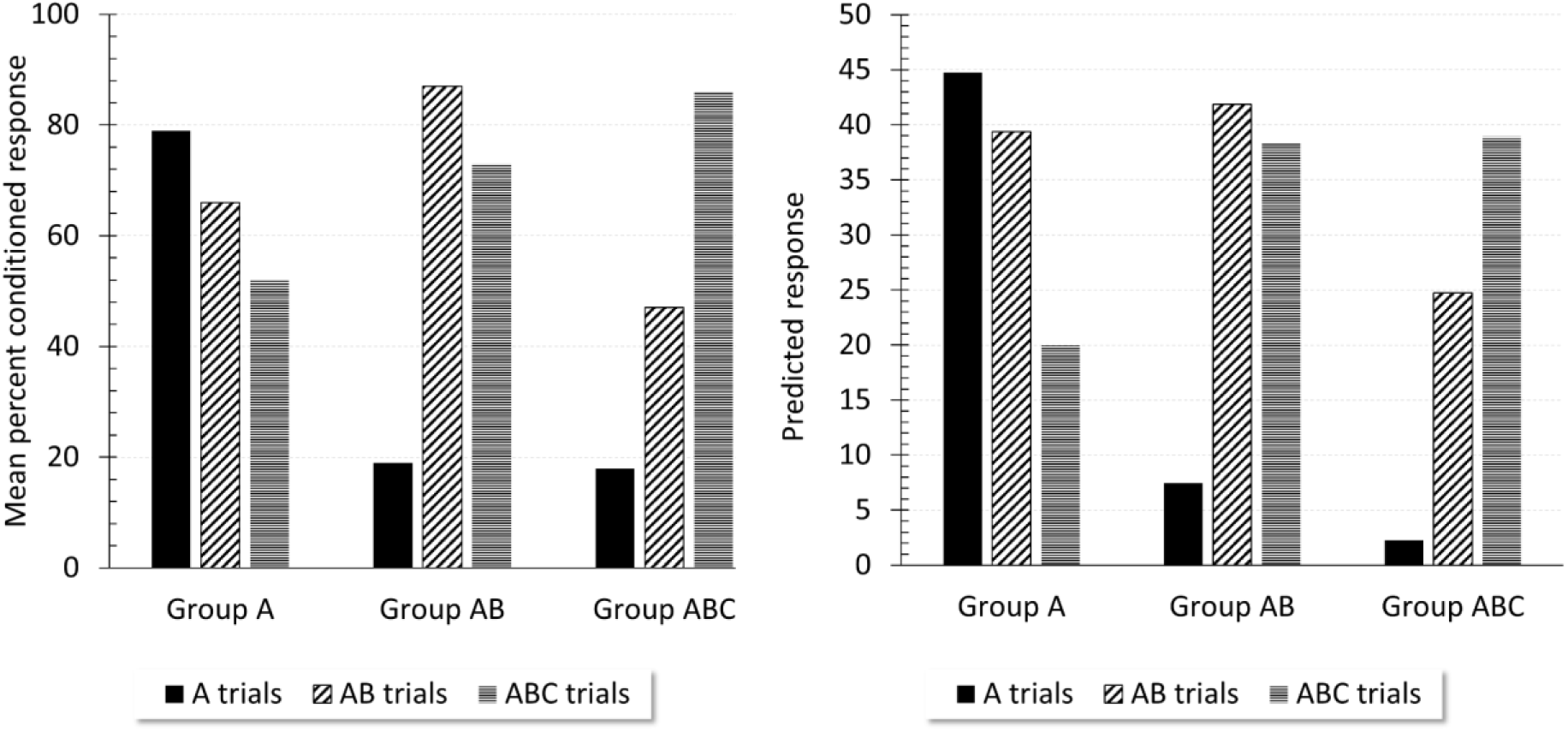
Generalization decrement in overshadowing and external inhibition. Empirical (original measurement units) and simulated results of a generalization decrement experiment in Brandon *et al.* (2000). The left panel is an adaptation of the published data showing responses during test to A, AB, and ABC. The right panel displays the corresponding simulated responses.

The experiment was simulated using the following temporal parameters: a 5 time-units CS, 1 time-unit US (intensity 1), and 50 time-units ITI (Table 2). In Phase 1 groups A, AB, and ABC received 20 programmed reinforced trials of type A, AB, and ABC, respectively. Phase 2 was programmed to present 20 randomly presented trials of types A, AB, and ABC. Each of these trial-types was reinforced on half of the presented trials.

Simulated results are an accurate reproduction of the empirical pattern. That is, the generalization decrement adjusted to the incremental change from the conditioned cue to the testing cue in testing, and this decrement in generalization was larger when the cues were removed than when they were added (Figure 23, right panel).

The DDA model produces such a result due to its elemental ontology leading to the summation assumption of constituent associative strengths of a compound. The effect observed by the addition of extra cues can be ascribed to the presence of shared elements. Since the test was conducted including reinforced trials, the unique features of the stimuli added could have undergone some inhibitory learning, thus reducing the net compound strength.

## GENERAL DISCUSSION

In this paper, we have introduced a real-time computational model of associative learning, the DDA model. Table 4 provides a comparative view of the predicted power of the DDA model against other models of associative learning, as discussed throughout the paper, in relation to relevant phenomena. Its ontology is founded upon a connectionist network, wherein nodes representing the attributes of a stimulus (clusters of elements), whether reinforced or not, enter into association with one another in proportion to their level of activity and are distributed as subsets of shared and unique elements to the stimuli involved. The defining algorithmic characteristics of the DDA model are threefold. First, it extends traditional error correction learning, familiar from models such as RW, TD, and SOP, by modulating learning by a predictor error term. This new error term measures the extent to which a predictor of an outcome is in return itself predicted by other stimuli (e.g., the context). An immediate implication is that more predictable predictors associate at a slower rate than less predictable ones. Secondly, the model conceptualizes a revaluation associability that adjusts independently for motivationally relevant and neutral outcomes. Thirdly, the model introduces a novel variable asymptote of learning that encapsulates more closely the principle of Hebbian learning, by postulating that elements with similar activity levels form stronger associations with one another. Hence the direction of learning between two cues at a given time can differ, in an otherwise equivalent condition, based on whether the prior link strength from one to the other is above or below the momentary level of this asymptote. This process critically allows the DDA model to predict seemingly contradictory mediated learning effects.

The associability of a cue operates by keeping track over a window of time, through a moving average, of uncertainty in the occurrence of a given category of outcome, reinforcement or otherwise, and thence modulating its respective rate of association. Thus, for instance, when the occurrence of reinforcement following a stimulus is uncertain, learning of associations between the latter and the reinforcer proceeds more rapidly. However, the stimulus associability is revaluated in the face of persistent uncertainty and a decay mechanism is triggered under such conditions. This revaluation in the cue associability operates based on whether a slower updating moving average of prediction errors crosses a threshold. Once this threshold is crossed, a decay mechanism kicks in that reduces the associability progressively. Though the Mackintosh model’s attentional rule seems opposed to that used in the DDA model (as the latter resembles the PH alpha), the DDA model revaluated alphas can in fact produce effects whereby the best predictor of an outcome captures the most selective attention, due to it producing the highest US prediction error before the onset of the US. This, however, is dependent upon sufficiently long CSs being used.

Similarly to the McLaren-Mackintosh, SLGK, and SOP models, learning between motivationally neutral cues allows the DDA model to reproduce various ‘silent learning’ and preexposure effects. It predicts these effects as arising from both learning between neutral CSs, and through the constituent elements of a CS forming unitized nodes during preexposure. Thus, the model does not require arbitrary memory states for producing an account of stimulus exposure and it postulates the same learning rules and processes both for reinforced and non-reinforced learning, thereby maintaining parsimony. That is, only one learning rule and one type of element are used in the model, i.e., we maintain that reinforced outcomes should be treated simply as a subset of normal stimuli, as opposed to having a completely unique ontology in terms of their stimulus representation and learning. The intensity of a conditioned response to a motivationally relevant stimulus (or to a neutral one, e.g., an orienting response to a light when predicted by a tone) is simply a function of its associative strength. The model’s integration of its unique alpha conceptualization together with its distinctive second error term results in it accounting for preexposure effects as arising from the expectancy of a cue decreasing due to learning. It hence avoids difficulties faced by other models, such as those derived from proposing that a noUS link forms during preexposure; a suggestion contradicted by evidence denoting US-specificity of inhibition.

The model predicts both context specificity and the sigmoidal shape of latent inhibition, with the latter resulting from the exponential changes in learning speed produced by the revaluation associability. It uniquely assumes, due to these processes being dissociable, that increasing the uncertainty of reinforcers in general should attenuate this sigmoidal shape, while not completely removing the latent inhibition effect. Furthermore, exposing the context by itself after CS exposure should likewise diminish the latent inhibition effect, as this would weaken the context-CS prediction during subsequent acquisition. In a similar manner, the model reproduces the effect observed by Leung et. al. whereby latent inhibition is enhanced by a preexposure of a CS compound. As this effect is explained in the model by the cue that undergoes reinforcement being predicted by the associative representation of the cue it retrieves, the implication is that presenting this retrieved cue in isolation before the acquisition phase should attenuate the observed effect. These preexposure related learning mechanisms also carry over to offering an account of the Hall-Pearce effect, by maintaining that the reinforcement of a CS, using a weak reinforcer, acts in many ways as a form of preexposure treatment from the point of view of the model, due to the lower intensity of the US in the first phase treatment. Therefore, the model’s explanation of this phenomena deviates from more traditional accounts like that of the Pearce-Hall model, which explains negative transfer as arising due to the CS fully predicting the US in the first phase, and thereby becoming less associable in later training. In contrast, the DDA model assumes that the negative transfer effect will persist even with few acquisition trials with the weak outcome, as the decay in the associability of the CS to the reinforcer is influenced more by the weak intensity of the US than by CS-US learning. This explanation for the Hall-Pearce effect additionally implies that conducting the second phase of the procedure in a novel context should lessen the observed effect. Lastly, for the effect of perceptual learning, differential CS-CS learning and variable attention shifts between intermixed and blocked presentations of CS preexposure trials allow the model to account for the effect being more pronounced with intermixed CS compounds. Specifically, weaker within-compound associations lead to differential rates of mediated conditioning in later stages of the procedure.

The DDA model’s dynamic asymptote of learning produces either excitatory or inhibitory learning depending on the causal connection between stimuli as tracked by the distance between the activations of the predicting and predicted stimulus. Through the interplay of this asymptote and its mechanism of associative retrieval the model accounts for mediated negative correlations and mediated conditioning. In the former, the claim is made that the poor correlation between the activity of the US representation and that of the CS representation interacts with the dynamic asymptote to induce the CS to form a slight inhibitory link towards the US through context-mediated revaluation. As this revaluation conditioned inhibition is dependent upon context retrieval of the US, the model predicts that strong context-US learning would strengthen the effect.

Mediated conditioning of a CS is produced in the DDA model through the combined effects of the aforesaid asymptote and associative retrieval by a second (previously paired) CS. Due to the asymmetry of the dynamic asymptote, learning between the associatively retrieved outcome of this CS and the US tends to be excitatory and proportional to the degree to which the CS representation is retrieved. Consequently, the DDA model assumes that the strength of mediated conditioning is proportional to the strength of within-compound associations, i.e., the length of CS-CS training. It is additionally predicted that the observed direction of learning in mediated learning will vary based on prior learning towards the outcome. For instance, though the asymptote of the retrieved CS in backward blocking and mediated conditioning is the same, the direction of learning in the DDA model follows an opposite direction for these phenomena due to the initial associative strength of the retrieved cue differing in the designs. Further, the extent of backward blocking, we claim, should be proportional to the amount of reinforced compound trials in the first phase of a retrospective revaluation treatment. This prediction supersedes the static learning rules between A1 and A2 stimuli formulated in the extensions of SOP – Dickinson and Burke (1996) and Holland (1983) – or the negative learning rate used in many models for mediated learning, which would each expect mediated learning to follow the same direction for the retrieved CS in both procedures. The model rather proposes that learning between an absent and present cue is dependent upon how strongly the absent cue is retrieved, as well as on the prior link the absent cue has towards the present cue. This account of mediated learning occurs completely within the confines of an associative learning rule, thus not relying on non-associative mechanisms such as a comparator or re-playing of past experiences.

Finally, while retaining an elemental nature, the DDA model can also reproduce a variety of non-linear discriminations without assuming that an animal has access to de novo configural information. It does so through a multi-factorial approach. The fully-connected network architecture of the model, which allows for shared elements among multiple stimuli, produces effects such as negative patterning and biconditional discriminations similarly to the Rescorla-Wagner model with common elements. Unlike the REM and Harris models, the DDA model does not propose that elements are differentially sampled or active depending on their context. Rather, shared elements exert their effect in the DDA model through their redundancy on compound trials (i.e., the same shared element cannot be sampled twice at the same time). The DDA model’s variable revaluation alpha can amplify discrimination learning through a unique decay process, which lowers the associability of cues presented with a persistently uncertain outcome. Silent learning and attentional processes, as mentioned before, produce unitization of shared and unique elements, as well as latent inhibition and a greater loss of associability of shared elements. In this latter aspect, it bears similarities to the CS-CS learning of the McLaren-Mackintosh and SLGK models. The model also explains the discrepancy between external inhibition and overshadowing without reference to additional representational structures such as the attentional buffer used in the Harris model. These shared elements and learning processes also furnish an account of non-linear discrimination effects, including negative patterning, biconditional discriminations, and perceptual learning using a completely elemental representation.

### New predictions of the DDA model

At this point, we are in a position to formulate some new predictions and to explain within the DDA framework experimental data that poses a challenge to existing models. The DDA model predicts the potentiation of blocking by compound conditioning in a different context. Compound conditioning in the new context would lead to mediated extinction of the context in which the blocking stimulus was trained, which would summate with the associative strength of the blocked cue when the response elicited by the latter is measured during test. Additionally, the predictor error would increase since the new context is not associated with the blocking stimulus, producing a prediction larger than the previous prediction, which would promote inhibitory learning for both compound stimuli. Similarly, the DDA model predicts that in a sensory preconditioning paradigm, conditioning in a novel context may foster retrospective revaluation. The model’s error-correction and revaluation alphas generate another interesting prediction, that preexposure in an excitatory or in an inhibitory context should potentiate latent inhibition. This effect would be asymmetric, with preexposure in an excitatory context producing and early delay in conditioning, whereas preexposing in an inhibitory context would generate a more prolonged retardation. The model also predicts that blocking would be attenuated in proportion to the perceptual similarity between the stimuli (Soto 2016; Soto, Gershman & Niv, 2014), a result which bears high relevance in the current debate about its alleged elusive nature (Maes, *et al.*, 2016). Finally, the model predicts that elongating the inter-trial interval would increase the speed of extinction and attenuate renewal effects (Urcelay, Wheeler & Miller, 2009). Such an effect arises in the DDA model due to increased ITI length deepening context extinction.

With regards to learning theory, the DDA model could contribute to the debate about the nature of human learning as an associative alternative to inferential and dual-process perspectives (for a review, Shanks, 2010). For instance, De Houwer *et al.*, (2002) suggested that blocking may arise as a result of reasoning, rather than from associative processes. They argued that providing the subjects with information about the maximal magnitude of the outcome would influence inference about the causal relationship between the blocked stimulus and the outcome. Consistently with this idea, they found that when the blocking stimulus signaled the maximal outcome magnitude, blocking was reduced, whereas when it signaled a submaximal intensity, blocking was enhanced. This result has been taken as inconsistent with associative theories. The DDA model could however account for these results by direct context strength summation: According to the model, if blocking training occurs in a previously trained highly excitatory context, which could parallel the provisioning of maximal information about the outcome, blocking would be diminished. On the contrary, if blocking training is given in an inhibitory context, a consequence of a discrepancy between the actual outcome and the expected outcome (submaximal), blocking would be deepened.

Offering a consistent account of these and other cue selection phenomena may prove decisive in unravelling the relationship between associative and cognitive processes of learning. Despite its apparent simplicity, we are still lacking a comprehensive associative theoretical analysis of the mechanisms underlying Pavlovian conditioning. By incorporating context modulation of learning via a predictor error and a unified mechanism to integrate attentional factors, and silent and reinforced learning into its associative structure, the DDA model offers a well-founded approach to stimulus selection that may play a crucial role in solving the credit assignment problem.

At a more theoretical level, the DDA model could be instrumental in scaling up associative accounts into high-order cognition. For instance, a case could be made that in the light of reinforcement learning models that establish the role of Pavlovian contingencies in goal-directed behavior (e.g., Balleine & Dickinson, 1998; Dayan & Berridge, 2014), a well-specified Pavlovian associative structure such as DDA’s could aid to understanding decisionmaking. Likewise, Pavlovian conditioning has also been integrated with traditional cognitive models such as drift-diffusion (e.g., Luzardo, Alonso, & Mondragón, 2017). However, a major limitation of associative theories arises from poor representational mechanisms in which events are treated as ex nihilo entities and for which the only semantics of the system relies on the strength of the associations. We claim that DDA’s associative architecture together with richer hierarchical representations (see Mondragón, Alonso, & Kokkola, 2017) might empower associationist paradigms to accommodate high-order cognitive functions.

**Table 4:** Experiments and predictions of associative learning models. R-W = Rescorla & Wagner (1972); P-H = Pearce & Hall (1980); M = Mackintosh (1975); H= Harris (2006); LP = Le Pelley (2004); Holland = Holland (1983); D-B= Dickinson & Burke (1996); SLGK = Kutlu & Schmajuk (2012); Pearce’s Configural = Pearce (1987); McLaren-Mackintosh = McLaren & Mackintosh (2000); Miller’s Comparator = Miller & Matzel (1988); Latent Causes = Gershman & Niv (2012); KTD = Gershman (2015); DDA = Double Error Dynamic Asymptote (this paper).

| Exp./Models | R-W | Attentional<br>(P-H, M, H, LP) | SOP Extensions<br>*Holland vs. D-B* |  | SLGK | Pearce's<br>Configural | McLaren-Mackintosh | Miller's<br>Comparator | Latent<br>Causes | KTD | DDA |
| --- | --- | --- | --- | --- | --- | --- | --- | --- | --- | --- | --- |
| Cue-competition | Y | Y but for M | Y |  | Y | Y | Y | Y | Y | Y | Y |
| Extinction related | N | N | partial |  | partial | N | partial | partial | partial | Y | Y |
| Temporal phenomena | N | N | partial |  | partial | N | partial | partial | N | partial | **partial** |
| Context Specificity of LI | N | P-H | Y |  | Y | N | Y | Y | unknown | unknown | Y |
| Compound latent inhibition | N | N | Y |  | N | N | unclear | N | unknown | Y | Y |
| Hall-Pearce Effect | N | P-H | N |  | Y | N | unclear | partial | Y | N | Y |
| Retrospective N. Correlation | N | M | unknown |  | unknown | N | Y | Y | unknown | N | Y |
| Perceptual Learning | N | H | unknown |  | N | N | Y | N | unknown | N | Y |
| SPC (Mediated Cond.) | N | N | Y | N | N | N | Y | partial | unknown | partial | Y |
| Mediated Extinction | N | N | Y | N | Y | N | Y | Y | unknown | N | Y |
| Unovershadowing | N | N | N | Y | N | N | Y | Y | unknown | unknown | Y |
| Backward Blocking | N | N | N | Y | Y | N | N | Y | unknown | unknown | Y |
| Backward SPC | N | N | Y | N | N | N | Y | N | unknown | N | Y |
| Negative Patterning | Y | N | unknown |  | Y | Y | Y | unknown | unknown | N | Y |
| Biconditional Discrimination | N | N | unknown |  | Y | Y | Y | unknown | unknown | N | Y |
| Ext. inhibition vs.overshadow. | Y | H | unknown |  | unknown | N | Y | unknown | Y | unknown | Y |

## Footnotes

1 To illustrate this idea mathematically, if an element *e*_1_ is active at time *t*, and a second element *e*_2_ has a sampling probability of 1/*x*, then the probability of excitatory learning taking place (within an acquisition trial) would be 1/*x* versus (*x*–1)/*x* for extinction. As such, only elements that are temporally highly correlated in terms of sampling probability would form robust associations. Therefore, learning would degenerate from a predictive relation to mere all or nothing correlation.

2 In Table 2, the US salience appears as *β*, following standard nomenclature in the literature.

3 In order to induce responding, the outcome should be a US.

## REFERENCES

Aguado, L., Symonds, M., & Hall, G. (1994). Interval between preexposure and test determines the magnitude of latent inhibition: Implications for an interference account. Animal Learning & Behavior, 22(2), 188–194.

Ahveninen, J., Kähkönen, S., Tiitinen, H., Pekkonen, E., Huttunen, J., Kaakkola, S., Ilmoniemi, R. J., & Jääskeläinen, I. P. (2000). Suppression of transient 40-Hz auditory response by haloperidol suggests modulation of human selective attention by dopamine D2 receptors. Neuroscience Letters, 292(1), 29–32.

Aitken, M. R., Larkin, M. J. & Dickinson, A. (2001). Re-examination of the role of within-compound associations in the retrospective revaluation of causal judgements. Quarterly Journal of Experimental Psychology, 54B, 27–51.

Alonso, E., Mondragón, E., & Fernández, A. (2012). A Java simulator of Rescorla and Wagner’s prediction error model and configural cue extensions. Computer Methods and Programs in Biomedicine, 108(1), 346–355.

Alonso, E., Sahota, P. & Mondragón, E. (2014). Computational Models of Classical Conditioning – A Qualitative Evaluation and Comparison. In B. Duval, J. van den Herik, S. Loiseau & J. Filipe (Eds.), Proceedings of the 6th International Conference on Agents and Artificial Intelligence (pp. 544–547). Setúbal, Portugal: SCITEPRESS.

Alonso, E., & Schmajuk, N. (2012). Special issue on computational models of classical conditioning guest editors’ introduction. Learning & Behavior, 40(3), 231–240.

Amundson, J. C., & Miller, R. R. (2008). CS-US temporal relations in blocking. Learning & Behavior, 36(2), 92–103.

Baker, A. G., Murphy, R. A., & Mehta, R. (2003). Learned irrelevance and retrospective correlation learning. The Quarterly Journal of Experimental Psychology: Section B, 56(1), 90–101.

Balkenius, C., & Morén, J. (1998). Computational models of classical conditioning: A comparative study. In R. Pfeifer, B. Blumberg, J.-A. Meyer & S. W. Wilson (Eds.), From Animals to Animats 5 (pp. 348–353). Cambridge, MA: MIT Press.

Balleine, B. W., & Dickinson, A. (1998). Goal-directed instrumental action: contingency and incentive learning and their cortical substrates. Neuropharmacology, 37(4-5), 407–419.

Blair, C. A. J., & Hall, G. (2003). Perceptual learning in flavor aversion: Evidence for learned changes in stimulus effectiveness. Journal of Experimental Psychology: Animal Behavior Processes, 29(1), 39.

Blaisdell, A. P., Gunther, L. M. & Miller, R. R. (1999). Recovery from blocking achieved by extinguishing the blocking CS. Animal Learning & Behavior, 27, 63–76.

Blough D. S. (1975). Steady state data and a quantitative model of operant generalization and discrimination. Journal of Experimental Psychology: Animal Behavior Processes, 1(1), 3–21.

Boddez, Y., Baeyens, F., Hermans, D. & Beckers, T. (2011). The hide-and-seek of retrospective revaluation: recovery from blocking is context dependent in human causal learning. Journal of Experimental Psychology: Animal Behavior Processes, 37, 230–240.

Bonardi, C., & Hall, G. (1996). Learned irrelevance: No more than the sum of CS and US preexposure effects? Journal of Experimental Psychology: Animal Behavior Processes, 22(2), 183.

Bouton M. E. (1993). Context, time and memory retrieval in the interference paradigms of Pavlovian learning. Psychological Bulletin, 114, 80–99.

Bouton M. E. (2004). Context and behavioral processes in extinction. Learning & Memory, 11(5), 485–94.

Brandon, S. E., Vogel, E. H., & Wagner, A. R. (2000). A componential view of configural cues in generalization and discrimination in Pavlovian conditioning. Behavioural Brain Research, 110(1), 67–72.

Brandon, S. E., Vogel, E. H., & Wagner, A. R. (2003). Stimulus representation in SOP: I. Theoretical rationalization and some implications. Behavioural Processes, 62(1-3), 5–25.

Brogden W. J. (1939). Sensory pre-conditioning. Journal of Experimental Psychology, 25(4), 323–332.

Bush, R. R., & Mosteller, F. (1955). Stochastic models for learning. New York, NJ: John Wiley & Sons, Inc.

Channell, S., & Hall, G. (1983). Contextual effects in latent inhibition with an appetitive conditioning procedure. Learning & Behavior, 11(1), 67–74.

Colwill, R. M., & Motzkin, D. K. (1994). Encoding of the unconditioned stimulus in Pavlovian conditioning. Learning & Behavior, 22(4), 384–394.

Corlett, P. R., Murray, G. K., Honey, G. D., Aitken, M. R. F., Shanks, D. R., Robbins, T. W., Bullmore, E. T., Dickinson, A., & Fletcher, P. C. (2007). Disrupted prediction-error signal in psychosis: evidence for an associative account of delusions. Brain, 130(9), 2387–2400.

Courville A. C., Daw, N. D., & Touretzky, D.S. (2006). Bayesian theories of conditioning in a changing world. Trends in Cognitive Sciences, 10, 294–300.

Dayan, P., & Berridge, K. C. (2014). Model-based and model-free Pavlovian reward learning: revaluation, revision, and revelation. Cognitive, Affective, & Behavioral Neuroscience, 14(2), 473–492.

De Houwer J., Beckers, T., & Glautier, S. (2002). Outcome and cue properties modulate blocking. Quarterly Journal of Experimental Psychology, 55A:965–85.

Dickinson A. (1996). Within compound Associations Mediate the Retrospective Revaluation of Causality Judgements. The Quarterly Journal of Experimental Psychology Section B, 49(1), 60–80.

Dickinson, A., & Burke, J. (1996). Within-compound associations mediate the retrospective revaluation of causality judgements. Quarterly Journal of Experimental Psychology, 37B, 397–416.

Dickinson, A., Hall, G., & Mackintosh, N. J. (1976). Surprise and the attenuation of blocking. Journal of Experimental Psychology: Animal Behavior Processes, 2(4), 313–322.

Dwyer D. M. (1999). Retrospective revaluation or mediated conditioning? The effect of different reinforcers. The Quarterly Journal of Experimental Psychology, 52(4), 289–306.

Estes W. K. (1950). Toward a statistical theory of learning. Psychological Review, 57(2), 94–107.

Gallistel, C. R., Fairhurst, S., & Balsam, P. (2004). The learning curve: implications of a quantitative analysis. Proceedings of the National Academy of Sciences of the United States of America, 101(36), 13124–31.

Gershman S. J. (2015). A Unifying Probabilistic View of Associative Learning. PLoS Computational Biology, 11(11): e1004567.

Gershman, S. J., Blei, D. M., & Niv, Y. (2010). Context, Learning, and Extinction. Psychological Review, 117(1), 197–209.

Gershman, S. J., & Niv, Y. (2012). Exploring a latent cause theory of classical conditioning. Learning & Behavior, 40(3), 255–268.

Ghirlanda S. (2015). On elemental and configural models of associative learning. Journal of Mathematical Psychology, 64, 8–16.

Ghirlanda, S., & Ibadullayev, I. (2015). Solution of the comparator theory of associative learning. Psychological Review, 122(2), 242.

Gibbon J. (1977). Scalar expectancy theory and Weber’s law in animal timing. Psychological Review, 84, 279–325.

Ginsburg, S., & Jablonka, E. (2010). The evolution of associative learning: A factor in the Cambrian explosion. Journal of Theoretical Biology, 266(1), 11–20.

Glautier S. (2013). Revisiting the learning curve (once again). Frontiers in Psychology, 4, 982.

Glautier, S., Redhead, E., Thorwart, A., & Lachnit, H. (2010). Reduced Summation with Common Features in Causal Judgments. Experimental Psychology, 57(4), 252–259.

Gomez, M., De Castro, E., Guarin, E., Sasakura, H., Kuhara, A., Mori, I., Bartfai, T., Bargmann, C. I., & Nef, P. (2001). Ca2+ Signaling via the Neuronal Calcium Sensor-1 Regulates Associative Learning and Memory in C. elegans. Neuron, 30(1), 241–248.

Hall G. (1991). Perceptual and associative learning Oxford, UK: Clarendon Press-Oxford University Press.

Hall G. (2002). Associative structures in Pavlovian and instrumental conditioning. In H. Pashler, S. Yantis, D. Medin, R. Gallistel & J. Wixted (Eds.), Stevens’ Handbook of Experimental Psychology, Volume 3 (pp. 1–45). Hoboken, NJ: John Wiley and Sons.

Hall, G., & Honey, R. C. (1989). Contextual effects in conditioning, latent inhibition, and habituation: associative and retrieval functions of contextual cues. Journal of Experimental Psychology: Animal Behavior Processes, 15(3), 232.

Hall, G., & Pearce, J. M. (1979). Latent inhibition of a CS during CS-US pairings. Journal of Experimental Psychology. Animal Behavior Processes, 5(1), 31–42.

Hall G. & Rodriguez G. (2010). Associative and nonassociative processes in latent inhibition: An elaboration of the Pearce-Hall model. In R.E. Lobow & I. Weiner (Eds.), Latent inhibition: Cognition, neuroscience, and applications to schizophrenia (pp. 114–136). Cambridge, England: Cambridge University Press.

Harris J. A. (2006). Elemental Representations of Stimuli in Associative Learning. Psychological Review, 113(3), 584–605.

Harris, J., Livesey, E. (2010). An attention-modulated associative network. Learning and Behavior, 38(1), 1–26.

Haselgrove, M., & Hogarth, L. (2011). Clinical Applications of Learning Theory. Hove, England: Psychology Press.

Holland P. C. (1983). Representation-mediated overshadowing and potentiation of conditioned aversions. Journal of Experimental Psychology: Animal Behavior Processes, 9(1), 1–13.

Holland P. C. (1984). Unblocking in Pavlovian appetitive conditioning. Journal of Experimental Psychology: Animal Behavior Processes 10, 476–497.

Holland P. C. (1985). The nature of conditioned inhibition in serial and simultaneous feature negative discriminations. In R. R. Miller & N. E. Spear (Eds.), Information processing in animals: Conditioned inhibition (pp. 267–297). Hillsdale, NJ: Erlbaum.

Holland, P. C., & Forbes, D. T. (1982). Representation-mediated extinction of conditioned flavor aversions. Learning and Motivation, 13(4), 454–471.

Holland, P. C., & Schiffino, F. L. (2016). Mini-review: Prediction errors, attention and associative learning. Neurobiology of Learning and Memory, 131, 207–215.

Hull L. C. (1943). Principles of behavior: an introduction to behavior theory. Oxford, England: Appleton-Century.

Jie H. L. (2008). Neuroimaging of associative learning (master’s thesis). National University of Singapore, Singapore.

Kamin L. J. (1968). “Attention-like” processes in classical conditioning. In M.R. Jones (Ed.), Miami Symposium on the Prediction of Behavior, 1967: Aversive Stimulation (pp. 9–31). Coral Gables, FL: University of Miami Press.

Kamin L. J. (1969). Selective association and conditioning. In N.J. Mackintosh & W.K. Honig (Eds.), Proceedings of the Symposium on Fundamental issues in Associative Learning (pp. 42–64). Halifax, Canada: Dalhousie University Press.

Kehoe, E. J., Schreurs, B. G., & Graham, P. (1987). Temporal primacy overrides prior training in serial compound conditioning of the rabbit’s nictitating membrane response. Animal Learning & Behavior, 15(4), 455–464.

Kobayashi, K., & Poo, M. (2004). Spike Train Timing-Dependent Associative Modification of Hippocampal CA3 Recurrent Synapses by Mossy Fibers. Neuron, 41(3), 445–454.

Kohler, E. A., & Ayres, J. J. B. (1979). The Kamin blocking effect with variable-duration CSs. Animal Learning & Behavior, 7(3), 347–350.

Konorski J. (1948). Conditioned reflexes and neuron organization. Cambridge, MA: Cambridge University Press.

Kurschke J. K. (2008). Bayesian approaches to associative learning: From passive to active learning. Learning & Behavior, 36(3), 210–226.

Kruschke J. K. (2011). Models of attentional learning. In: E. M. Pothos & A. J. Wills (Eds.), Formal Approaches in Categorization (pp. 120–152). Cambridge, England: Cambridge University Press.

Kutlu, M. G., & Schmajuk, N. A. (2012). Solving Pavlov’s puzzle: Attentional, associative, and flexible configural mechanisms in classical conditioning. Learning & Behavior, 40(3), 269–291.

Lachnit, H., Schultheis, H., König, S., Üngör, M., & Melchers, K. (2008). Comparing elemental and configural associative theories in human causal learning: A case for attention. Journal of Experimental Psychology: Animal Behavior Processes, 34(2), 303–313.

Le Pelley, M. E. (2004). The role of associative history in models of associative learning: A selective review and a hybrid model. The Quarterly Journal Of Experimental Psychology, 57B(3), 193–243.

Le Pelley, M. E., Haselgrove, M., & Esber, G. R. (2012). Modeling attention in associative learning: Two processes or one? Learning & Behavior, 40(3), 292–304.

Le Pelley, M. E., & McLaren, I. P. L. (2001). Retrospective revaluation in humans: Learning or memory? The Quarterly Journal of Experimental Psychology: Section B, 54(4), 311–352.

Leung, H. T., Killcross, A. S., & Westbrook, R. F. (2011). Additional exposures to a compound of two preexposed stimuli deepen latent inhibition. Journal of Experimental Psychology: Animal Behavior Processes, 37(4), 394.

Liljeholm M. & Balleine B. W. (2006). Stimulus salience and retrospective revaluation. Journal of Experimental Psychology: Animal Behavior Processes, 29, 97–106.

Liljeholm M. & Balleine B. W. (2009). Mediated conditioning versus retrospective revaluation in humans: The influence of physical and functional similarity of cues. Quarterly Journal of Experimental Psychology, 62(3), 470–482.

Lober, K., & Lachnit, H. (2002). Configural learning in human Pavlovian conditioning: acquisition of a biconditional discrimination. Biological Psychology, 59(2), 163–168.

Lubow R. E. (1965). Effects of frequency of nonreinforced preexposure of the CS. Journal of Comparative and Physiological Psychology, 60(3), 454–457.

Ludvig, E. A., Bellemare, M. G., & Pearson, K. G. (2011). A primer on reinforcement learning in the brain: Psychological, computational, and neural perspectives. In E. Alonso & E. Mondragón, Computational Neuroscience for Advancing Artificial Intelligence: Models, Methods and Applications (pp. 111–144). Hershey, PA: IGI Global.

Ludvig, E. A., & Koop, A. (2008). Learning to Generalize through Predictive Representations: A Computational Model of Mediated Conditioning. In M. Asada, J. C. T. Hallam, J.-A. Meyer & J. Tani, From Animals to Animats 10 (pp. 342–351). Berlin, Heidelberg: Springer Berlin Heidelberg.

Ludvig, E. A., Mirian, M. S., Kehoe, E. J., & Sutton, R. S. (2017). Associative Learning from Replayed Experience. bioRxivPreprint. doi: https://doi.org/10.1101/100800.

Ludvig, E. A., Sutton, R. S., Verbeek, E., & Kehoe, E. J. (2009). A computational model of hippocampal function in trace conditioning. In D. Koller, D. Schuurmans, Y. Bengio & L. Bottou (Eds.), Advances in Neural Information Processing Systems 21 (pp. 993–1000). Red Hook, NY: Curran Associates, Inc.

Luzardo, A., Alonso, E. & Mondragón, E. (2017). A Rescorla-Wagner drift-diffusion model of conditioning and timing. PLOS Computational Biology, 13 (11): e1005796.

Mackintosh N. J. (1973). Stimulus selection: Learning to ignore stimuli that predict no change in reinforcement. In R. A. Hinde & J. Stevenson-Hinde, Constraints on learning: Limitations and predispositions (pp. 75–96). Oxford, England: Academic Press.

Mackintosh N. J. (1975). A theory of attention: Variations in the associability of stimuli with reinforcement. Psychological Review, 82(4), 276–298.

Mackintosh, N. J., Kaye, H., & Bennett, C. H. (1991). Perceptual learning in flavour aversion conditioning. The Quarterly Journal of Experimental Psychology, 43(3), 297–322.

Maes, E., Boddez, Y., Alfei, J. M., Krypotos, A. M., D’Hooge, R., De Houwer, J., & Beckers, T. (2016). The elusive nature of the blocking effect: 15 failures to replicate. Journal of Experimental Psychology: General, 145(9), e49–e71.

Marschner, A., Kalisch, R., Vervliet, B., Vansteenwegen, D., & Büchel, C. (2011). Neural correlates of human associative learning. Tsinghua Science & Technology, 16(2), 140–144.

Matzel, L. D., Schachtman, T. R. & Miller, R. R. (1985). Recovery of an overshadowed association achieved by extinction of the overshadowing stimulus. Learning and Motivation, 16, 398–412.

McLaren I. P. L. (1993). APECS: A solution to the sequential learning problem. In M. Ringle (Ed.), Proceedings of the Fifteenth Annual Convention of the Cognitive Science Society (pp. 717–722). Hillsdale, NJ: Lawrence Erlbaum.

McLaren, I. P. L., & Mackintosh, N. J. (2000). An elemental model of associative learning: I. Latent inhibition and perceptual learning. Animal Learning & Behavior, 28(3), 211–246.

Miller, R. R., Barnet, R. C., & Grahame, N. J. (1995). Assessment of the Rescorla-Wagner model. Psychological Bulletin, 117(3), 363–386.

Miller, R. R., & Matzel, L. D. (1988). The Comparator Hypothesis: A Response Rule for The Expression of Associations. Psychology of Learning and Motivation, 22, 51–92.

Miller, R. R., & Witnauer, J. E. (2016). Retrospective revaluation: The phenomenon and its theoretical implications. Behavioural Processes, 123, 15–25.

Mondragón, E., Alonso, E., Fernández, A., & Gray, J. (2013). An extension of the Rescorla and Wagner Simulator for context conditioning. Computer Methods and Programs in Biomedicine, 110(2), 226–230.

Mondragón, E., Alonso, E. & Kokkola, N. (2017). Associative learning should go deep. Trends in Cognitive Sciences, 21(11), 822–825.

Mondragón, E., Gray, J., & Alonso, E. (2013). A Complete Serial Compound Temporal Difference Simulator for Compound stimuli, Configural cues and Context representation. Neuroinformatics, 11(2), 259–261.

Mondragón, E., Gray, J., Alonso, E., Bonardi, C., & Jennings, D. J. (2014). SSCC TD: A Serial and Simultaneous Configural-Cue Compound Stimuli Representation for Temporal Difference Learning. PLoS ONE, 9(7), e102469.

Mondragón, E., & Hall, G. (2002). Analysis of the perceptual learning effect in flavour aversion learning: Evidence for stimulus differentiation. The Quarterly Journal of Experimental Psychology: Section B, 55(2), 153–169.

Mondragón, E., & Murphy, R. A. (2010). Perceptual learning in an appetitive Pavlovian procedure: Analysis of the effectiveness of the common element. Behavioural Processes, 53(3), 247–256.

Montague, P. R., Dayan, P., & Sejnowski, T. J. (1996). A framework for mesencephalic dopamine systems based on predictive Hebbian learning. Journal of Neuroscience, 16(5), 1936–47.

Moore, J. W., Choi, J.-S., & Brunzell, D. H. (1998). Predictive timing under temporal uncertainty: The time derivative model of the conditioned response. In D. A. Rosenbaum & C. E. Collyer (Eds.), Timing of behavior: Neural, psychological, and computational perspectives (pp. 3–34). Cambridge, MA: The MIT Press.

Murphy, R.A., Mondragón, E. & Murphy, V. A. (2008). Rule learning by rats. Science, 319(5871), 1849–1851.

Navarro, D. J., Lee, M. D., Dry, M. J., & Schultz, B. (2008). Extending and testing the Bayesian theory of generalization. In Proceedings of the 30th Annual Conference of the Cognitive Science Society (pp. 1746–1751). Austin, TX: Cognitive Science Society.

Nieoullon A. (2002). Dopamine and the regulation of cognition and attention. Progress in Neurobiology, 67(1), 53–83.

Niv Y. (2009). Reinforcement learning in the brain. Journal of Mathematical Psychology, 53(3), 139–154.

Niv, Y., Edlund, J. A., Dayan, P., & O’Doherty, J. P. (2012). Neural Prediction Errors Reveal a Risk-Sensitive Reinforcement-Learning Process in the Human Brain. Journal of Neuroscience, 32(2).

Panayi, M. C., & Killcross, S. (2014). Orbitofrontal cortex inactivation impairs between- but not within-session Pavlovian extinction: An associative analysis. Neurobiology of Learning and Memory, 108, 78–87.

Pavlov I. P. (1927). Conditioned reflexes: an investigation of the physiological activity of the cerebral cortex. Oxford, England: Oxford University Press.

Pearce J. M. (1987). A model for stimulus generalization in Pavlovian conditioning. Psychological Review, 94(1), 61–73.

Pearce, J. M., & Bouton, M. E. (2001). Theories of associative learning in animals. Annual Review of Psychology, 52, 111–39.

Pearce, J. M., George, D. N., & Aydin, A. (2002). Summation: Further assessment of a configural theory. The Quarterly Journal of Experimental Psychology: Section B, 55(1), 61–73.

Pearce, J. M., & Hall, G. (1980). A model for Pavlovian learning: Variations in the effectiveness of conditioned but not of unconditioned stimuli. Psychological Review, 87(6), 532–552.

Pearce, J. M., & Redhead, E. S. (1993). The influence of an irrelevant stimulus on two discriminations. Journal of Experimental Psychology: Animal Behavior Processes, 19(2), 180.

Pearce, J. M., & Wilson, P. N. (1991). Failure of excitatory conditioning to extinguish the influence of a conditioned inhibitor. Journal of Experimental Psychology: Animal Behavior Processes, 17(4), 519–529.

Razran G. H. S. (1939). Studies in configural conditioning: I. Historical and preliminary experimentation. The Journal of General Psychology, 21, 307–330.

Rescorla R. A. (1970). Reduction in the effectiveness of reinforcement after prior excitatory conditioning. Learning and Motivation, 1(4), 372–381.

Rescorla R. A. (1971a). Variation in the effectiveness of reinforcement and nonreinforcement following prior inhibitory conditioning. Learning and Motivation, 2(2), 113–123.

Rescorla R. A. (1971b). Summation and retardation tests of latent inhibition. Journal of Comparative and Physiological Psychology, 75(1), 77–81.

Rescorla R. A. (1972). “Configural” conditioning in discrete-trial bar pressing. Journal of Comparative and Physiological Psychology, 79(2), 307–317.

Rescorla R. A. (1985). Conditioned inhibition and facilitation. In R. R. Miller & N. E. Spear (Eds.), Information processing in animals: Conditioned inhibition (pp. 299–326). Hillsdale, NJ: Erlbaum

Rescorla R. A. (2004). Superconditioning from a reduced reinforcer. Quarterly Journal of Experimental Psychology Section B, 57(2), 133–152.

Rescorla, R. A., Grau, J. W., & Durlach, P. J. (1985). Analysis of the unique cue in configural discriminations. Journal of Experimental Psychology: Animal Behavior Processes, 11(3), 356–366.

Rescorla, R. A., & Wagner, A. R. (1972). A theory of Pavlovian conditioning: Variations in the effectiveness of reinforcement and nonreinforcement. In A. H. Black & W. F. Prokasy (Eds.). Classical Conditioning II: Current Research and Theory (pp. 64–99). New York, NJ: Appleton Century Crofts.

Roesch, M. R., Esber, G. R., Li, J., Daw, N. D., & Schoenbaum, G. (2012). Surprise! Neural correlates of Pearce-Hall and Rescorla-Wagner coexist within the brain. European Journal of Neuroscience, 35(7), 1190–1200.

Rosas, J. M., & Bouton, M. E. (1998). Context change and retention interval can have additive, rather than interactive, effects after taste aversion extinction. Psychonomic Bulletin & Review, 5(1), 79–83.

Saavedra M. A. (1975). Pavlovian compound conditioning in the rabbit. Learning and Motivation, 6(3), 314–326.

Schachtman, T. R., & Reilly, S. (2011). Associative Learning and Conditioning Theory: Human and Non-Human Applications. Oxford, England: Oxford University Press.

Schultheis, H., Thorwart, A., & Lachnit, H. (2008). Rapid-REM: a MATLAB simulator of the replaced-elements model. Behavior Research Methods, 40(2), 435–41.

Schultz W. (2004). Neural coding of basic reward terms of animal learning theory, game theory, microeconomics and behavioural ecology. Current Opinion in Neurobiology 14(2), 139–47.

Schultz W. (2006). Behavioral theories and the neurophysiology of reward. Annual Review of Psychology, 57, 87–115.

Schultz W. (2010). Dopamine signals for reward value and risk: basic and recent data. Behavioral and Brain Functions 6:24.

Schultz, W., Dayan, P., & Montague, P. R. (1997). A neural substrate of prediction and reward. Science 275(14), 1593–9.

Shanks D. R. (1995). The Psychology of Associative Learning. Cambridge, England: Cambridge University Press.

Shanks D. R. (2010). Learning: From Association to Cognition. Annual Review of Psychology, 61, 273–301.

Soto F. A. (in press). Contemporary associative learning theory predicts failures to obtain blocking. Comment on Maes et al. (2016). Journal of Experimental Psychology: General.

Soto, F.A., Gershman, S.J. & Niv, Y. (2014). Explaining compound generalization in associative and causal learning through rational principles of dimensional generalization. Psychological Review, 121(3):526–558.

Spence K. W. (1936). The nature of discrimination learning in animals. Psychological Review, 43(5), 427–449.

Spence K. W. (1937). The differential response in animals to stimuli varying within a single dimension. Psychological Review, 44(5), 430–444.

Sutton, R. S., & Barto, A. G. (1981). Toward a modern theory of adaptive networks: Expectation and prediction. Psychological Review, 88(2), 135–70.

Sutton, R. S., & Barto, A. G. (1987). A temporal-difference model of classical conditioning. In J. D. Moore & J. F. Lehman (Eds.), Proceedings of the Ninth Annual Conference of the Cognitive Science Society (pp. 355–378). Mahwah, NJ: Erlbaum.

Swartzentruber, D., & Bouton, M. E. (1986). Contextual control of negative transfer produced by prior CS-US pairings. Learning and Motivation, 17(4), 366–385.

Symonds, M., & Hall, G. (1995). Perceptual learning in flavor aversion conditioning: Roles of stimulus comparison and latent inhibition of common stimulus elements. Learning and Motivation, 26(2), 203–219.

Tenenbaum, J., & Griffiths, T. (2001). Generalization, similarity, and Bayesian inference. Behavioral and Brain Sciences, 24(4), 629–640.

Turkkan J. S. (1989). Classical conditioning beyond the reflex: An uneasy rebirth. Behavioral and Brain Sciences, 12(1), 161–179.

Urcelay, G.P., Wheeler, D.S. & Miller, R.R. (2009). Spacing extinction trials alleviates renewal and spontaneous recovery. Learning & Behavior, 37(1), 60–73.

Urushihara, K., & Miller, R. R. (2010). Backward blocking in first-order conditioning. Journal of Experimental Psychology. Animal Behavior Processes, 36(2), 281–95.

Wagner A.R. (1981). SOP: A model of automatic memory processing in animal behavior. In N.E. Spear & R.R. Miller (Eds.), Information processing in animals: Memory mechanisms (pp. 5–47). Hillsdale, NJ: Erlbaum.

Wagner A. R. (2003). Context-sensitive elemental theory. The Quarterly Journal of Experimental Psychology: Section B, 56(1), 7–29.

Wagner A. R. (2008). Evolution of an elemental theory of Pavlovian conditioning. Learning & Behavior, 36(3), 253–265.

Wagner, A. R., & Brandon, S. E., (2001) A componential theory of Pavlovian Conditioning. In R.R. Mowrer and S.B. Klien (Eds.), Handbook of Contemporary Learning Theories (pp. 23–64). Mahwah, NJ: Erlbaum.

Wagner, A. R., & Rescorla, R. A. (1972). Inhibition in Pavlovian conditioning: Application of a theory. In R.A. Boakes & M.S. Halliday (Eds.), Inhibition and learning (pp. 301–336). London, England: Academic Press.

Ward-Robinson, J., & Hall, G. (1996). Backward sensory preconditioning. Journal of Experimental Psychology: Animal Behavior Processes, 22(4), 395.

Ward-Robinson, J., & Hall, G. (1998). Backward Sensory Preconditioning When Reinforcement is Delayed. The Quarterly Journal Of Experimental Psychology, 51B(4), 349–362.

Westbrook, R. F., & Bouton, M. E. (2010). Latent inhibition and extinction: Their signature phenomena and the role of prediction error. In R. E. Lubow & I. Weiner (Eds.), Latent inhibition: Cognition, neuroscience, and applications to schizophrenia (pp. 23–39). New York, NY: Cambridge University Press.

Westbrook R, & Jones M. L. & Bailey G. K. & Harris J. A. (2000). Contextual control over conditioned responding in a latent inhibition paradigm. Journal of experimental psychology. Animal Behavior Processes 26, 157–73.

Whitlow, J. W., & Wagner, A. R. (1972). Negative patterning in classical conditioning: Summation of response tendencies to isolable and configurai components. Psychonomic Science, 27(5), 299–301.

Wilson, P. N., & Pearce, J. M. (1992). A configural analysis for feature-negative discrimination learning. Journal of Experimental Psychology: Animal Behavior Processes, 18(3), 265.

Zeithamova, D., Dominick, A. L., & Preston, A. R. (2012). Hippocampal and Ventral Medial Prefrontal Activation during Retrieval-Mediated Learning Supports Novel Inference. Neuron, 75(1), 168–179.

